# Ablative radiotherapy improves survival in autochthonous cancer mouse models

**DOI:** 10.1101/2022.07.01.498353

**Authors:** Daniel R. Schmidt, Iva Monique T Gramatikov, Allison Sheen, Christopher L. Williams, Martina Hurwitz, Laura E. Dodge, Edward Holupka, W S Kiger, Milton R. Cornwall-Brady, Wei Huang, Howard H. Mak, Kathy Cormier, Charlene Condon, K Dane Wittrup, Ömer H. Yilmaz, Mary Ann Stevenson, Julian D. Down, Scott R. Floyd, Jatin Roper, Matthew G. Vander Heiden

## Abstract

Genetically engineered mouse models (GEMMs) of cancer are powerful tools to study mechanisms of disease progression and therapy response, yet little is known about how these models respond to multimodality therapy used for curative intent in patients. Radiation therapy (RT) is frequently used to treat localized cancers with curative intent, delay progression of oligometastases, and palliate symptoms of metastatic disease. RT can be combined with surgery, systemic therapy, and/or immunotherapy for definitive treatment of solid tumors. Here we report the development, testing, and validation of a murine platform that faithfully emulates human stereotactic ablative radiotherapy (SABR). We demonstrate that SABR regimens used in clinical practice can be effectively delivered in mouse models and establish the intestinal tract as the dose limiting organ for abdominopelvic radiation with notable, but transient, lymphodepletion. SABR alters tumor stroma and immune environment, improves survival in GEMMs of primary prostate and colorectal cancer, and synergizes with androgen deprivation in prostate cancer. While SABR is capable of fully ablating xenografts it is unable to completely eradicate disease in GEMMs. These data show that some GEMMs are resistant to curative therapy suggesting the existence of treatment-resistant persister cells in these models.

## INTRODUCTION

Genetically engineered mouse models (GEMMs) have enabled many aspects of tumor biology to be studied that are not possible in other model systems. By combining gene editing techniques with technologies that allow temporal and spatial control of genetic alterations, all stages of oncogenic transformation and tumor progression can be studied for many cell lineages in an endogenous tissue context (1, 2). In addition to providing a window into the genetic mechanisms leading to cancer development, the ability to probe autochthonous tumors in their native microenvironment enables study of how immune cells, stroma, and local nutrients contribute to tumor progression and treatment response (3, 4).

Because many GEMMs progress from preinvasive lesions to invasive local disease and ultimately metastasis, these models have been extensively used to study cancer progression; however, they are also useful for studying mechanisms of treatment resistance and disease recurrence (5). Although GEMMs have been used to evaluate treatment efficacy (6–11), they have not yet been extensively used to evaluate response to potentially curative therapies or to examine mechanisms of recurrence post treatment. Most cancer patients receive multimodality therapy and understanding how to best sequence existing and novel therapies is an area where GEMMs could impact how clinical trials are designed. Another advantage of GEMMs is the ability to measure desired anti-cancer effects and unwanted side effects on normal tissues, both of which are critical to understanding how to best treat patients.

It is estimated that approximately 50% of cancer patients receive radiation therapy (RT) as part of their management, and about half of those patients receive RT with curative intent (12). For cancers that have not metastasized, RT is often given before or after surgery to reduce the risk of local recurrence. RT is also used as the primary local treatment and can be curative as monotherapy for some cancers. For more advanced disease treated with curative intent, RT is often combined with chemotherapy and in some cases surgery, and the combination of RT and immunotherapy for metastatic disease is currently being evaluated in clinical trials.

Historically, curative radiation therapy regimens consisted of 25 to 40 fractions of 1.8 to 2 Gy per fraction. A cumulative dose of 45 to 50 Gy is used for microscopic disease and 60 to 80 Gy for more bulky disease. Due to the relatively large size of early radiation fields, small fraction sizes were needed to allow normal tissues within the radiation field to recover between fractions and minimize toxicity. Technical advances in imaging and radiotherapy delivery have allowed larger doses per fraction (5 to 25 Gy) to be delivered to the tumor with a smaller margin while maintaining a low dose to surrounding normal tissues. Initially developed as a way to treat benign brain lesions, stereotactic radiotherapy involves precise positioning of a target within a conformal radiation field combined with target immobilization to ensure that it remains within the radiation field during treatment. This approach can deliver high doses of radiotherapy in a small number of fractions and has subsequently been adapted to treat malignant tumors. For historical reasons treatment of brain tumors with a single fraction of radiation is referred to as stereotactic radiosurgery, while treatment of extracranial tumors, typically with 3 to 5 fractions, is referred to as stereotactic ablative body radiotherapy (SABR) or stereotactic body radiation therapy (SBRT) (13).

SABR is now well established in the clinic and in some cases provides an alternative to surgery with comparable cure rates for early-stage tumors. SABR is also increasingly used in the oligometastatic setting to achieve long disease-free intervals. Treatment failure resulting in persistent or recurrent disease after RT has been attributed to various factors including intrinsic cancer cell radio-resistance and extrinsic factors such as hypoxia (14, 15). Cancer cells that are able to repair sub-lethal DNA damage can resume replication and repopulate a tumor (16), but the mechanisms underlying these events are incompletely understood.

Multiple studies have shown that stereotactic RT delivered in a single fraction or sub-ablative multi-fraction regimens is well tolerated in mice and can delay tumor growth in GEMMs (17–19). While the regimens used in these studies can be useful to examine some research questions, they are not representative of curative SABR regimens used in the clinic. A few studies have examined clinically relevant RT regimens in GEMMs (20, 21); however, what dose of radiation is tolerable and effective in mice has not been established and is a barrier to conducting pre-clinical work involving RT in mice. Moreover, whether radiation at definitive doses with or without adjuvants is able to cure tumors in GEMMs is not known.

In this study we describe the development of a novel stereotactic radiotherapy platform used to determine the maximum tolerated SABR dose that can be delivered to the lower abdomen and pelvis in mice. We demonstrate that complete responses are achievable in flank xenografts. In contrast, SABR is able to reduce tumor growth and improve survival in autochthonous prostate and colorectal tumor models, but does not achieve complete pathologic responses. These data demonstrate that GEMMs can be used to study SABR-induced changes in the tumor microenvironment, and suggest that radioresistant persister cells are present in these tumors.

## RESULTS

### Design of a stereotactic radiation delivery platform

A number of small animal radiation delivery systems offer image-guidance to deliver focal radiotherapy (22, 23); however, these platforms are costly and impractical for multi-fraction regimens and large treatment cohorts. Most platforms also require the animal to be anesthetized during imaging and treatment, which poses minimal risk for brief single fraction treatments, but is disadvantageous for the multi-fraction, high-dose regimens used in patients (24, 25). Since autochthonous tumors in mouse models generally develop in a defined anatomic location with characteristic growth rates, we reasoned that an appropriately sized radiation field combined with a method for immobilization for non-anesthetized animals would be sufficient to accurately deliver SABR. Such a platform should meet criteria mandated for clinical SABR, including high dose per fraction, steep dose gradient, stereotactic localization of the target, and method to limit or compensate for target movement during treatment (26). To this end we designed a series of modular restrainers for laboratory mice to allow accurate stereotactic positioning within the treatment field, minimize target movement during treatment, and accommodate male and female mice of various sizes (Fig. 1A, Supplementary Fig. 1A, Supplementary File 1-7). We found that the restrainers enabled consistent anatomic alignment in the treatment field (Fig. 1B), minimal intrafraction movement in a conscious animal (Fig. 1C, Supplementary Video 1), and reproducible positioning of male and female mice that span a wide range of body weights (Supplementary Fig. 1B). Importantly, animals tolerated immobilization in the restrainer for up to 30 minutes without overt signs of distress or changes in body weight. An accompanying set of lead shields and circular collimators with apertures ranging from 0.5 to 6 cm were designed to allow focal radiation of the immobilized target via a radiation source located above and/or below the animal (Fig. 1D, Supplementary File 8, 9).

**Figure 1.**
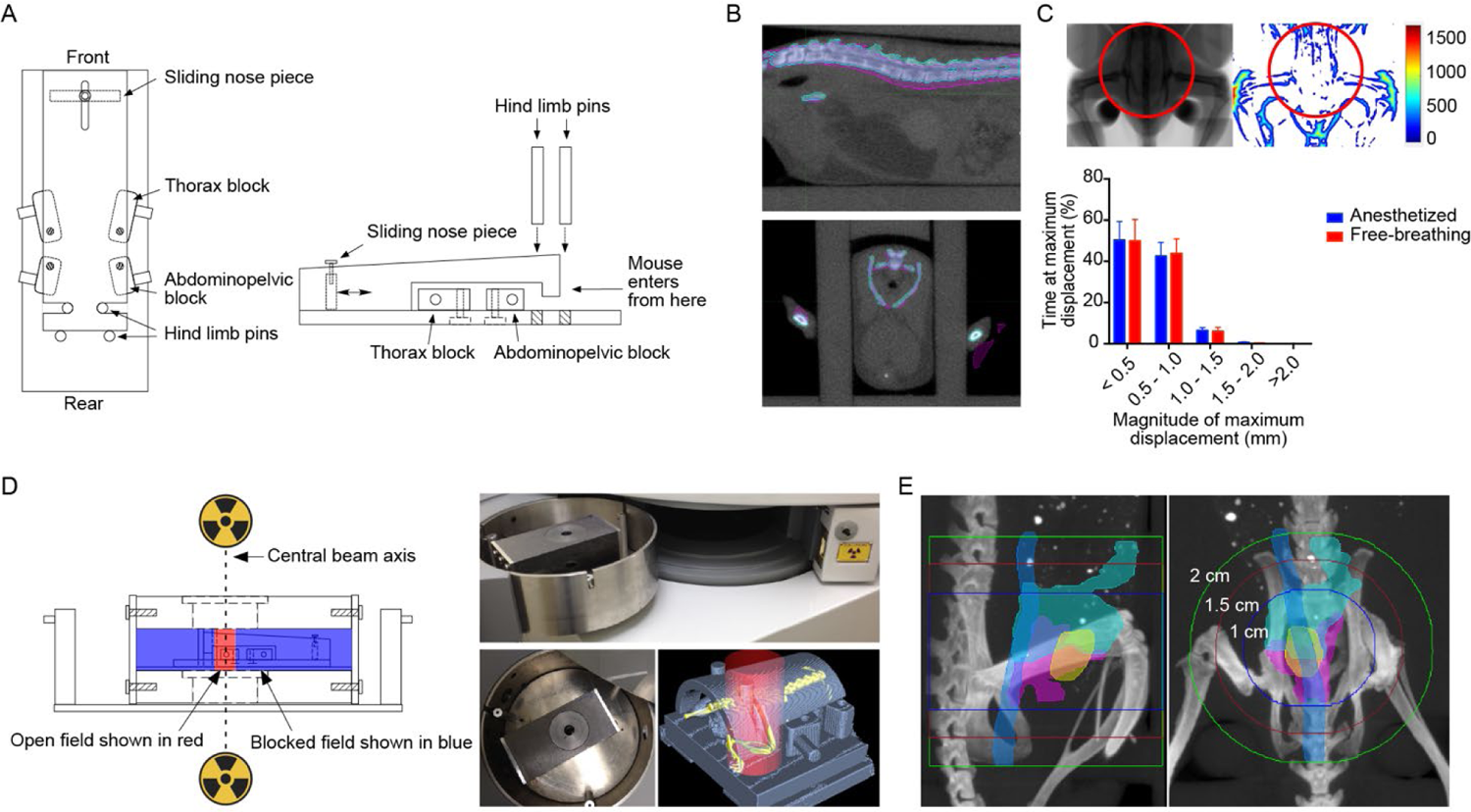
Developing a stereotactic radiotherapy platform. (**A**) Schematic of restrainer (top and side view) for delivery of stereotactic pelvic radiotherapy. (**B**) Serial imaging demonstrates negligible anatomic displacement between fractions. The pink and cyan colors indicate the bony anatomy of a mouse imaged on two separate days in the restrainer, with overlap demonstrated for the spine on sagittal view (upper panel) and pelvis on axial view (lower panel). (**C**) Intrafraction motion assessed by fluoroscopic imaging. The representative anatomic heatmap on the right shows the maximum displacement (in µm) of a free-breathing mouse during 3 minutes of continuous imaging. The histogram shows the percent of time that a maximum displacement occurs in the pelvic radiation field (area within the red circle). Data are plotted as mean with standard deviation (n=7 per group). (**D**) Schematic and photographs of lead shields with interchangeable collimators (2 cm aperture shown) used to focus radiation sources located above and below the animal. Example of a focal radiation field (red cylinder) targeted to the pelvis of an immobilized mouse (skeleton in yellow). (**E**) Visualization of field size relative to pelvic organs of a male mouse. Organs were outlined on MRI and overlaid with bone windows from CT after rigid image registration. Organs depicted include bladder (yellow), prostate (pink), seminal vesicles (cyan), and colorectum (blue).

We next determined whether landmarks on the restrainer could be used to accurately localize the anatomic region to be targeted with radiotherapy. Autochthonous tumors arising in the prostate of genetically engineered mouse models can occur simultaneously in all prostate lobes. While the anterior and ventral lobes are easily identified on MR imaging, the lateral and dorsal lobes are less apparent. In order to define the borders of the prostate and its relationship to bony anatomy, both CT and MR imaging were correlated with whole mount histologic sections (Supplementary Fig. 1C). These studies showed that the inferior border of the prostate extends to approximately 2-3 mm above the pubic symphysis and the geometric center of the prostate is approximately 2-3 mm above the upper edge of the acetabulum (Fig. 1E, Supplementary Fig. 1D). Since both of these bony landmarks can be reproducibly positioned relative to reference points on the restrainer, we used them as a surrogate for the anatomical position of the prostate. We next identified the optimal position of the restrainer relative to the collimator aperture in order to center the prostate in the radiation field. Using reference points on the restrainer, the center of the prostate was able to be reproducibly aligned in the center of the radiation field (Supplementary Fig. 1E, F). Although a 1.5 cm diameter field is sufficient to target the entire normal prostate gland with a 2-3 mm margin, the size of the prostate increases with tumor growth and therefore a 2 cm diameter field was used to target autochthonous prostate tumors in mouse models (Fig. 1E, Supplementary Fig. 1D). This ability to achieve adequate tumor immobilization and reproducible positioning in conscious animals facilitates multi-fraction stereotactic radiotherapy, and eliminates the need for daily image-guidance and repeated exposure to general anesthesia.

### Dosimetry and verification of dose distribution in vivo

For the SABR platform to be generalizable, we used a widely available gamma-ray irradiator in which the irradiation chamber is centered between two motorized Cesium-137 sources located above and below the chamber (Fig. 1D). An advantage of this geometry is that the dose distribution within the animal is more homogeneous compared to a single beam arrangement in which the entrance dose can be significantly higher than the deep tissue dose due to beam attenuation. In addition, the narrow energy distribution of Cesium-137 radiation with a peak at 662 keV allows for a relative reduction in skin dose in the dose buildup region (∼1 mm), thus reducing dermatologic toxicity. A combination of optically stimulated luminescent dosimeters (OSLDs) and radiochromic film dosimetry were used to assess absolute dose and the dose distribution of the collimated radiation field (Supplementary Fig. 2A-I). Manufacturer reported dose uniformity for the entire sample chamber is ± 7%. Our measurements showed a dose uniformity of ± 2.3% in the central zone of the sample chamber where the animal restrainer is placed during treatment. Collimation of the radiation beam reduces the central dose rate by a factor that depends on the size of the collimated field. Based on OSLD measurements, the output factor for a circular field with a diameter of 1, 2, and 3 cm were 0.62, 0.72, and 0.8 respectively. Independent confirmation of the output factors by radiochromic film dosimetry was within 2% of these estimates. In the shielded zone the dose was attenuated by 95% (± 2%). Within the collimated field, radiochromic film dosimetry showed a homogeneous dose distribution and narrow penumbra (Fig. 2A and Supplementary Fig. 2 G-I).

**Figure 2.**
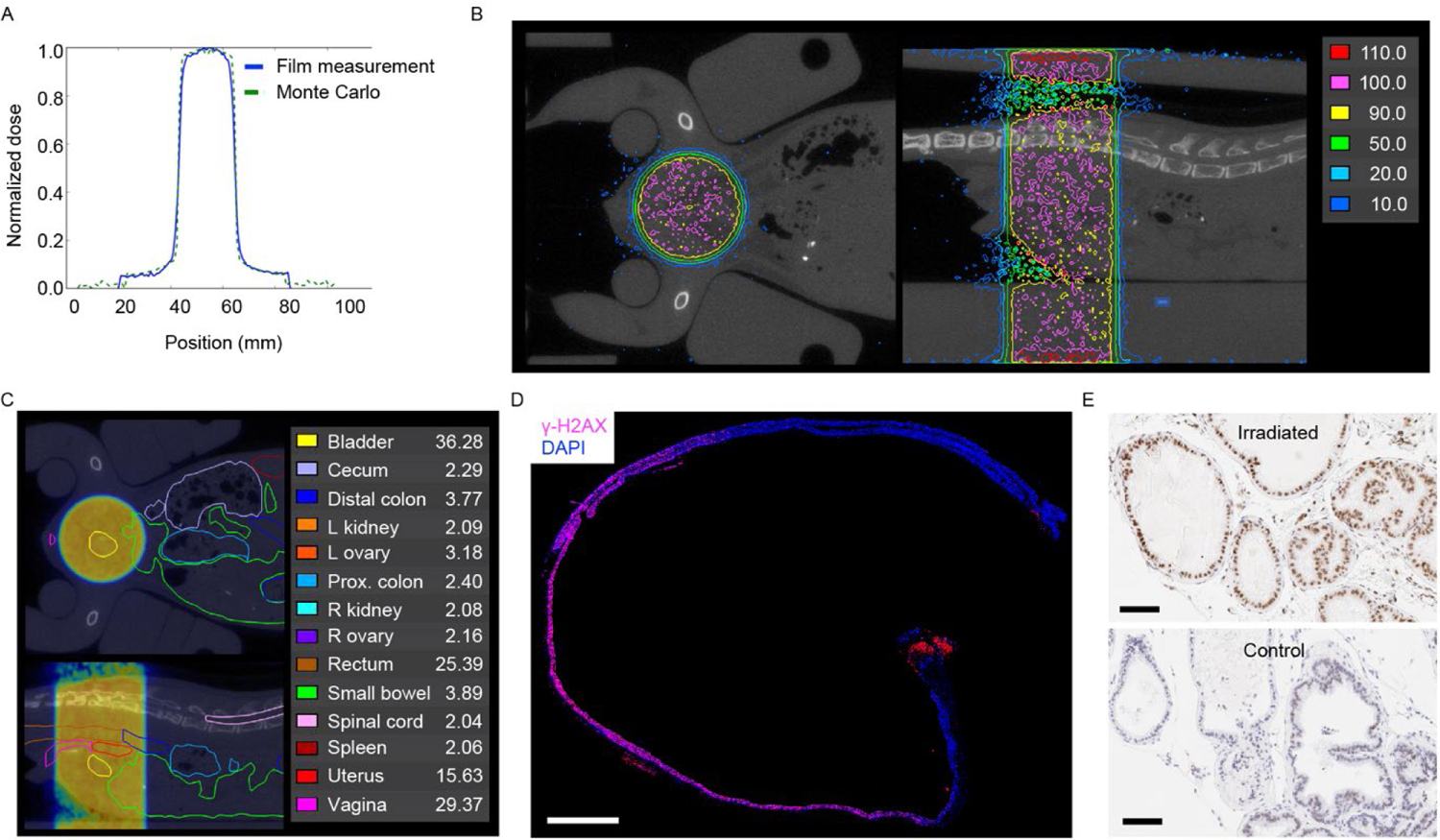
Modeling and verification of radiation dose distribution in mice. (**A**) Comparison of measured and modeled dose distribution for a 2 cm radiation field. (**B**) Representative dose distribution of a 1 cm circular pelvic radiation field mapped onto coronal (top view) and sagittal (side view) CT image of a female mouse. The reference isodose (100%) is shown in pink. Areas enclosed by the red line receive a dose >110% of the reference dose. Areas outside the dark blue line receive a dose <10% of the reference dose. (**C**) Coronal and sagittal abdominopelvic CT image of a female mouse with organs outlined as indicated. Mean organ dose in Gy is shown for a prescribed dose of 37.5 Gy. Dose distribution of a 1 cm circular pelvic radiation field is shown as a dose wash to allow visualization of organs. Dose color scheme as in (B). (**D**) Evaluation of focal DNA damage in a longitudinal cross-section of colorectum by γ-H2AX staining. This animal was treated with 15 Gy to a 2 cm field centered in the pelvis 30 minutes prior to tissue collection. Nuclear γ-H2AX foci are seen only in the portion of the colorectum that was within the radiation field. The red signal at the anus is due to autofluorescence. Scale bar = 2 mm. (**E**) Evaluation of focal DNA damage by γ-H2AX IHC in prostate glands. The animal was treated with 15 Gy to a 2 cm field centered over the prostate 30 minutes prior to tissue collection. The lower panel shows prostate tissue from an unirradiated control. Scale bar = 100 µm.

MRI is optimal for anatomic evaluation of the mouse abdomen and pelvis; however, it lacks electron density information needed to calculate absorbed radiation dose. We therefore used both MRI and μCT imaging to generate an accurate spatial dosimetric analysis (dose-volume relationship) for abdominopelvic organs. Organs outlined on MRI were registered to μCT to allow for dose calculation (Fig. 2B, C and Supplementary Fig. 2J). Treatment plans were generated for male and female mice with 1 and 2 cm radiation fields centered in the pelvis to simulate treatment of the prostate and rectum. Critical organs such as the kidneys and spinal cord received <10% of the prescribed dose (Fig. 2C and Supplementary Fig. 2 K-N). Phosphorylation of histone H2A.X at Ser139 (γ-H2AX) is a sensitive marker of DNA double strand breaks induced by ionizing radiation (27), and γ-H2AX immunohistochemistry (IHC) can be used to evaluate the focal effects of radiation in tissue. Consistent with the dosimetric predictions, we found homogeneous induction of γ-H2AX in prostate tissue and the portion of the colorectum that was within the radiation field, but not in adjacent colon and distal rectum that were outside the radiation field (Fig. 2 D, E). These data demonstrate that with appropriate collimation and immobilization, stereotactic focal radiation can be delivered to mouse models using widely available radiation sources.

### Determining the maximum tolerated dose of pelvic SABR

To evaluate the tolerance of pelvic organs in mice to SABR we performed a series of dose escalation studies. Single fraction and 2, 3, and 5-fraction regimens were evaluated at 4 dose levels for males and 5-fraction regimens at 3 dose levels for females (Table 1). To reduce toxicity and control for time effects, fractionated regimens were delivered over 8-9 days with equal number of days between each fraction. The biologically effective dose (BED) model (28) was used to calculate isoeffective doses for the different fractionation schemes (see methods for details). Isoeffective dose levels were set to account for late effects of radiation that cause permanent damage to normal tissues leading to organ failure due to loss of regenerative capacity (Supplementary Fig. 3A). Within each dose level, regimens with higher number of fractions, and thus higher total dose, are anticipated to have greater effects on proliferating tumor cells as well as proliferating cells in normal tissues leading to early-onset toxicity in bone marrow, intestine, and skin (Supplementary Fig. 3A). All animals were monitored for a minimum of 6 months and the maximum tolerated dose was defined as the highest dose at which at least 2 of 3 animals survived. For the 5-fraction regimen the maximum tolerated dose in males was 9 Gy per fraction and in females was 7.5 Gy per fraction (Table 1 and Fig. 3A).

**Figure 3.**
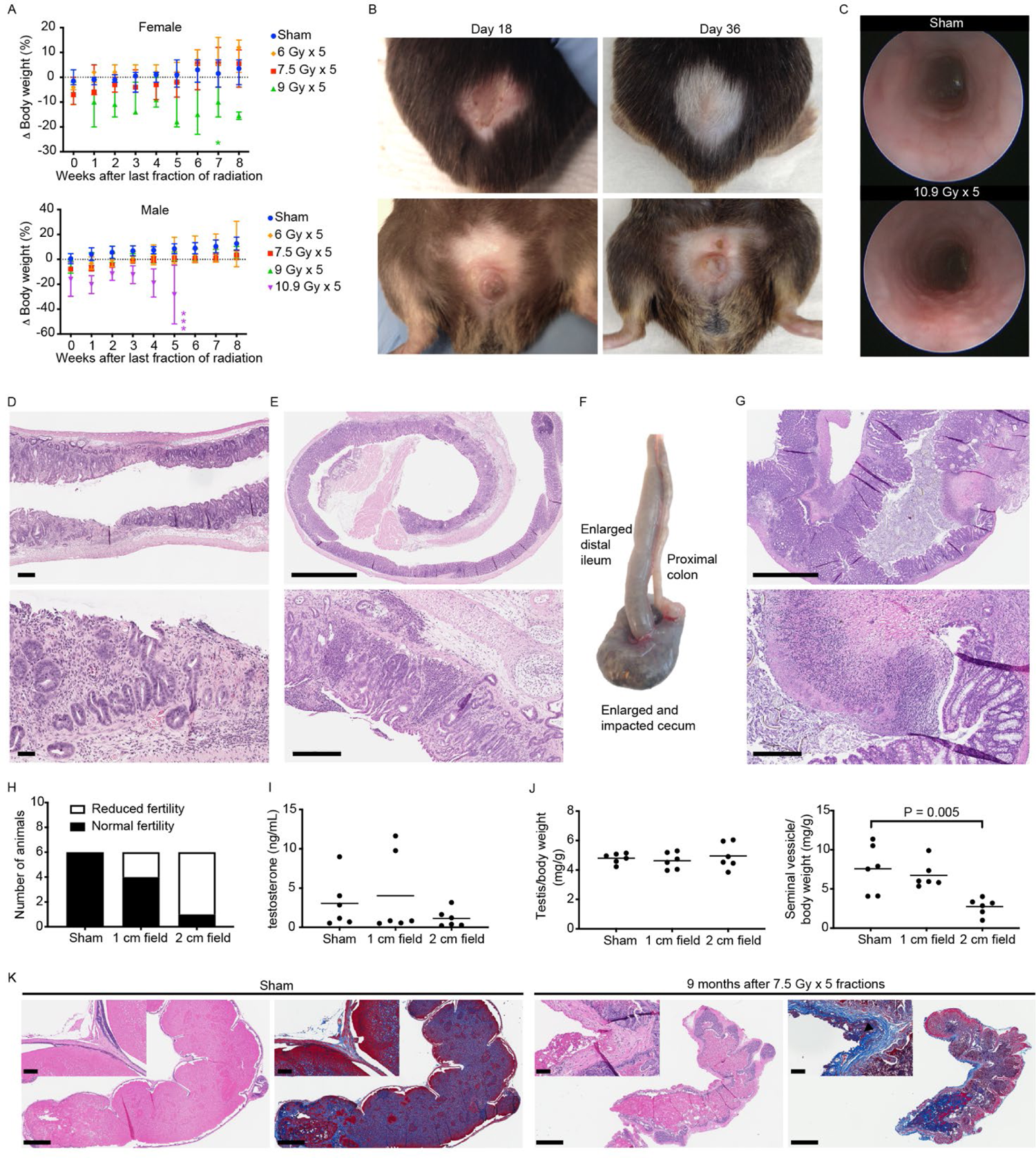
Acute dose-limiting toxicity and late effects of pelvic SABR. (**A**) Body weight change relative to pretreatment baseline for female and male mice treated with a 5-fraction regimen as shown. Median +/- 95% CI. Female n=6 for 0 and 7.5 Gy, n=3 for 6 and 9 Gy. Male n=9 for 0, 7.5, and 9 Gy, n=3 for 6 and 10.9 Gy. Each asterisk (*) indicates time of death/euthanasia of a single animal. (**B**) Representative skin changes in a male mouse at the indicated time points after completing 5 fractions of 10.9 Gy to a 2 cm pelvic field. Upper panels show dorsal side and lower panels ventral side. (**C**) Endoscopic images show colorectal mucosa 1 week after completing the regimen shown. (**D**) H&E shows loss of mucosal integrity in the rectum 5.5 weeks after completing 5 fractions of 10.9 Gy to a 2 cm pelvic radiation field. Scale bar = 250 µm for the low power view and 100 µm for the high-power view. (**E**) H&E shows severe chronic proctitis 4 months after completing 3 fractions of 14.4 Gy to a 2 cm pelvic radiation field. Scale bar = 2 mm for the low power view and 300 µm for the high-power view. (**F**) Gross pathology of intestines 11 weeks after completing 5 fractions of 9 Gy to a 2 cm pelvic radiation field. (**G**) H&E shows ulceration and inflammation in the cecum 11 weeks after completing 5 fractions of 9 Gy to a 2 cm pelvic radiation field. Corresponds to the specimen in F. Scale bar = 2 mm for the low power view and 300 µm for the high-power view. (**H**) Fertility of male mice 8-9 months after sham treatment or 5 fractions of 7.5 Gy to a 1 or 2 cm pelvic radiation field. (**I**) Serum testosterone in male mice 9 months after sham treatment or 5 fractions of 7.5 Gy to a 1 or 2 cm pelvic radiation field. (**J**) Wet weight of testis and seminal vesicles in male mice 9 months after sham treatment or 5 fractions of 7.5 Gy to a 1 or 2 cm pelvic radiation field. P value estimate by unpaired, two-tailed t test. (**K**) Representative sagittal section of one seminal vesicle 9 months after sham treatment or 5 fractions of 7.5 Gy to a 2 cm pelvic field. H&E and blue trichrome stain showing fibrosis (black arrowhead). Note that secretions in the seminal vesicle lumen also stain blue. Scale bar = 1 mm. Inset shows higher magnification of the seminal vesicle wall. Scale bar = 100 µm.

**Table 1.** Pelvic SABR dose escalation study. Dose levels and fractionation schemes evaluated in the pelvic SABR dose escalation study are shown, including number of fractions, dose per fraction, cumulative nominal dose, BED, and incidence of severe toxicity or death occurring within 6 months of treatment. Elapsed days is the total number of days over which radiation was delivered, including the first and last day. Radiation fractions were equally distributed across the elapsed days. The single fraction regimen at DL4 was not evaluated as the length of time in the restrainer would have exceeded the maximum time in the restrainer approved under our animal protocol. All mice were treated with a 2 cm diameter circular radiation field centered in the pelvis over the prostate and/or bladder. α/β, alpha-beta ratio is further described in the methods. DL, dose level; BED, biologic effective dose; N/A, not applicable; ND, not determined.

| Dose level | Number of fractions | Elapsed days | Dose per fraction (Gy) | Total Dose (Gy) | BED $\alpha/\beta = 3$ | BED $\alpha/\beta = 10$ | Incidence of severe toxicity/death | |
| --- | --- | --- | --- | --- | --- | --- | --- | --- |
|  |  |  |  |  |  |  | Males | Females |
| DL1 | 1 | N/A | 15 | 15 | 90 | 38 | 0 of 3 |  |
|  | 2 | 8 | 10.2 | 20.4 | 90 | 41 | 0 of 3 |  |
|  | 3 | 9 | 8.1 | 24.3 | 90 | 44 | 0 of 3 |  |
|  | 5 | 9 | 6 | 30 | 90 | 48 | 0 of 3 | 0 of 3 |
| DL2 | 1 | N/A | 18.3 | 18.3 | 130 | 52 | 0 of 3 |  |
|  | 2 | 8 | 12.6 | 25.2 | 131 | 57 | 0 of 3 |  |
|  | 3 | 9 | 10 | 30 | 130 | 60 | 0 of 3 |  |
|  | 5 | 9 | 7.5 | 37.5 | 131 | 66 | 0 of 3 | 1 of 3 |
|  | 5 | 5 | 7.5 | 37.5 | 131 | 66 | 1 of 5 | 3 of 5 |
| DL3 | 1 | N/A | 21.8 | 21.8 | 180 | 69 | 0 of 2 |  |
|  | 2 | 8 | 15 | 30 | 180 | 75 | 0 of 3 |  |
|  | 3 | 9 | 12 | 36 | 180 | 79 | 0 of 3 |  |
|  | 5 | 9 | 9 | 45 | 180 | 86 | 1 of 3 | 2 of 3 |
| DL4 | 1 | N/A | 26 | 26 | 251 | 94 | ND |  |
|  | 2 | 8 | 18 | 36 | 252 | 101 | 0 of 3 |  |
|  | 3 | 9 | 14.4 | 43.2 | 251 | 105 | 3 of 3 |  |
|  | 5 | 9 | 10.9 | 54.5 | 253 | 114 | 3 of 3 |  |

In line with predictions from the BED calculations, we found that at each dose level acute gastrointestinal and dermatologic toxicity occurred more frequently, and with greater severity, in animals treated with higher numbers of fractions. At dose level 4 (DL4), the highest dose level tested in males, all animals that received the 5-fraction regimen showed signs of acute dermatologic toxicity (alopecia and desquamation) that peaked between 2-3 weeks after radiation (Fig. 3B). At this dose level acute gastrointestinal toxicity was also apparent, including colorectal edema (Fig. 3C), loose stools, and hematochezia. By two weeks after radiation most animals recovered their baseline weight; however, at DL4 weight loss persisted and gradually worsened in animals treated with 3- and 5-fraction regimens (Fig. 3A and Supplementary Fig. 3B). All animals that received the 5-fraction regimen died or required euthanasia within six weeks of treatment (Fig. 3A). Necropsy revealed focal areas of bleeding and ulceration in the distal colon and rectum that was grossly apparent and confirmed by histology (Fig. 3D and Supplementary Fig. 3C). The bladder showed signs of mild-moderate edema (submucosal thickening) but no loss of epithelial integrity (Supplementary Fig. 3D). The prostate and seminal vesicles appeared largely normal, but in some cases had reduced luminal secretions (Supplementary Fig. 3D). The testis showed reduced sperm counts (Supplementary Fig. 3D). Animals in DL4 treated with the 3-fraction regimen died or required euthanasia within 3-5 months after radiation and showed histologic evidence of chronic damage to the colorectum (Fig. 3E and Supplementary Fig. 3B), while animals treated with two fractions at DL4 showed no signs of acute toxicity other than transient mild weight loss, but had evidence of chronic tissue damage at planned necropsy 6 months after treatment, including focal atrophy of prostate and seminal vesicles, fat necrosis in the epididymal fat pad, and focal atrophy of seminiferous tubules in one testis in 1 of 3 animals. (Supplementary Fig. 3D-G).

In contrast to the overt signs of acute gastrointestinal and dermatologic toxicity observed at DL4, the only consistent sign of acute toxicity at lower dose levels was decreased activity and characteristic weight loss that peaked two days after completing radiation (Supplementary Fig. 3H). Interestingly, animals that died or required euthanasia at DL3 did not have proctitis, but showed signs of chronic inflammation and loss of epithelial integrity in the cecum (Fig. 3F and G). In some cases, concurrent enlargement of the distal ileum was observed (Fig. 3F and Supplementary Fig. 3E). At this dose level, gastrointestinal toxicity occurred more frequently in female mice. Since age matched female mice are smaller than male mice and toxicity was seen more frequently in mice with lower baseline body weight, we hypothesize that the cause of increased toxicity in female mice was due to larger volume of cecum and small bowel in the treatment field. To test this, the field size was reduced to 1 cm and targeted to either the low abdomen or pelvis. The 5-fraction regimen targeted to the pelvis was well tolerated, yet several mice developed severe toxicity when the treatment field was targeted to the low abdomen (Supplementary Fig 3I). Together, these data argue that excess toxicity in female mice is due to increased bowel volume in the treatment field.

While the cause of death or severe toxicity in all cases was attributable to intestinal toxicity, we were surprised that at DL3 and below there were no overt signs of intestinal toxicity such as diarrhea or hematochezia. However, when intestines were examined at earlier time points, there were signs of moderate small bowel toxicity including edema, bloating, and gassy or frothy luminal contents, but no evidence of bleeding, ulceration, or stricture. Acute small intestine toxicity occurred more frequently at higher doses and in female mice (Supplementary Fig. 3J).

While the BED formula does not account for time effects, it is well established that accelerated radiation regimens are associated with greater incidence of acute toxicity. Indeed, we observed increased acute to subacute gastrointestinal toxicity when the DL2 5-fraction regimen was delivered daily (5 fractions in 5 days) rather than every other day (5 fractions in 9 days) resulting in higher incidence of treatment-related mortality (Table 1).

The late effects of radiation were assessed at 6 months for animals in DL2 and DL3 and at 12 months for DL1. Some animals at DL2 and DL3 showed small foci of fat necrosis or fibrosis, but there were no other gross signs of chronic tissue damage. Histologic evaluation showed no signs of chronic inflammation or fibrosis of bladder or rectum. Although some prostate glands appeared smaller, there was no evidence of senescence (Supplementary Fig. 3K, L). In some animals, the testis appeared smaller than untreated controls; however, there was no clear correlation between dose level and testis size. Germ cells are highly radiosensitive, as doses less than 1 Gy can cause prolonged azoospermia, while permanent sterility occurs in males at doses above 2 Gy (29). The variable results we observed suggested that the testis in some mice were likely close to or within the radiation field while others were completely shielded. Testosterone producing Leydig cells are less radiosensitive (29, 30); however, since altered testosterone levels could impact prostate cancer progression, we assessed the effects of pelvic radiation on circulating testosterone. Serum testosterone levels were not decreased in treated mice, but untreated C57BL/6 mice had very low baseline serum testosterone levels (Supplementary Fig. 3M). We therefore evaluated the effects of pelvic radiation on gonadal function in the outbred CD-1 strain, which reportedly have higher levels of serum testosterone (31). Four-month-old male CD-1 mice were treated with 5 fractions of 7.5 Gy, delivered every other day to either a 1 or 2 cm pelvic radiation field. Eight months after treatment each male was housed individually with a female of breeding age for one month. While all untreated controls fathered a litter over this time frame, only 4 of 6 mice treated with the 1 cm field, and 1 of 6 mice treated with the 2 cm field, produced offspring (Fig. 3H). Despite affecting fertility, we found no difference in serum testosterone or testis volume in irradiated mice relative to controls (Fig. 3I, J). Furthermore, on histologic evaluation spermatogenesis appeared to be normal (Supplementary Fig. 3N). However, the seminal vesicles, anterior prostate, and ventral prostate were on average smaller in mice treated with the 2 cm radiation field (Fig. 3J and Supplementary Fig. 3O). Histologic evaluation confirmed decreased prostate and seminal vesicle secretions and showed evidence of fibrosis (Fig. 3K and Supplementary Fig. 3P). Taken together, these data suggest that when the radiation field is targeted to the prostate, reduced fertility is not due to effects on gonadal function, but rather due to direct effects of radiation on the prostate causing fibrosis and decreased secretory function.

In summary, our SABR dose escalation studies indicate that the colorectum is the dose limiting organ in the pelvis with acute lethal injury occurring with 5 fractions of 10.9 Gy. The distal colon and rectum tolerate doses below this, but small intestine and proximal large intestine develop subacute to chronic toxicity with 5-fraction regimens in the 7.5-9 Gy range, with severity likely dependent on the volume of intestine within the radiation field. In contrast the bladder, prostate, and seminal vesicles are relatively resistant to these doses, yet in some cases develop fibrosis as a late effect of radiation.

### Hematologic and immunologic effects of focal pelvic radiation

An advantage of GEMMs is that the effects of tumors and therapy on the native immune system can be studied. Thus, a series of studies were conducted to determine the effect of SABR on circulating blood cells and immune cells in spleen and lymph nodes. For all of these studies radiation was targeted to 2 cm circular field centered in the pelvis.

In circulation, red blood cells (RBCs) are considered to be the most radioresistant, while white blood cells (WBCs), particularly lymphocytes, are the most radiosensitive (32). Accordingly, we found only a small decrease in RBCs after radiation that normalized 8 weeks after treatment (Fig. 4A), although the female cohort treated with 5 fractions of 9 Gy showed persistent anemia due to hematochezia. Platelet levels reached a nadir one week after radiation and normalized by week 8 (Fig. 4A). In contrast WBCs declined by more than 50% within 2 days of completing radiation and the decline was primarily due to decreased lymphocytes (Fig. 4A). Neutrophil counts normalized relatively quickly, while lymphopenia persisted longer, but also resolved by week 8 (Fig. 4A). Increased neutrophil counts at 8 weeks in females treated with 5 fractions of 9 Gy may be due to chronic intestinal injury and inflammation.

**Figure 4.**
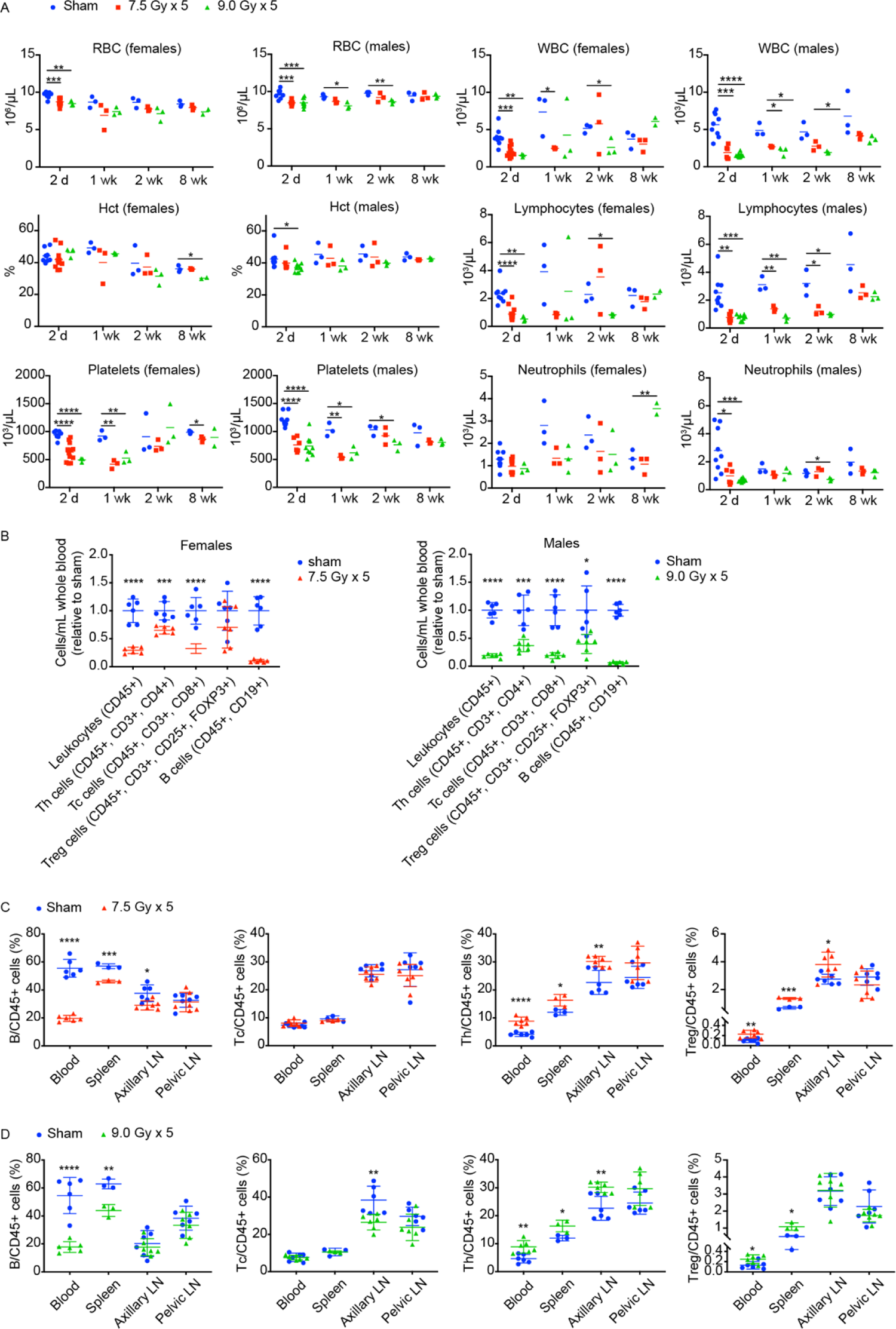
Hematologic effects of pelvic SABR. (**A**) Blood cell counts at the indicated number of days or weeks after completion of 5 fractions of radiation or sham treatment. For females n=3-12 for 2-day time point and n=3 for all other time points (except n=2 for 9 Gy x 5 group at 8-week time point due to 1 treatment-related death). For males n=6-9 for 2-day time point and N=3 for all other time points. (**B**) Lymphocyte subtype counts determined by multiplexed flow cytometry in whole blood 2 days after completion of 5 fractions of radiation or sham treatment as indicated. Mean +/- SD. Data are normalized to the mean of untreated controls. (**C**) Lymphocyte subtypes as a percent of total leukocytes 2 days after completion of radiation in females. Mean +/- SD. (**D**) Lymphocyte subtypes as a percent of total leukocytes 2 days after completion of radiation in males. Mean +/- SD. For all panels * p<0.05, ** p<0.01, ***p<0.001, ****p<0.0001 by unpaired 2-tailed t test.

Further characterization of circulating lymphocyte populations revealed the greatest decrease in B cells, with T cells being more radioresistant (Fig. 4B). T helper cells (Th) also appeared to be less sensitive than cytotoxic T cells (Tc), while regulatory T cells (Tregs) showed the least relative decrease relative to controls (Fig. 4B). Similar relative changes in lymphocyte subtypes occurred in spleen, albeit to a lesser extent than in blood, perhaps due to the spleen being outside the radiation field (Fig. 4C, D). Axillary lymph nodes, which were also outside the radiation field, showed minimal changes in relative lymphocyte populations, while pelvic lymph nodes that were within the radiation field showed no relative differences in lymphocyte subtypes (Fig. 4C, D). An ability to assess absolute changes in lymphocyte populations in these organs by flow cytometry was not possible; however, lymph node weight and 2-dimentional morphometric analysis indicated that there was a trend toward smaller pelvic lymph nodes two days after radiation, the time point when maximum changes in lymphocyte populations were observed in blood (Supplementary Fig. 4A-C). At the maximum tolerated dose in females (5 fractions of 7.5 Gy) there was no measurable effect on total mass of spleen, axillary or pelvic lymph nodes (Supplementary Fig. 4A). At the maximum tolerated dose in males (5 fractions of 9 Gy) there was a small but measurable effect on total mass of spleen and axillary lymph nodes, but not pelvic lymph nodes at 2 days after completing the radiation course (Supplementary Fig. 4B).

### Efficacy of SABR regimens in human-derived mouse xenografts

Having established that pelvic radiotherapy in mice using potentially curative SABR regimens has an acceptable toxicity profile, we next evaluated efficacy of 4-5 fraction regimens in human prostate cancer-derived mouse xenografts. Nude mice with bilateral flank xenografts were immobilized using the custom restrainer to allow targeting of one of the flank tumors (Fig. 5A). In all cases tumor regression was observed in the treated flank while the tumor in the shielded flank progressed (Fig. 5B-D). Some SABR-treated tumors showed slow regrowth between 3-4 weeks after treatment but most remained smaller than their pretreatment size by week 4, and some regressed completely (Fig. 5B-E). SABR treated prostate cancer xenografts that did not regress completely still showed a reduction in proliferative index compared to untreated controls (Supplementary Fig. 5A). Eight of 14 (57%) clinical complete responders (cCR) also had a pathologic complete response (pCR) defined as no viable tumor cells present on histologic examination of the tumor implantation site (Supplementary Fig. 5B). Although PC3 xenografts were more likely than 22Rv1 xenografts to have a complete response, PC3 xenografts in our study were also smaller at time of treatment and there was a clear trend toward better responses in smaller tumors (Fig. 5E). To determine if pathological complete responders could be considered “cured,” we used bioluminescence imaging (BLI) to monitor for recurrence or metastasis of luciferase-expressing PC3 xenografts after treatment with SABR. BLI showed no evidence of local recurrence or metastasis 5 months after treatment indicating that these animals did indeed remain disease-free for a prolonged period after treatment (Supplementary Fig. 5C).

**Figure 5.**
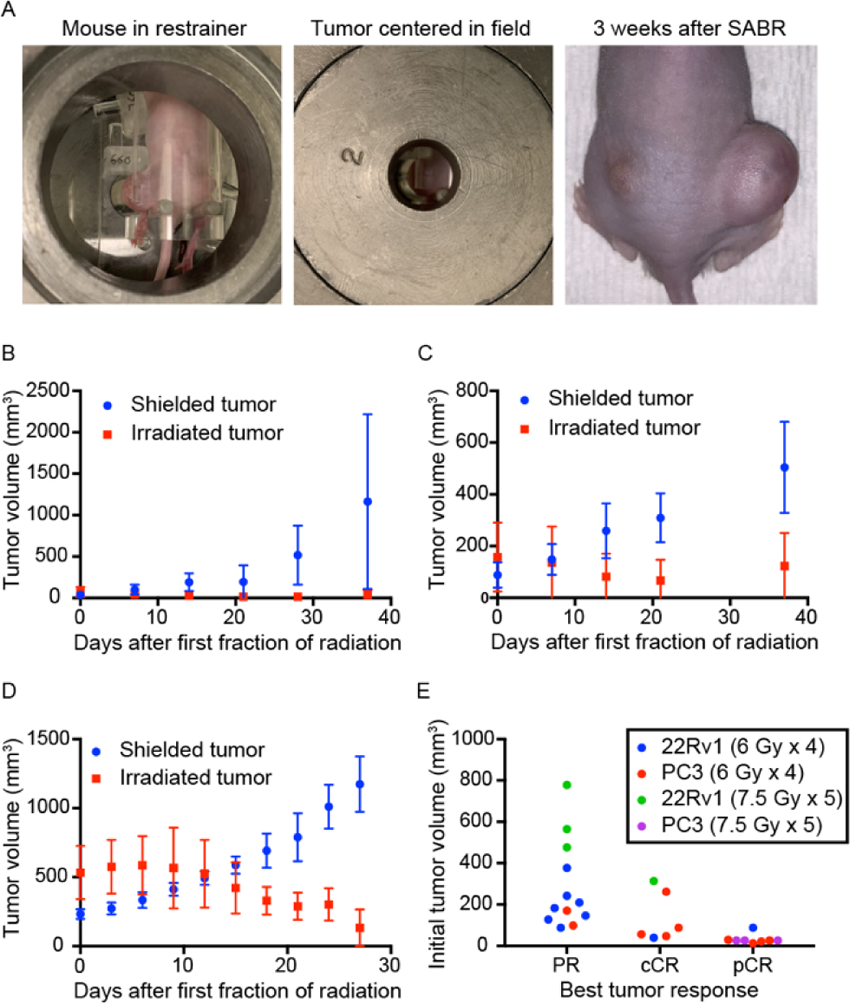
SABR can induce complete responses in flank xenografts. (**A**) Nude mouse with bilateral flank 22Rv1 xenografts showing the larger flank tumor immobilized for radiation and centered in a 2 cm radiation field. Image on the right shows treatment response 3 weeks after completing radiation therapy. (**B**) Growth of PC3 xenografts (n=5) treated with 4 fractions of 6 Gy QOD to the larger flank tumor as shown in (A). Mean tumor volume +/- SD. (**C**) Growth of 22Rv1 xenografts (n=5) treated with 4 fractions of 6 Gy QOD to the larger flank tumor as shown in (A). Mean tumor volume +/- SD. (**D**) Growth of 22Rv1 xenografts (N=4) treated with 5 fractions of 7.5 Gy QOD to the larger flank tumor as shown in (A). Mean tumor volume +/- SD. (**E**) Best tumor response by initial tumor size. PR, partial response; cCR, clinical complete response; pCR, pathological complete response.

### Therapeutic radiation improves survival in a prostate cancer GEMM

*PTEN* and *TP53* are commonly deleted or mutated tumor suppressors in prostate cancer, and their loss correlates with poor prognosis (33–37). In mice, combined deletion of *Pten* and *Trp53* in prostate epithelium is sufficient to drive tumorigenesis with short latency resulting in rapidly progressive and lethal prostate adenocarcinoma (38, 39). Using an established GEMM (*Pten^flox/flox^; Trp53^flox/flox^; Pbsn-Cre* herein referred to as *Pten;Trp53^pc-/-^)* we confirmed that conditional loss of Pten and p53 in mouse prostate epithelium results in locally aggressive prostate cancer that is universally fatal. Macroscopic tumors, visible by MRI, developed as early as 4 months of age (Fig. 6A) and tumors were palpable by 5 months of age. All mice died or required euthanasia due to poor body condition score between 6 and 8 months of age. Necropsy showed no evidence of thoracic or abdominopelvic metastasis. In some cases, pelvic lymph nodes appeared mildly enlarged, but were found to be reactive on histologic evaluation. In all cases the cause of death could be attributed to local mass effect of the prostate tumor resulting in genitourinary or gastrointestinal obstruction (Fig. 6B, C).

**Figure 6.**
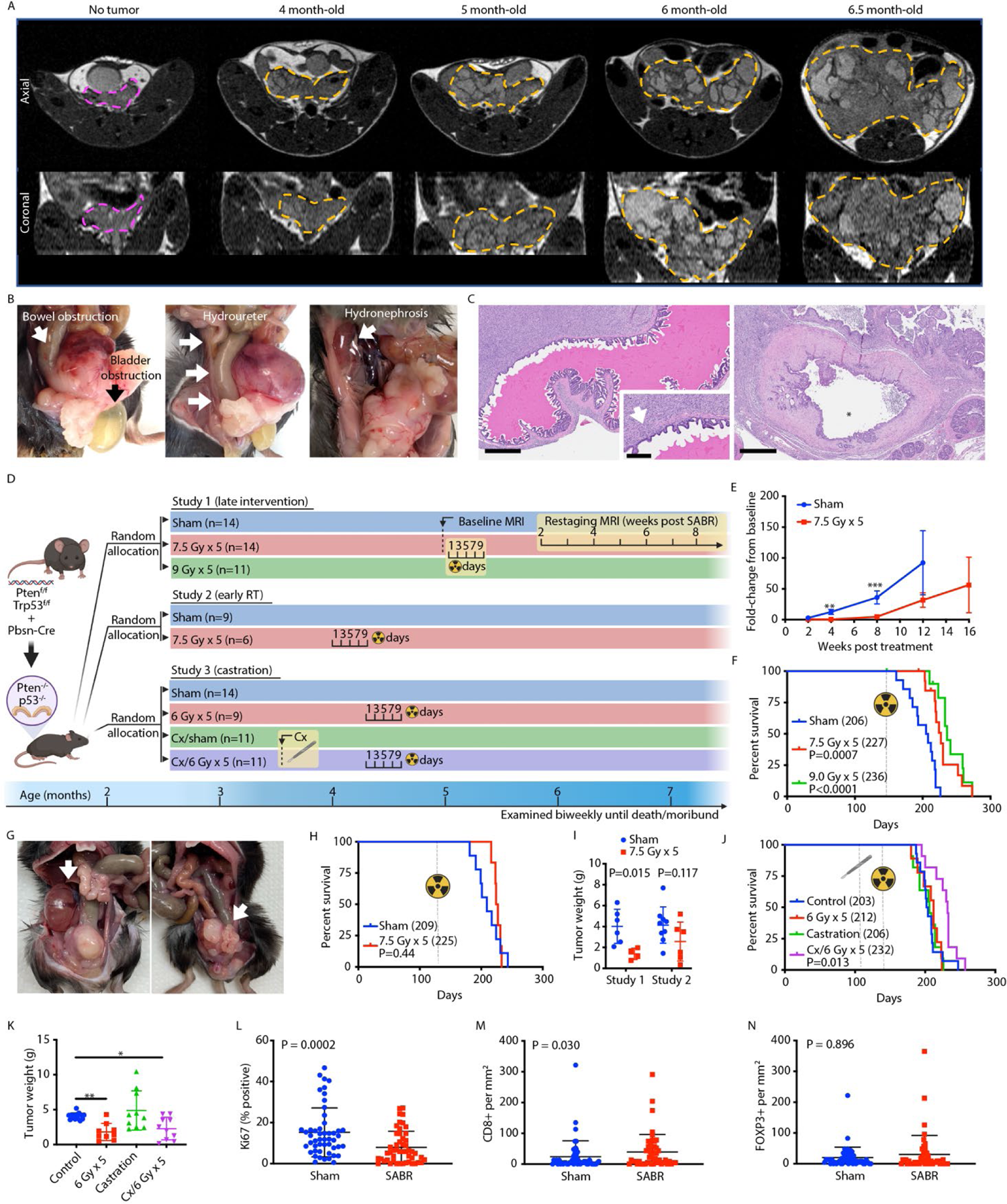
SABR modestly improves survival an autochthonous model of locally aggressive prostate cancer. (**A**) Tumor progression in *Pten;Trp53^pc-/-^* mice assessed by serial MRI. Representative axial (upper) and coronal (lower) views with prostate tumor outlined in orange. In all cases the axial and coronal slice with the largest square area of tumor is shown. A 3-month-old mouse without visible prostate tumor outlined in magenta is shown for comparison. (**B**) Obstruction of bladder, ureter, and rectum documented at time necropsy in untreated mice. (**C**) Histologic sections showing seminal vesicle invasion (arrow) and compressed bladder with small lumen (*) engulfed by tumor. Scale bar = 600 um (main image) 200 um (inset). (**D**) Graphical representation of prostate cancer GEMM and study design, including timing of interventions. Cx, castration. (**E**) Tumor progression assessed by MRI in study 1 as defined in D. n=7 for sham cohort, n=14 for SABR cohort. Mean +/- SEM. **p<0.01, ***p<0.001 by unpaired two-tailed t test. (**F**) Cancer-specific survival in study 1. Event defined as death or euthanasia due to tumor burden. Animals with cause of death other than tumor burden were censored (2 in each radiation cohort developed severe gastrointestinal toxicity requiring euthanasia). Number in parenthesis = median survival in days. P value estimate by log-rank test. (**G**) Hydronephrosis (left) and rectal obstruction (right) documented at time necropsy in mice treated with SABR. (**H**) Overall survival in study 2 specified in D. Number in parenthesis = median survival in days. P value estimate by log-rank test. (**I**) Tumor weight at time of necropsy in study 1 (SABR at 5 months) and study 2 (SABR at 4 months). Mean +/- SD. P value estimate by unpaired two-tailed t test. (**J**) Overall survival in study 3 specified in in D. Number in parenthesis = median survival in days. P value estimate by log-rank test. (**K**) Tumor weight at time of necropsy in study 3. Mean +/- SD. P value *p<0.05, **p<0.01 by unpaired two-tailed t test. (**L**) Percent Ki-67 positive cells assessed by IHC in SABR-treated tumors and controls (n=5). For each tumor 10 high power fields (HPF) were analyzed. Mean -/+ SD. P value estimate by Mann-Whitney test. (**M**) CD8+ T cells assessed by IHC in SABR-treated tumors and controls (n=5), 10 HPF per tumor. Mean -/+ SD. P value estimate by Mann-Whitney test. (**N**) FOXP3+ T cells assessed by IHC in SABR-treated tumors and controls (n=5), 10 HPF per tumor. Mean -/+ SD. P value estimate by Mann-Whitney test.

To test whether SABR is effective in this prostate cancer model, *Pten;Trp53^pc-/-^*male mice were randomized to 5 fractions of 0, 7.5, or 9 Gy (Fig. 6D, study 1) at 5 months of age, a time point when tumors were clinically apparent. SABR significantly delayed tumor progression assessed by serial MRI (Fig. 6E, Supplementary Fig. 6A, B) and median cancer-specific survival was extended in a dose dependent manner (Fig. 6F). Although pilot studies suggested that 5 fractions of 9 Gy was tolerable in tumor bearing mice, a few animals in the radiation cohorts developed severe gastrointestinal toxicity requiring euthanasia (Fig. 6F). In all cases these mice had very small tumors at time of necropsy indicative of treatment effect. Necropsy of SABR-treated animals that died or required euthanasia due to tumor progression showed overall smaller primary tumors (Fig. 6I); however, death was still caused by mass effect on organs adjacent to the prostate (bladder, ureters, rectum) (Fig. 6G). There was no evidence of macroscopic metastases even in animals that survived beyond 8 months.

Given that tumor size at the time of radiation correlates with treatment effect (Fig. 5E) we reasoned that the marginal improvement in survival we observed was due to high tumor cell burden at time of treatment. To determine whether treating tumors at an earlier stage would improve outcomes, *Pten;Trp53^pc-/-^* mice were randomized to 5 fractions of 7.5 Gy or sham irradiation (Fig. 6D, study 2) at 4 months of age, a time point when tumors first become radiographically apparent, but are not palpable. The median tumor size by MRI at time of treatment was 52 mm^3^ (range 25-258 mm^3^). Surprisingly, we did not find that earlier treatment improved survival or tumor burden at time of death (Fig. 6H); in fact, there was a trend toward large tumors in the early intervention cohort (Fig. 6I).

Randomized clinical trials have demonstrated that combining radiation with androgen deprivation therapy (ADT) for prostate cancer improves outcomes over radiation alone (40, 41). To determine whether androgen deprivation improves response to radiotherapy we conducted a study in which mice were surgically castrated prior to SABR (Fig. 6D, study 3). A pilot study showed a high incidence of penile edema and prolapse in animals treated with combination therapy. Therefore, castrated animals were allowed to recover for 3 weeks prior to initiating radiotherapy and the radiation dose was limited to 5 fractions of 6 Gy. Notably, at this dose, radiation alone did not improve survival despite decreasing tumor size (Fig. 6J, K). However, median survival was prolonged with combination therapy to a similar extent as high dose SABR monotherapy regimens (Fig. 6F, J). Castration alone did not improve survival or tumor burden in this model, consistent with prior reports (Fig. 6J, K) (39, 42). These results show that although *Pten*;*p53*-null tumors are resistant to ADT, this model still recapitulates the synergy between radiation and ADT observed in human tumors.

Consistent with smaller tumors in SABR treated animals, the number of Ki67 positive cells was significantly lower on average in irradiated tumors (Fig. 6L). Yet, in some regions, irradiated tumors had a similar proliferation index as untreated controls suggesting that tumor repopulation was occurring at the time tumors were collected (Supplementary Fig. 6C). We noticed that proliferation tended to be highest in areas of the tumor near normal prostate glands and these areas also frequently showed prominent lymphocytic infiltrates (Supplementary Fig. 6D).

To further characterize tumor infiltrating lymphocytes (TILs) and understand how they are affected by radiotherapy, relevant T cell populations were examined by IHC. Although cytotoxic T cells were overall rare in tumors (in most cases <50 cells per mm^2^), they were more abundant in regions near the tumor periphery and adjacent to normal glands (Supplementary Fig. 6E). Furthermore, levels were slightly higher on average in irradiated tumors (Fig. 6M). Regulatory T cells were also rare in tumors, with a few areas showing higher levels, generally where cytotoxic T cells were also more abundant (Supplementary Fig. 6F). Although regulatory T cells trended higher in irradiated tumors, unlike cytotoxic T cells, the difference was not significant (Fig. 6N). Taken together these results indicate that prostate tumors in the *Pten;Trp53^pc-/-^*model are characterized by low, yet heterogeneous levels of TILs, and suggest that TIL activity may be most relevant at a time when tumor regrowth is occurring in irradiated tumors.

### Therapeutic radiation improves survival in a colorectal adenoma GEMM

Since radiation therapy plays an important role in the management of non-metastatic rectal cancer, we also evaluated SABR in a mouse adenoma model in which tumors are induced by focal deletion of *Apc* in intestinal mucosa (43). Under endoscopic guidance *Apc^f/f^;Villin^CreERT2^* mice were injected with 4-hydroxytamoxifen at a single site 2-3 cm from the anal verge to induce colorectal tumors as previously described (44). We confirmed that tumors induced in this location could be accurately targeted with a 2 cm pelvic radiation field (Supplementary Fig. 7A, B). Although these tumors do not metastasize, they are characterized by slow, persistent growth that can ultimately lead to death due to gastrointestinal bleeding, bowel obstruction, and/or rectal prolapse. In a pilot study, a treatment regimen of two fractions of 15 Gy (BED_3_ = 180) was effective at decreasing tumor burden, but evidence of late fibrosis was found 6 months post treatment. Thus, 5 fractions of 7.5 Gy (BED_3_ = 130), which is well tolerated in male and female mice (Fig. 3A), was used for treatment. After stratifying based on tumor size, mice were randomized to radiation or sham treatment and then followed for a minimum of 6 months (Fig. 7A, study 1). While more than half of the sham-treated mice died due to bowel obstruction or required euthanasia due to severe rectal prolapse, none of the SABR-treated mice developed bowel obstruction or rectal prolapse (Fig. 7B and C). At endpoint, some tumors were exophytic while others were flatter and more infiltrative making it difficult to compare tumor burden between individual mice; however, irradiated tumors were generally smaller and had less fluorodeoxyglucose (FDG) uptake on PET scan than controls, yet none showed complete regression and there was no clear difference in the proliferation index (Fig. 7D and Supplementary Fig. 7C).

**Figure 7.**
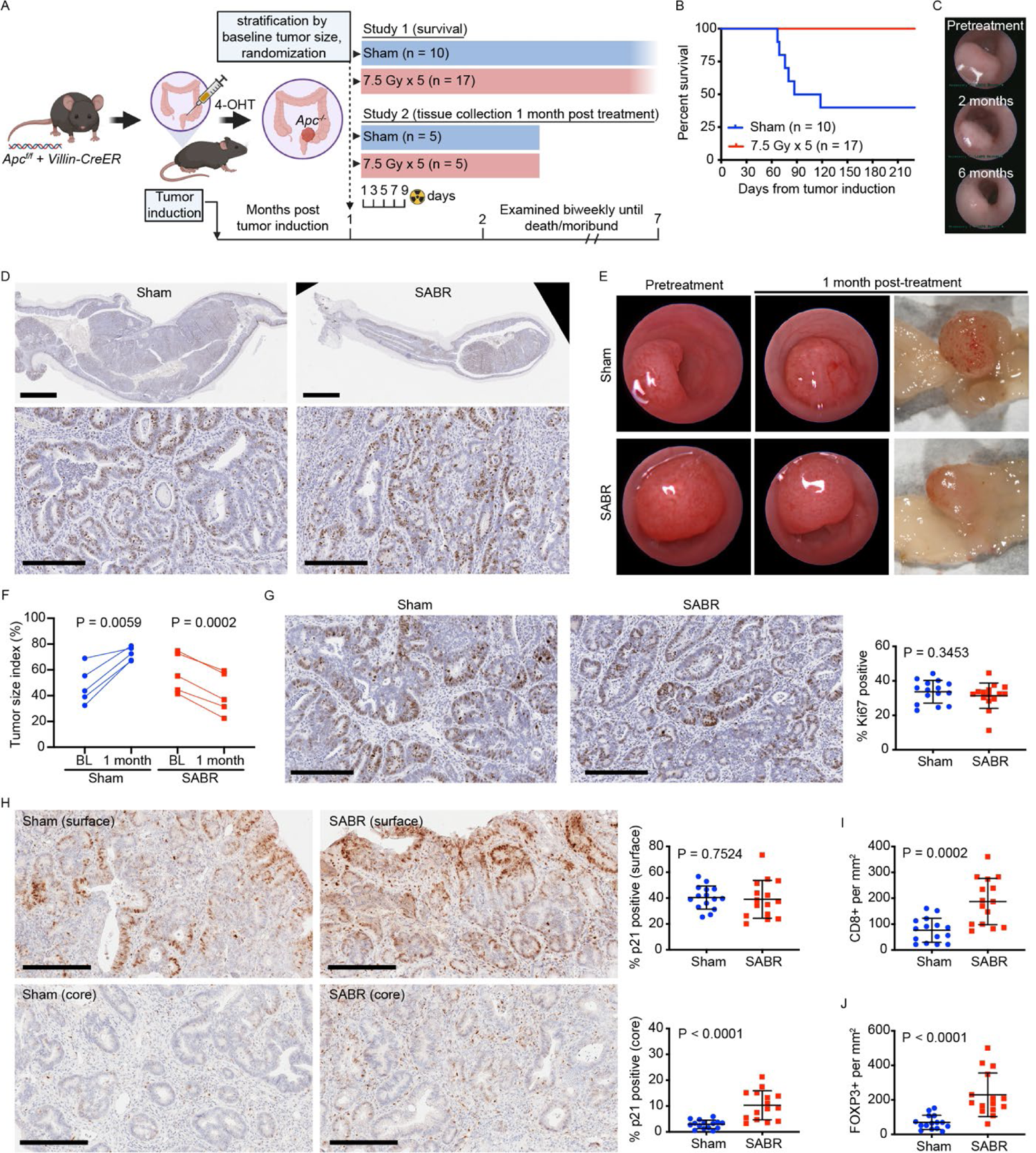
SABR improves survival in an autochthonous model of colorectal adenoma. (**A**) Graphical representation of colorectal cancer GEMM and study design, including timing of interventions. Study 1 was conducted with approximately equal numbers of male and female mice that were 2-3 months of age at time of tumor induction. Study 2 involved only male mice. 4-OHT, 4-hydroxytamoxifen. (**B**) Overall survival in study 1 specified in A. (**C**) Representative endoscopic images of rectal adenoma prior to treatment and at the indicated time points after SABR. (**D**) Representative Ki67 IHC of a rectal adenoma 7 months after radiation (8 months after tumor induction). A tumor from an untreated animal two and a half months after tumor induction is shown for comparison. Scale bar = 2 mm (low power view), 0.2 mm (high power view). (**E**) Representative endoscopic and gross images of a SABR-treated tumor and untreated control from study 2 specified in A. (**F**) Size of individual tumors from study 2 as evaluated by endoscopy at baseline (BL) and 1 month post SABR. N=5 for both treatment groups. P value estimate by paired, two-tailed t test. (**G**) Representative Ki67 IHC images and proliferative index of tumors from study 2. Three high power fields per tumor were assessed. Scale bar = 0.2 mm. P value estimate by Mann-Whitney test. (**H**) Representative p21 IHC images and quantitation of p21 positive cells at the surface and core region of tumors from study 2. Three high power fields per region/tumor were assessed. Scale bar = 0.2 mm. P value estimate by unpaired, two-tailed t test. (**I**) Quantitation of cytotoxic T cells assessed by IHC in tumors from study 2. Three high power fields per tumor were assessed. P value estimate by unpaired, two-tailed t test. (**J**) Quantitation of regulatory T cells assessed by IHC in tumors from study 2. Three high power fields per tumor were assessed. P value estimate by unpaired, two-tailed t test.

To determine how radiation was affecting tumors to improve survival in this model, a second study was conducted with a planned endpoint one month after treatment to allow earlier histopathologic evaluation (Fig. 7A, study 2). At this timepoint, tumor size was different between treatment groups (Fig. 7E), and all irradiated tumors demonstrated regression, while all untreated tumors grew over the same time interval leading to near complete luminal obstruction (Fig. 7E, F). Surprisingly, no difference in the proliferation index, as assessed by Ki-67 immunostaining, was found in irradiated tumors (Fig. 7G). We also found no evidence of apoptosis (Supplementary Fig. 7D). Based on these findings, we hypothesized that cells in SABR-treated tumors were undergoing proliferation arrest. Since, *Apc*-null adenomas have intact p53 signaling, and p21 (CDKN1A) is key mediator of p53-mediated growth arrest in response to DNA damage in colorectal cancer cells (45, 46), we investigated p21 expression by IHC in adenomas one month post treatment. Interestingly, we found spatial heterogeneity in p21 expression in the adenomas. Tumor cells near the luminal surface showed high levels of p21 expression, and there was no difference between SABR-treated tumors and controls. In contrast, adenoma cells deeper in the tumor had low p21 expression; however, the core of irradiated tumors had a significantly higher fraction of p21 positive cells, the majority of which were in the stroma (Fig. 7H).

Since immune cells can exhibit anti-tumor effects independent of apoptosis (47) we quantified TILs by IHC. Cytotoxic T cells were present in intraepithelial and stromal compartments while regulatory T cells were restricted to the stromal compartment, and both were increased in irradiated tumors (Fig. 7I, J and Supplementary Fig. 7E, F). These data suggest that SABR affects tumor stroma and immune cells in a way that could contribute to therapeutic effects of radiation in this autochthonous colorectal cancer model.

## DISCUSSION

Response to radiotherapy depends on both cell intrinsic and cell extrinsic factors. Nonetheless, how radiation alters the tumor microenvironment is incompletely understood. GEMMs provide an opportunity to study tumors as they develop in a relevant tissue environment, yet have been underutilized to investigate radiotherapy as part of potentially curative regimens used in the clinic. The approach described here illustrates how the response of tumors and normal tissues to curative doses of radiation, with and without adjuvants, can be studied in autochthonous mouse cancer models. Although effects on pelvic tumors were investigated, similar approaches could be used to treat tumors in other anatomic locations, including the upper abdomen, thorax, head and neck.

One of the barriers to conducting preclinical radiotherapy trials in GEMMs has been the time and cost required to deliver clinically relevant regimens using existing radiotherapy platforms. Apart from cost and ease of use, an advantage of the platform described here is that animals are conscious during treatment, thus avoiding repeated exposure to anesthesia that could affect animal health and alter the efficacy of radiation therapy. Moreover, due to the predictable anatomy of laboratory mice, pretreatment imaging is not required to target autochthonous tumors with focal radiotherapy provided that appropriate immobilization and stereotactic technique are used.

### Normal tissue effects

The acute effects of whole body radiation on various organ systems in mice have been well documented (48) but may contrast from tolerance of individual organ systems to focal radiotherapy at clinically relevant doses. Mice exposed to 6-10 Gy whole body radiation die within 2-3 weeks from hematologic injury, while whole body doses in the 10-15 Gy range cause death within a week of exposure due to gastrointestinal toxicity (48). In contrast, the current study shows that focal pelvic radiation is well tolerated with no observable signs of toxicity after a single dose of 15 Gy to the pelvis. The most apparent acute radiation toxicity observed with 5-fraction pelvic SABR regimens was weight loss that occurred within a few days of treatment initiation, peaked within 2 days of completion, and typically resolved 1-2 weeks thereafter. While the cause of weight loss is likely multifactorial, it coincided with acute hematologic and gastrointestinal toxicity. The only other overt sign of radiation effects were dermatologic changes that manifested within 2 weeks after the highest dose level tested and were much less pronounced and presented later at lower dose levels.

Gastrointestinal toxicity is commonly observed in patients treated with radiotherapy to the abdomen and/or pelvis and may occur early (within 3 months of treatment) or late (months to years after treatment) with significant correlation between the incidence and severity of early and late toxicity (49). In both humans and mice the incidence and severity of GI toxicity depends on the volume of intestine irradiated, cumulative radiation dose, and length of time over which the radiation is delivered. Single fraction doses of 15 Gy or more to the entire abdomen in mice results in acute lethality due to gastrointestinal leak syndrome (50, 51). Smaller segments of intestine can tolerate higher doses, and fractionation with longer intervals between fractions increases the total dose that can be delivered to the intestine (52). While the pathogenesis of acute gastrointestinal leak syndrome following whole abdomen radiation is an area of active investigation (53–56), intestinal toxicity from focal radiotherapy is less well studied (20, 57, 58). In the 5-fraction regimen used here no small bowel toxicity was observed at doses up to 9 Gy per fraction (BED_10_ = 85.5) targeted to a 2 cm pelvic radiation field in male mice. Female mice developed mild-moderate toxicity, primarily bowel enlargement and edema involving the ilium, with 5 fractions of 7.5 Gy (BED_10_ = 65.6) and have a greater incidence and severity of this toxicity at higher doses. The reason for the sex differences is likely the smaller volume of the intestine within the radiation field of male mice.

When compared to small bowel, the large bowel and rectum are more tolerant of radiation. Consistent with prior reports (59, 60), single fraction doses less than 20 Gy targeted to the distal colon and rectum were well tolerated. For fractionated regimens the highest dose level tested caused acute rectal toxicity (ulceration, colitis) resulting in 100% mortality around 5 weeks of treatment with the 5-fraction regimen (BED_10_ = 114) and between 3 to 5 months with the 3-fraction regimen (BED_10_ = 105). However, at lower dose levels only subtle histologic changes were observed in the colorectum. These results are consistent with older mouse toxicity studies using fractionation schemes not commonly used in the clinic, but of comparable effective dose (61, 62).

In contrast to early GI toxicity observed in the high dose groups, GU toxicity was not apparent in the acute phase at any dose level; however, fibrosis and atrophy was seen at later time points. The etiology of these changes was not a focus of this study, yet histologic evaluation implicated altered stromal biology. We found no evidence of senescence, perhaps due to removal of senescent cells by the immune system (63). The testis, which is in close proximity to the prostate was at least partially exposed to radiation in some mice as evidenced by histologic changes consistent with decreased spermatogenesis, a process that can be affected by doses as low as 1 Gy (64). Importantly, most animals studied had no evidence of impaired spermatogenesis, validating the precision of this radiation platform.

Regarding acute systemic hematologic effects, our findings recapitulate some findings involving single-fraction whole body radiation and show that similar changes occur with clinically relevant SABR regimens (32, 65, 66). Consistent with differences in radiosensitivity among blood cell lineages reported by others (67) (68–70) the highest relative decrease and most prolonged effect was observed for lymphocytes. B cells were the most sensitive lymphocyte population, while the relative radiosensitivity amongst T cells was Tc > Th > Treg. An open question is whether the effects we observed on peripheral blood cells are caused by radiation effects on cells circulating through the radiation field or due to depletion of hematopoietic progenitors in the radiation exposed bone marrow. Two lines of evidence suggest the former. Firstly, the timing of the observed changes relative to the half-life of circulating blood cells is inconsistent with marrow suppression; secondly, the fraction of functional bone marrow within a 2 cm pelvic radiation is relatively small and should be compensated for by bone marrow outside the radiation field (71). While it is formally possible that background radiation (estimated to be 5-8% of the prescription dose) could affect lymphocyte populations outside the radiation field, such doses are unlikely to affect erythrocyte, granulocyte, and megakaryocyte precursors in the bone marrow (67). Interestingly, similar effects on peripheral blood cells have been observed in humans receiving pelvic radiotherapy as well as a variety of other treatment sites, where again the timing of changes are more suggestive of effects on peripheral circulating blood cells rather than bone marrow suppression (72, 73).

In addition to lymphocytes in circulation we also evaluated lymphocyte subsets in spleen and lymph nodes within and outside the radiation field. Although not within the radiation field, the spleen showed similar, albeit less pronounced changes in lymphocyte populations as in whole blood, namely a relative decline in B cells and increase in Th and Treg cells; however, no relative changes were seen in axillary or pelvic lymph nodes. Taken together these data suggest that lymphocytes in lymph nodes may have a different radiosensitivity profile than circulating lymphocytes and that some lymphatic tissues outside the radiation field may undergo transient lymphodepletion. Although morphometric analysis suggested that there was a relative lymphodepletion of in-field lymph nodes, we could not demonstrate this based on lymph node tissue weights, possibly due to the small size of pelvic lymph nodes and considerable variation in lymph node weight between animals at baseline. Interestingly we observed a small but statistically significant decrease in weight of spleen and lymph nodes outside the radiation field in mice treated with 5 fractions of 9 Gy but not 7.5 Gy. While these changes may have been the result of background radiation, an alternate possibility is that lymphocyte populations outside the radiation field were mobilized in response to peripheral lymphocyte depletion.

To summarize these findings, pelvic SABR in mice using clinically relevant doses is feasible and has some characteristic hematologic effects similar to those observed with lower dose whole-body radiation. Pelvic SABR regimens consisting of 5 fractions with BED_3_ = 250 Gy uniformly result in acute lethal toxicity within 6 weeks of treatment, where the dose limiting organ is the distal colon. In contrast, regimens consisting of 5 fractions with BED_3_ = 180 Gy rarely result in subacute lethal toxicity, where the dose limiting organ is the distal small intestine and proximal colon. While these studies primarily involved C57BL/6J mice, which are of intermediate sensitivity with regard to whole body radiation effects (74), a limited number of studies were also done in outbred CD-1 mice with similar results. However, caution is advised in extrapolating these results to strains known to be more sensitive to ionizing radiation (75), or animals with underlying comorbidities, genetic alterations, or undergoing concurrent therapy, where toxicity is likely to be increased. In these situations, it is advisable to begin with regimens in the BED_3_ = 90-130 range with a maximum field size of 2 cm and minimum body weight of 20 grams, while the field size should be decreased for smaller mice.

### Tumor responses to SABR

Notably, potentially curative SABR regimens improve survival in genetically engineered mouse models of rectal adenoma and locally aggressive prostate cancer. In prostate cancer, we also observed synergy between radiation and androgen deprivation that is consistent with findings from clinical practice (40, 41). However, in both cases tumors never regressed completely, and in the case of the aggressive *Pten;Trp53^pc-/-^*prostate cancer model, progressed rapidly after therapy.

In human prostate cancer the frequency of p53 mutations increases with disease progression and is associated with worse prognosis; yet, the role of p53 in prostate cancer progression and treatment resistance remains incompletely understood (76–78). In culture models, loss of p53 can confer resistance to multiple therapies (79), and p53 loss may confer radio-resistance to SABR in GEMMs. However, complete ablation of p53-null PC3 xenografts was achieved with doses that were ineffective in the *Pten;Trp53^pc-/-^* model, arguing that factors other than loss of p53 must be involved in resistance to radiation in this context.

Although the studies were not specifically designed to examine mechanisms of radiation resistance, the finding that treating autochthonous tumors at an earlier stage and smaller size did not improve outcomes is notable. This contrasts with flank xenograft models where tumor control probability was associated with initial tumor size. Furthermore, upon autochthonous tumor regrowth there was heterogeneity in proliferative index, with some areas showing similar proliferation as untreated tumors. This suggests that even small autochthonous tumors may harbor radioresistant persister cells capable of repopulating a tumor after SABR. It has been shown that genetically engineered models do not acquire significant mutations other than the targeted driver mutation (80), and since treatment is completed within a matter of days it is unlikely that newly acquired mutations contribute to treatment resistance. Rather transient cell states are likely to underlie treatment resistance.

Tumor associated stroma in the rectal adenoma model was unexpectedly altered by SABR treatment. This raises the question of whether effects on tumor associated stroma contribute to the therapeutic effects of radiation in this model. Tumor stroma consists of multiple cell types, including cancer associated fibroblasts, immune cells, and vascular cells, all of which can be affected by radiation and also interact with each other in complex ways (81–84). Given the effects observed on tumor cells and stroma, the rectal adenoma model described here may be a model of how radiation alters tumor stroma to shape response to therapy.

In the era of immunotherapy, how radiation and other cytotoxic therapies interact with the immune system is an area of active investigation (85, 86). In preclinical models, radiation has been shown to have both immunostimulatory and immunosuppressive effects (84, 87, 88). The GEMMs of prostate and rectal cancer used in our study had variable amounts of TILs, including cytotoxic T cells. Interestingly, TILs were increased after SABR in both the prostate and rectal cancer models, implying that the immune system may help define how autochthonous tumors respond to SABR. This increase was observed even at time points with transient reduction in circulating lymphocytes. In the rectal model both Tc and Treg cells were increased, while in prostate only Tc cells showed a statistically significant increase. These findings are in line with prior reports of increased intratumoral Tc and Treg cells after radiation in syngeneic mouse xenograft models and an apparent increased survival of intratumoral T cells compared to lymphocytes in circulation and in lymphoid tissues (89–91). Importantly, our data show that this phenomenon also occurs in autochthonous tumors, which have an intact tumor stroma that can impact TIL recruitment (92).

## Conclusions

Disease recurrence after curative therapy remains a major challenge in the clinic. Multiple processes have been implicated, including treatment resistant persister cells and an immunosuppressive tumor microenvironment. We show that clinically relevant SABR regimens can be delivered safely to mouse cancer models, but in some cases are unable to cure autochthonous tumors. Thus, these models can be used to examine how cancer evades radiation treatment and develop alternative approaches to improve therapeutic efficacy.

## MATERIALS AND METHODS

### Mouse strains, husbandry, and tumor induction

Mice were housed in a specific pathogen-free (SPF) facility at the Koch Institute at MIT with ad lib access to standard chow and water. The light cycle was 7 AM to 7 PM. C57BL/6J breeders were purchased from Jackson Laboratories (RRID:IMSR_JAX:000664). CD-1 breeders were purchased from Charles-River (Strain Code: 022). Wild-type mouse colonies were expanded in house and re-crossed with founders from Jackson Laboratories or Charles-River at least once per year to prevent genetic drift. *Pten^f/f^* (MGI: 2679886) (93), *Trp53^f/f^* (MGI: 1931011) (94), and *Pbsn-Cre* (MGI:2385927) (95) mice were maintained on a mixed C57BL/6J × 129SvJ background. *Pten^f/+^; Trp53^f/f^; Pbsn-Cre* males were bred with *Pten^f/f^; Trp53^f/f^* females to generate *Pten^f/f^; Trp53^f/f^; Pbsn-Cre* male mice for prostate cancer GEMM studies. *Apc^f/f^*(MGI: 3688435) (96) and *Villin^CreERT2^* (MGI: 3053826) (97) mice were maintained on a pure C57BL/6J background. To induce colorectal tumors, 50-100 µL of 100 µM 4-hydroxytamoxifen (Calbiochem Cat. # 579002) in PBS was injected into the submucosa of 6-8 week-old male and female *Apc^f/f^;Villin^CreERT2^* mice under endoscopic guidance 2-3 cm from the anal verge. Injections were performed using a custom injection needle (Hamilton Inc., 33G, small Hub RN NDL, 16 inches long, point 4, 45-degree bevel, part # 7803-05), a syringe (Hamilton Inc., part # 7656-01), a transfer needle (Hamilton Inc., part # 7770-02), and a colonoscope with integrated working channel (Richard Wolf 1.9 mm/9.5 French pediatric urethroscope, model # 8626.431). Prostate cancer xenograft studies were conducted with male CD-1 nude mice (Crl:CD1-Foxn1nu/nu) mice purchased from Charles-River laboratories (Strain Code: 086). PC3 cells (RRID:CVCL_0035) were obtained from the Broad Institute cell line repository, STR tested using the ATCC Cell Line Authentication Service, and passaged in DMEM (Corning Cat # 10-013-CV) supplemented with 10% heat-inactivated fetal bovine serum (Sigma). 22Rv1 cells (RRID:CVCL_1045) confirmed by STR testing were kindly donated by Dr. Massimo Loda (Dana Farber Cancer Institute) and passaged in RPMI (Corning Cat # 15-040-SV) supplemented with 2 mM glutamine (Invitrogen) and 10% heat-inactivated fetal bovine serum (Sigma). Both cell lines were routinely tested for mycoplasma. Cells harvested from cultures in exponential growth phase were resuspended in PBS, mixed 1:1 with Matrigel (Corning Cat # 356231) and injected subcutaneously into the caudal flank of 8-10 week-old male CD-1 nude mice. Each flank was injected with 1.5 million PC3 cells or 3 million 22Rv1 cells.

### Radiation delivery and dosimetry

Radiation was performed on a Gammacell 40 Exactor (Best Theratronics) located in the same SPF facility where the mice were housed. This instrument has a ventilated circular sample chamber (diameter 31.2 cm, height 10.5 cm, volume 8.0 L) that is centered between two Cesium-137 sources located 68 cm apart. During the course of this study the dose rate was between 0.9 – 1.0 Gy per minute. Mice were immobilized in the restrainer for up to 30 minutes during radiation delivery. Only one animal was irradiated at a time. Following irradiation or sham treatment, animals were returned to their home cage with ad lib access to standard chow and water and were monitored daily for two weeks and weighed at least once per week for 2 months. Thereafter body condition and activity were assessed once weekly. Any animals showing signs of distress or poor body condition score were euthanized by CO2 inhalation.

Thermoluminescent dosimeters (TLDs) were purchased from Radiation Dosimetry Services at MD Anderson (Houston, TX, USA). TLDs were sandwiched between clinical bolus material (0.5 cm thick), placed in the restrainer in the collimated or open field, and irradiated for 0 to 6 minutes. TLDs were promptly returned to Radiation Dosimetry Services for analysis. TLD measurements were performed in triplicate at 3 dose levels. Confirmatory experiments were performed using optically stimulated luminescent dosimeters (OSLDs) purchased from Landauer.

Radiochromic film dosimetry was performed using Gafchromic EBT-3 film (Ashland) and Epson 10000XL scanner. Gafchromic EBT-3 film was placed between plexiglass sheets such that the film was located in the central plane of the collimated field as shown in Supplementary Fig. 2D-F. Film dosimetry calibration was generated by exposing film in an open field (between plexiglass sheets placed on a Styrofoam block) in the same irradiator over a dose range between 0 and 8 Gy. Independent dose-response functions were determined for each of the color channels in the scanned film, and overall dosimetry was calculated through a joint fitting procedure. To evaluate the dose distribution horizontally and vertically along the central axis of the radiation field, Gafchromic EBT-3 film was placed between plexiglass sheets (6 mm thick) and irradiated with a range of doses between 0 and 8 Gy. In total, 5 films and 6 plexiglass sheets were used to cover the full height of the effective radiation field as shown in Supplementary Fig. 2E. Two independent radiochromic film measurements were performed with 2-3 dose levels per experiments.

### Surgical Castration

Male mice were anesthetized with isoflurane and placed on a warmed surface in the surgical suite. Using aseptic technique, the testis was exposed via 1 cm vertical midline incision in the ventral scrotum and 0.5 cm incision in the tunica. The spermatic blood vessels and vas deferens were cauterized after which the testis and epididymis was removed. Remaining tissue was gently returned into the scrotum and the process was repeated with the other testis via the same scrotal incision. The tunica was closed with tissue glue and the scrotal incision closed with a single wound clip. After recovery from anesthesia mice were returned to a clean cage. Analgesic (Carprofen) was administered once daily for 3 days. Wound clips were removed between day 7-10.

### Radiobiologic calculations

The linear-quadratic model is used describe the biologic effect of fractionated radiation regimens on tumors and normal tissues (98). The linear-quadratic (LQ) formula is second-degree polynomial with a linear and a quadratic term that is fitted to empirical clonogenic survival data in order to determine the coefficient of the linear term (alpha) and the coefficient of the quadratic term (beta). In its simplest form the LQ formula describes the relationship between radiation dose and effect on clonogenic survival

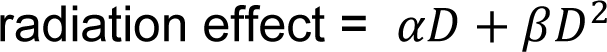

where D is the radiation dose, ⍺ is the coefficient of the linear term, and β the coefficient of the linear term. The ratio of alpha to beta (⍺/β) is the dose at which the linear and quadratic terms contribute equally to the radiation effect, which in turn reflects capacity for DNA repair. Most tumors and proliferating tissues are generally considered to have a high alpha/beta ratio (⍺/β ≥ 10) while slow growing tumors and quiescent tissues are considered to have a low alpha/beta ratio (⍺/β ≤ 3). In clinical practice the alpha/beta ratio is used to calculate iso-effective doses for a different dose-fractionation regimens (28). The formula that is used for this is the biologically effective dose (BED)

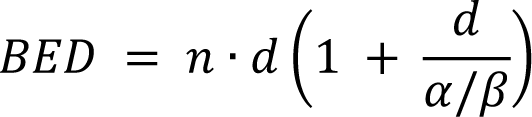

where *d* is the dose per fraction, *n* is the number of fractions, and ⍺/β is the alpha/beta ratio derived empirically for a given cell line, tumor, or tissue by fitting the survival curve to the LQ formula as described above. In the current study we used ⍺/β = 3 to calculate the biologically effective dose (BED_3_) for quiescent tissues where radiation toxicity develops at late time points. For early-responding tissues we used ⍺/β = 10 to calculate the biologically effective dose (BED_10_). While additional terms can be incorporated in the BED formula to account for time over which the radiation course is delivered and kinetics of tumor repopulation, these are not commonly used in clinical practice (99). Moreover, the LQ model has been shown to be appropriate for determining iso-effective fractionated radiation regimens with doses up to 18 Gy per fraction, but is less reliable for higher doses and single fraction regimens (100).

### Mouse imaging and radiation treatment planning

Magnetic Resonance Imaging (MRI) was performed on a 7T MRI system (7T/210/ASR, Agilent/Varian). Mice were anesthetized by inhalation of 3% isoflurane and maintained on 1-2% isoflurane throughout data collection with heated air delivery. Axial proton density weighted images were obtained using fast spin echo sequence (fsems) with the following parameters: TR/TE = 2000/12 ms, ETL = 4, 256×256 matrix, FOV = 40×40 mm2, interleaved number of slices = 30 to 50, no gap and slice thickness = 0.5 mm, number of averages = 2. Scans were collected with respiratory gating (PC-SAM version 6.26 by SA Instruments Inc.) to minimize motion artifact. Contouring of normal tissues for defining the radiation field size and dose-volume analysis was performed in MIM Version 6.0 (MIM Software Inc.).

For image registration and dosimetric studies an eXplore CT 120 scanner (GE Healthcare) was used to acquire micro computed tomography (µCT) images with X-ray tube voltage 70kVp, current 50.0 mA, and exposure time of 32 ms. Data was acquired over a 360-degree rotation with a step size of 0.5 degrees. Detector binning of 2×2 resulted in an isotropic resolution after reconstruction of 50 microns (Parallax Innovations, GPU-based reconstruction). After immobilization in the restrainer, mice were anesthetized by inhalation of 3% isoflurane and maintained on 1-2% isoflurane introduced via nosecone into the front end of the restrainer. Mice were then imaged by MRI, followed immediately by µCT. Contours of abdominopelvic organs were generated on axial MRI images. MR-CT image registration was performed in MIM. MRI and CT images were co-registered using rigid fusion to the upper pins of the restrainer, after which MR structure set was transferred to CT. The treatment isocenter was placed mid-plane between the top and bottom of the restrainer and centered 3 mm above the top edge of the upper pins for the 1 cm circular field, or 8 mm above the top edge of the upper pins for the 1 cm circular field. A Monte Carlo dose distribution model based on radiochromic film dosimetry was generated. In brief, we modeled the Gammacell irradiator and our collimation system in the EGSnrc software package (101). Monte Carlo dose distributions were normalized using the OSLD dosimetry measurements, and validated through comparison with beam profiles extracted from our film dosimetry. Treatment plans for a parallel opposed beam arrangement were generated for male and female mice and both 1 cm and 2 cm diameter circular radiation fields. Dose (37.5 Gy) was prescribed to the isocenter. Dose-volume histograms were generated for all major abdominopelvic organs, and the maximum, minimum, and mean doses for each organ were calculated. MIM was used for treatment planning and generating dose-volume histograms. VelocityAI version 3.1 (Varian Medical Systems Inc.) was used for visualization of the dose distribution and MRI-defined structure.

For assessment of inter-fraction setup error conscious animals were secured in the restrainer and imaged on the eXplore CT 120 µCT scanner. Repeat scans for the same animal were obtained on 3 separate days. DICOM files were imported into MIM and bony anatomy was outlined on each scan using the threshold tool. Images were co-registered using rigid fusion to the upper pins of the restrainer. Contours of the bony anatomy were transferred to a single CT scan to compare alignment of the pelvic bones between scans.

For assessment of intra-fraction motion a Skyscan 1276 (Bruker) was used to acquire serial images with a stationary gantry. 1200 images were acquired over a period of approximately 3 minutes. Images were acquired with x-ray tube settings of 100 kV, 200 mA, and an exposure time of 33 ms with a 0.5 mm aluminum beam filter. 8×8 detector binning was used for an isotropic resolution after reconstruction of 80.3 microns in NRecon 2.0 (Bruker). Mice were imaged twice in succession; first while awake and free breathing, then immediately thereafter while under 2.5% isoflurane anesthesia. The two-dimensional (2D) grayscale images collected were analyzed to determine the amount of movement during each imaging period. It was observed that the subjects would experience brief periods of movement separated by relatively long motionless intervals. Calculating a median value for each pixel position during the imaging period results in a neutral position representing the neutral, resting position of the subject. With this as a reference, the deviation from the neutral position was calculated for each image by first applying a threshold for bone to both images and then calculating the absolute value of the difference of the two images. The thickness of non-zero regions of this image were taken to represent the distance the bone moved. The maximum thickness, representing maximum displacement, in each image was used to represent the overall movement in that image. This was expressed as a percent time spent at a given displacement from the neutral position. Further, this number was limited to include only a 2 cm diameter region centered in the pelvis. Data analysis was done in MATLAB (MathWorks).

For bioluminescence imaging, mice were injected subcutaneously with 150 mg/kg D-luciferin (Caliper Life Sciences). After 10 minutes they were anesthetized by inhalation of 3% isoflurane and maintained on 1.5% isoflurane introduced via nosecone. Mice were imaged on an IVIS Spectrum in vivo optical imaging system (PerkinElmer) equipped with a heated platform to maintain a physiological body temperature during imaging. Images were collected every 5 minutes over a 40 minute period in order to capture the period of maximum photon emission. Living Image 3.1 (PerkinElmer) was used for image processing.

For PET imaging, mice were fasted overnight and dosed intravenously with ∼80 µCi of F-18 fluorodeoxyglucose (PETNET Solutions, Woburn, MA) via tail vein. Mice were kept anesthetized at 2% isoflurane for 1 hour uptake at 37°C and then imaged for 10 minutes PET acquisition and 2 minutes CT using a G8 PET/CT system (Perkin Elmer). Images were reconstructed using default MLEM 3D protocol and CT attenuation correction and were visualized in AMIDE Version 1.0.5.

### Colonoscopy and volumetric analysis of tumors

Optical colonoscopy was used to document colorectal mucosal toxicity in the dose escalation studies and to monitor tumor size in the colorectal adenoma studies, as previously described (44, 102). Endoscopic evaluation of tumor size was performed three weeks after tumor induction. Animals were stratified into 4 groups by baseline tumor size and the largest and smallest groups were excluded. Animals in the two middle groups were separately randomized to radiation or sham treatment to ensure similar baseline tumor size between groups. Tumors were imaged with optical colonoscopy before radiation treatment and after. Offline images were analyzed with FIJI (103) and the Tumor Size Index (TSI) was calculated as:

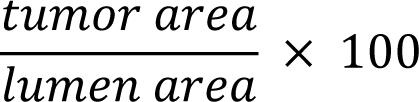

For xenograft studies the length, width, and height of xenografts was measured using digital calipers. All measurements were performed by a single operator. Tumor size was calculated as the volume of an ellipsoid

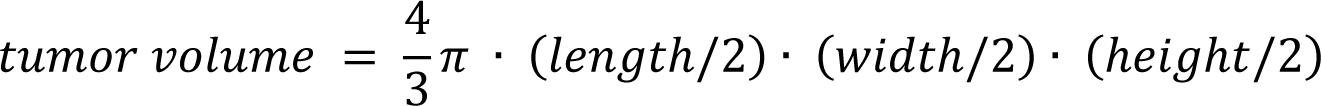

MRI was used for autochthonous prostate tumor monitoring. A coronal scout was obtained to estimate tumor size after which axial images were collected one slice inferior to the pubic symphysis to one slice above the cranial edge of the tumor. Tumors were contoured using the ROI tool in OsiriX Version 7.0.2 (Pixmeo Sarl) to determine tumor volume.

### Histology and Immunohistochemistry

Tissues collected at time of necropsy were fixed in 10% neutral buffered formalin for a minimum of 3 days at 4°C. Formalin fixed tissues were dehydrated, paraffin embedded, sectioned, and stained with hematoxylin and eosin (H&E) or Masson’s Trichrome. For whole-mount histology, a male mouse was removed from the restrainer after CO2 euthanasia, fixed in Bouin’s fixative for 1 week at 4°C, decalcified in 0.5 M EDTA pH 8.0 solution for 3 weeks, step sectioned (approximately 750 μm between sections) and H&E stained.

For IHC the following primary antibodies were used: Ki-67 (Biocare Medical, Cat# CRM 325, RRID:AB_2721189) diluted 1:50 in TBST, Cleaved Caspase-3 (Asp175) (Cell Signaling, Cat# 9664, RRID:AB_2070042) diluted 1:800 in TBST, CD8 (Cell Signaling, Cat# 98941, RRID:AB_2756376) diluted 1:400 in TBST, FoxP3 (Cell Signaling, Cat# 12653, RRID:AB_2797979) diluted 1:200 in TBST, Phospho-Histone H2A.X (Ser139) (Cell Signaling, Cat# 9718, RRID:AB_2118009) diluted 1:600 in TBST, p21 (Abcam, Cat# ab188224, RRID:AB_2734729) diluted 1:1000 in TBST. For γH2AX and p21 IHC 5 µm thick formalin-fixed paraffin-embedded (FFPE) sections were deparaffinized, rehydrated, and immediately underwent heat-mediated antigen retrieval a pressure cooker (Biocare Medical) at 125°C for 5 minutes in Citra pH 6.0 solution (Biogenex, HK086) for γH2AX, or in Tris-EDTA (pH 9.0) antigen retrieval solution (Abcam, Cat # ab93684) for p21. After cooling to room temperature, sections were equilibrated in distilled water prior to processing. Endogenous peroxidase activity was quenched with BLOXALL (Vector Labs, SP-6000) for 20 minutes. Sections were then blocked for 30 minutes with 3 percent normal goat serum, incubated overnight with primary antibody at 4°C, incubated with avidin/biotin/HRP reagents per manufacturer recommended protocol (Vector Labs, ABC-HRP Kit, Cat# PK-4001), incubated with DAB substrate (Vector Labs, Cat# SK-4100) for 5 minutes at room temperature, and counterstained with hematoxylin. Ki67, Cleaved caspase-3, CD8, and FoxP3 IHC was performed on the Thermo Scientific LabVision 360 autostainer. Antigen retrieval was done using the PT module (LabVision) at 97°C for 20 minutes in citrate buffer pH 6 (Abcam, Cat# 3678), followed UltraVision Hydrogen Peroxide Block (ThermoScientific, TA-125-H202), Rodent Block M (Biocare Medical, #RBM961), primary antibody for 1 hour, Rabbit on Rodent HRP (Biocare Medical #RMR622), DAB Quanto (ThermoScientific, #TA-125-QHDX) for 5 minutes, and counterstained with hematoxylin. For visualization and image processing, slides were scanned using an Aperio AT2 digital slide scanner (Leica Biosystems) at 20X magnification.

For γH2AX immunofluorescence, primary antibody incubation was done as above. Sections were then incubated in the dark at room temperature for 30 minutes with goat anti-rabbit Alexa Fluor 568 (Invitrogen Cat # A-11036, RRID:AB_10563566) diluted 1:50 in TBST, mounted in VECTASHIELD with DAPI (Vector Laboratories, Cat # H-1200), and imaged with an inverted fluorescence microscope (Nikon Eclipse Ti-S, SPOT RT3 camera, SPOT5.2 software). Images were taken with 10X objective in phase contrast, DAPI, and TRITC channels. Channels were merged in ImageJ Version 1.53a (NIH) and images were stitched in Canvas version 9 (ACD Systems) to generate the composite shown in Fig. 2D.

Senescence-associated β-galactosidase staining was performed according to protocol (104). Freshly collected tissue was immersed in OCT and frozen on dry ice. A cryotome was used to cut 4 µm thick sections that were fixed in 1% paraformaldehyde in PBS for 1 minute at room temperature, rinsed in PBS, and incubated in a CO2 free incubator at 37°C for 12-16 hours in X-gal staining solution which was prepared fresh as described (104), after which sections were counterstained with eosin.

Percent Ki-67 positive cells in adenomas was quantitated using an automated algorithm in QuPath Version 0.3.0 (105). The analyzer was blinded to treatment group. Three high power fields per adenoma (image perimeter 2.5 mm) were selected from the most cellular portion with highest gland to stroma ratio. Total number of tumor cells and Ki-67 positive cells per high power field were quantitated using the analyze -> cell detection -> positive cell detection function with the following parameters: setup parameters (optical density sum, pixel size 0.5 μm), nucleus parameters (background radius 8 μm, median filter radius 0 μm, sigma 1.5 μm, min area 10 μm2, max area 400 μm2), intensity parameters (threshold 0.1, max background intensity 2, split by shape selected), cell parameters (cell expansion 5 μm, include cell nucleus selected), general parameters (smooth boundaries selected, make measurements selected), intensity threshold parameters (nucleus DAB OD mean, single threshold). Percent p21 positive cells in adenomas was quantitated in QuPath as described above with the following modifications. Three high power fields (image perimeter 2 mm) per adenoma were selected from the luminal region of the adenoma and three high power fields from the region adjacent to the muscularis propria (base). CD8+ and FOXP3+ cells per unit area were quantitated in QuPath as described above with the following modifications. Three high power fields per adenoma (image perimeter 2 mm) were selected from the most cellular portion with highest gland to stroma ratio. The same areas were used to quantitate CD8+ and FOXP3+ cells using the analyze -> cell detection -> positive cell detection function. For CD8 the score compartment was Cell DAB OD mean. For FOXP3 the score compartment was Nucleus DAB OD mean.

### Hematology and serum chemistry

Whole blood was collected as a terminal procedure. Mice were euthanized by CO2 inhalation. After confirming absence of respiration, venous blood was collected from the inferior vena cava using a 26G needle and 200 µL immediately transferred to an EDTA tube (Sarstedt, Item # 20.1288.100). Blood counts and hematology profiles were measured in the Division of Comparative Medicine at MIT on a HemaVet 950 FS (Drew Scientific) within a few hours of blood collection. To measure serum testosterone, whole blood collected as above was transferred to a serum separator tube (BD Biosciences, Cat # 365967), allowed to clot for 30 minutes at room temperature, and centrifuged for 90 seconds at 10,000 g. Serum was stored at −80°C until day of assay. Serum testosterone was measured by ELISA in 96-well plate format according to manufacturer recommendations (Cayman, Item # 582701). Absorbance (410 nm) was measured at 90 minutes on a Tecan infinite 200Pro. For standards, absorbance was plotted versus testosterone concentration and the curve was fitted by a 4-parameter logistic model, which was used to determine the testosterone concentration in serum samples. Calculations were performed in Prism version 8 (GraphPad Software).

### Flow Cytometry

Cells were stained using antibodies to CD4 (BD Biosciences Cat# 612761, RRID:AB_2870092), CD19 (BD Biosciences Cat# 550992, RRID:AB_398483), CD45 (BioLegend Cat# 103116, RRID:AB_312981), CD3 (BioLegend Cat# 100220, RRID:AB_1732057), CD8a (BioLegend Cat# 100759, RRID:AB_2563510), FOXP3 (BioLegend Cat# 126404, RRID:AB_1089117), CD25 (BioLegend Cat# 102012, RRID:AB_312861), CD19 (BioLegend Cat# 152409, RRID:AB_2629838), and NK1.1 (BioLegend Cat# 108705, RRID:AB_313392). All antibodies were diluted 1:100. Viability was assessed using Zombie Aqua (Biolegend, Cat# 423101) or Zombie UV (Biolegend, Cat# 423107) diluted 1:1000. Intracellular staining for FoxP3 was performed using the eBioscience FoxP3 Transcription Factor Buffer Set (ThermoFisher, Cat# 00-5523-00).

Blood, spleen, pelvic and axillary lymph nodes were harvested 2 days or 8 weeks after the last dose of radiation. All tissue samples were weighed and kept in RPMI media (ATCC, Cat# 30-2001) on ice during collection. Spleen and lymph nodes were mechanically digested through 70 um nylon cell strainers to prepare single-cell suspensions for staining. Red blood cells in spleen and blood samples were lysed in ACK Lysing Buffer (Gibco, Cat# A10492-01). All samples were resuspended in ice-cold PBS and stained for viability, then resuspended in ice-cold PBS containing 1% (w/v) BSA and 2 mM EDTA before labeling. Cells were analyzed using BD FACS LSR Fortessa (BD Biosciences), or BD FACS Symphony A3 (BD Biosciences) flow cytometers. Data was analyzed in FlowJo version 10 (BD Biosciences).

### Statistics

Continuous data were first assessed using the D’Agostino-Pearson normality test. Data that passed the normality test were then evaluated using two-tailed Student’s t test; independent data utilized unpaired tests, while dependent data utilized a paired test. Data that were not normally distributed were evaluated using the Mann-Whitney U test. Survival curves were compared using a log-rank test. The statistical test used and the level of statistical significance applied are described in each figure legend. In most figures, only P values <0.05 are noted. Unless otherwise indicated, data are presented as mean ± standard deviation. Prism Version 9 (GraphPad Software) was used for statistical analyses and data visualization.

### Study approval

Mouse studies were performed according to our animal protocol, which was approved by the MIT Institutional Animal Care and Use Committee.

## Supporting information

Supplementary File 1: Restrainer assembly and use instructions

Supplementary File 2: Restrainer A blueprint

Supplementary File 3: Restrainer A blueprint metric

Supplementary File 4: Restrainer B blueprint

Supplementary File 5: Restrainer B blueprint metric

Supplementary File 6: Side blocks blueprint

Supplementary File 7: Side blocks blueprint metric

Supplementary File 8: Shield and collimators blueprint

Supplementary File 9: Shield and collimators blueprint metric

Supplementary Video 1: This 24 second movie shows an approximately 5-minute fluoroscopic image time-lapse (12X speed) of a mouse in the restrainer.

## AUTHOR CONTRIBUTIONS

Designing research studies: DRS, JR, JDD, SRF, MVH

Conducting experiments: DRS, JR, IMTG, AS, CLW

Method Development: DRS, JDD, JR, AS, CLW, MRC, KC, CC, EH

Acquiring Data: DRS, JR, IMTG, AS, CLW, WH, HHM,

Analyzing Data: DRS, IMTG, AS, CLW, MH, WSK, MRC

Statistics review: LED

Supervision: MVH, MAS, OHY, KDW

Manuscript writing: DRS, MVH

Manuscript review & editing: JR, SRF, JDD, IG, AS

## ACKNOWLEDGMENTS

DRS was supported by DF/HCC SPORE in Prostate Cancer Training Award P50 CA090381-15, Harvard University KL2/Catalyst Medical Research Investigator Training award TR002542, and Joint Center for Radiation Therapy Foundation. MGVH acknowledges support from Ludwig Center at MIT, MIT Center for Precision Cancer Medicine, Stand up to Cancer, Emerald Foundation, Howard Hughes Medical Institute faculty scholar award, and NIH grants (P30CA1405141, R35CA242379, P50CA090381). JR acknowledges support from NIH grants (R37CA259363, R21CA256414, R21DK125911, R41EB032693, R01CA254108, R01CA256530, and R01CA244359), DOD grant W81XWH-20-1-0203, and Duke-NC State Translational Research Grant. Additionally, this work was supported by NCATS/NIH Award UL1 TR002541 to Harvard Catalyst and The Harvard Clinical and Translational Science Center and financial contributions from Harvard University and its affiliated academic health care centers. We thank Gerry Rondeau for constructing the lead shields, collimators, and restrainers; Xing-Qi Lu for assisting with dosimetry; Michael Brown and Alicia Caron for assisting with histology; Ellen Buckley for assisting with hematology; Dr. Pier Paolo Pandolfi for providing the *Pten;Trp53^pc-/-^* model and Dr. Sean Clohessy for helpful discussions regarding husbandry for this model. Graphical representations of the GEMM models and timelines were created with Biorender.

## List of Supplementary Materials

Figures S1 to S7

Supplementary File 1: Restrainer assembly and use instructions.

Supplementary File 2: Restrainer A blueprint.

Supplementary File 3: Restrainer A blueprint metric.

Supplementary File 4: Restrainer B blueprint.

Supplementary File 5: Restrainer B blueprint metric.

Supplementary File 6: Side blocks blueprint.

Supplementary File 7: Side blocks blueprint metric.

Supplementary File 8: Shield and collimators blueprint.

Supplementary File 9: Shield and collimators blueprint metric.

Supplementary Video 1: This 24 second movie shows an approximately 5-minute fluoroscopic image time-lapse (12X speed) of a mouse in the restrainer imaged by µCT. IV contrast was used to aid in visualization of kidneys and bladder.

