## Supplementary material for "Ablative radiotherapy improves survival in autochthonous cancer mouse models": Figures S1 to S7

### SUPPLEMENTARY FIGURES (Schmidt et al.)

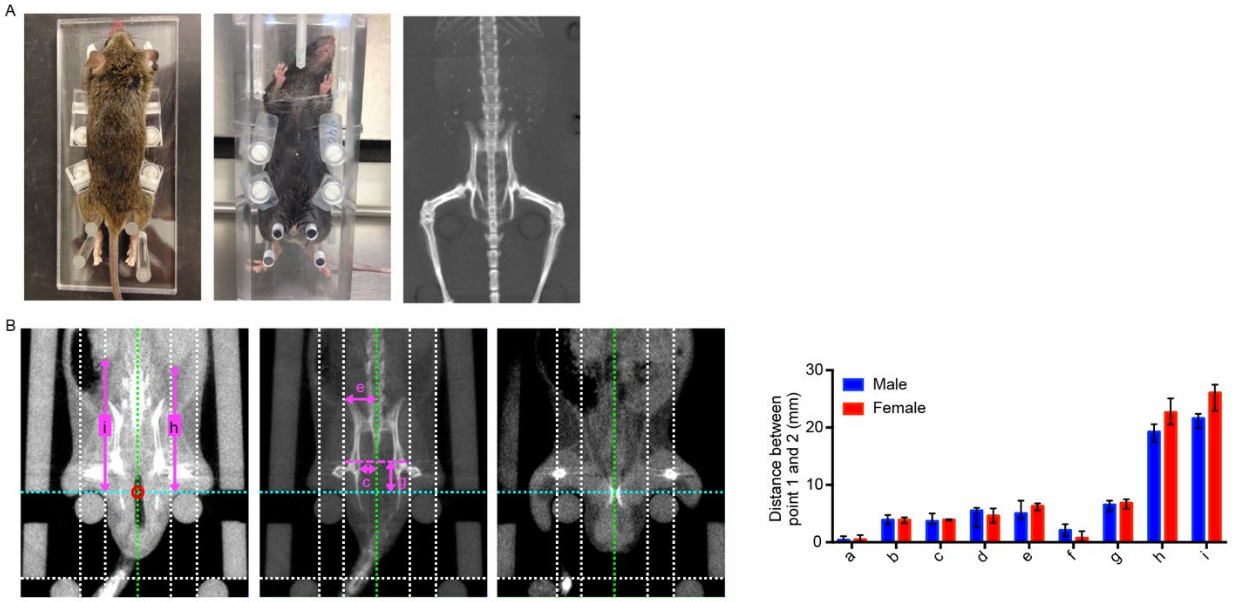


| | Point 1 | Point 2 | F 24-25g | F 26-28g | F 29-32g | F 33-34g | F 35-40g | F 41-47g | M 24-25g | M 26-28g | M 29-32g | M 33-34g | M 35-40g | M 41-47g |
|---|---|---|---|---|---|---|---|---|---|---|---|---|---|---|
| a | Pubic symphysis | Sagittal ref. plane | 0.0, 0.9 | 0.4, 1.2 | 0.3, 1.2 | 0.4, 0.6 | 0.3, 0.6 | 0.1, 0.4 | 0.0, 0.7 | 0.1, 1.1 | 0.2, 1.0 | 0.2, 0.3 | 0.1, 0.5 | 0.2, 0.3 |
| b | Medial edge of left fem. head | Sagittal ref. plane | 3.4, 3.8 | 3.6, 4.3 | 3.3, 4.2 | 3.1, 4.1 | 3.5, 4.8 | 3.8, 4.1 | 3.2, 3.8 | 3.1, 4.4 | 3.8, 4.7 | 4.0, 4.5 | 3.6, 4.2 | 4.3, 4.4 |
| c | Medial edge of right fem. head | Sagittal ref. plane | 3.7, 4.1 | 3.8, 4.0 | 3.8, 4.0 | 4.0, 5.0 | 3.9, 4.7 | 4.3, 4.9 | 2.8, 3.9 | 3.2, 5.0 | 3.3, 3.9 | 3.6, 3.8 | 4.2, 4.5 | 4.7, 4.8 |
| d | Lateral edge of left iliac crest | Sagittal ref. plane | 3.4, 3.7 | 3.4, 4.8 | 4.4, 5.9 | 4.4, 4.6 | 5.1, 6.0 | 4.7, 5.0 | 4.5, 6.5 | 2.7, 5.6 | 5.5, 6.0 | 6.3, 6.4 | 5.1, 5.6 | 5.1, 5.7 |
| e | Lateral edge of right iliac crest | Sagittal ref. plane | 6.2, 7.1 | 6.1, 7.0 | 5.5, 6.5 | 5.8, 6.6 | 5.1, 5.4 | 6.0, 6.2 | 3.7, 5.2 | 4.1, 7.2 | 4.7, 5.2 | 4.1, 4.2 | 5.1, 6.0 | 6.1, 6.0 |
| f | Top of pubic symphysis | Axial ref. plane | 0.4, 2.6 | 0.4, 1.9 | 0.1, 1.1 | 0.4, 1.2 | 0.1, 1.6 | 1.1, 1.9 | 0.1, 0.4 | 1.0, 3.1 | 2.0, 2.6 | 2.1, 2.9 | 2.4, 3.6 | 2.1, 4.1 |
| g | Upper edge of fem. heads | Axial ref. plane | 4.8, 6.0 | 6.4, 7.1 | 5.8, 7.5 | 6.7, 7.3 | 6.2, 7.3 | 8.1, 8.4 | 4.8, 5.7 | 5.3, 7.1 | 6.3, 7.2 | 6.7, 7.1 | 6.7, 8.0 | 7.7, 9.5 |
| h | Inferior pole of left kidney | Axial ref. plane | 19.7, 21.5 | 20.5, 25.1 | 21.8, 24.2 | 22.5, 27.1 | 23.3, 24.3 | 24.4, 25.3 | 18.7, 20.2 | 17.5, 20.6 | 19.0, 20.1 | 21.1, 23.4 | 22.7, 24.1 | 20.6, 22.1 |
| i | Inferior pole of right kidney | Axial ref. plane | 22.4, 23.9 | 22.9, 27.1 | 25.9, 27.1 | 25.7, 28.0 | 25.4, 27.9 | 27.2, 28.9 | 19.6, 21.9 | 19.9, 22.4 | 20.7, 22.4 | 23.0, 25.1 | 23.3, 25.6 | 24.3, 25.1 |

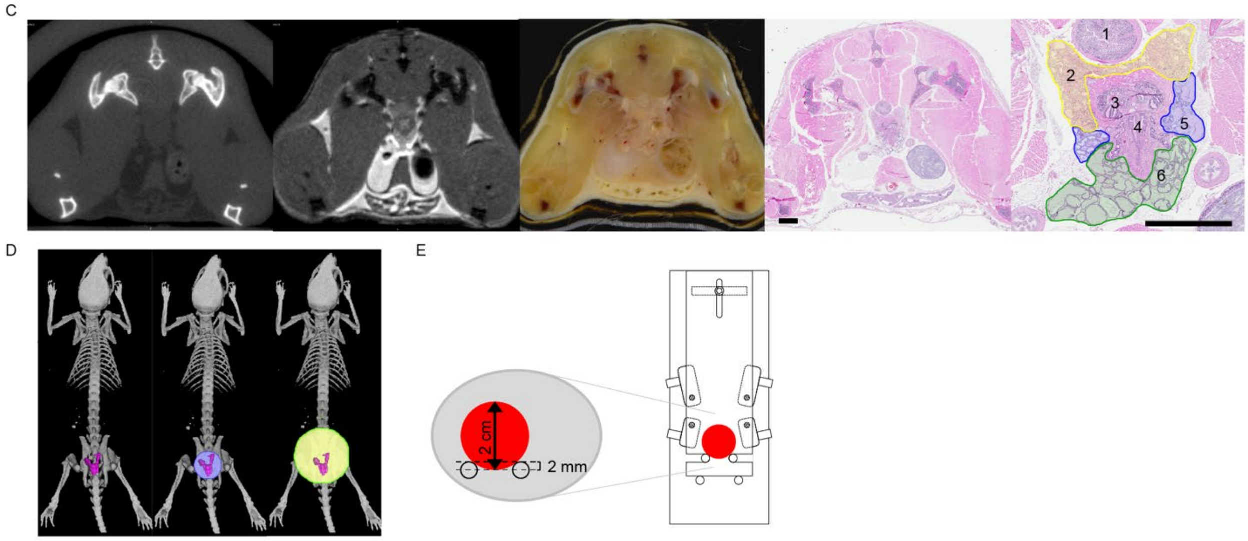


Supplementary Figure 1

F

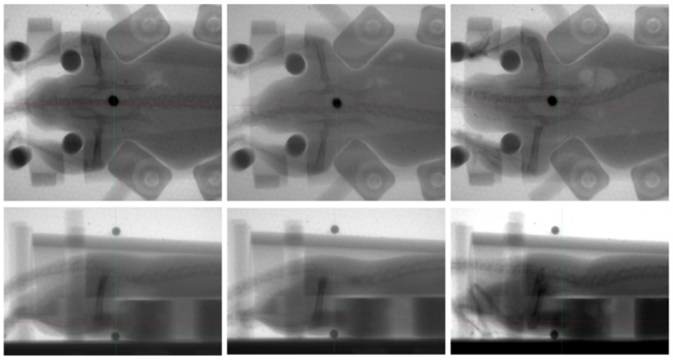

**Supplementary Figure 1. Design and validation of a mouse restrainer for SABR delivery.** (**A**) Initial prototype of restrainer and MIP CT image of the lower torso and hind limbs showing optimal positioning of the pelvis and immobilization of hind limbs. (**B**) Quantitative assessment of reproducible animal positioning relative to reference points on the restrainer. Representative images show the anatomic and reference landmarks evaluated. Letters correspond to measured distances between points described in the table. The sagittal reference plane (shown in green) is defined as a straight line midpoint between the medial edge of the upper and lower pins. The axial reference plane (shown in cyan) is defined as a straight line connecting the upper edge of the upper pins. The central axis of the field (red circle in left panel) is defined as the intersection of the sagittal and axial reference planes. Values show in the table are the minimum and maximum distances (in mm) observed between point 1 and 2. The top row indicates the sex (F, female; M, male) and body weight range (g, grams). To know whether results would be generalizable to mice of different strains, we used mixed-strain (C57Bl/6J; 129/SvJ), inbred (C57Bl/6J), and outbred (CD-1) strains for this analysis. All mice were age 3 months or older. Shown in the graph are distances (median and range) between point 1 and 2 as described in the table for mice weighing 26 to 32 grams (n=8 for male, n=6 for female). (**C**) In situ anatomy of the prostate shown on formalin-fixed paraffin-embedded tissue sections of a mouse previously evaluated in the same anatomic position by CT, MRI and wet mount cross-sectional imaging. 1, rectum; 2, dorsal prostate; 3, ducts from anterior prostate and seminal vesicles (both of which are located cranial to the axial slice shown); 4, urethra; 5, lateral prostate; 6, ventral prostate. Scale bar = 2 mm. (**D**) Location of the prostate relative to bony anatomic landmarks (see text for details). Relationship of the prostate to 1 and 2 cm diameter pelvic radiation fields is shown. (**E**) Optimal alignment of the restrainer with the collimator aperture in order to position the prostate within a 2 cm radiation field as shown in (D). (**F**) Fluoroscopic images (top and side view) of three male mice immobilized in the restrainer. The radio-opaque markers on the restrainer were used to indicate the center of the radiation field shown in (E). Note that the bony pelvis is accurately aligned to place the geometric center of the prostate at the center of the beam.

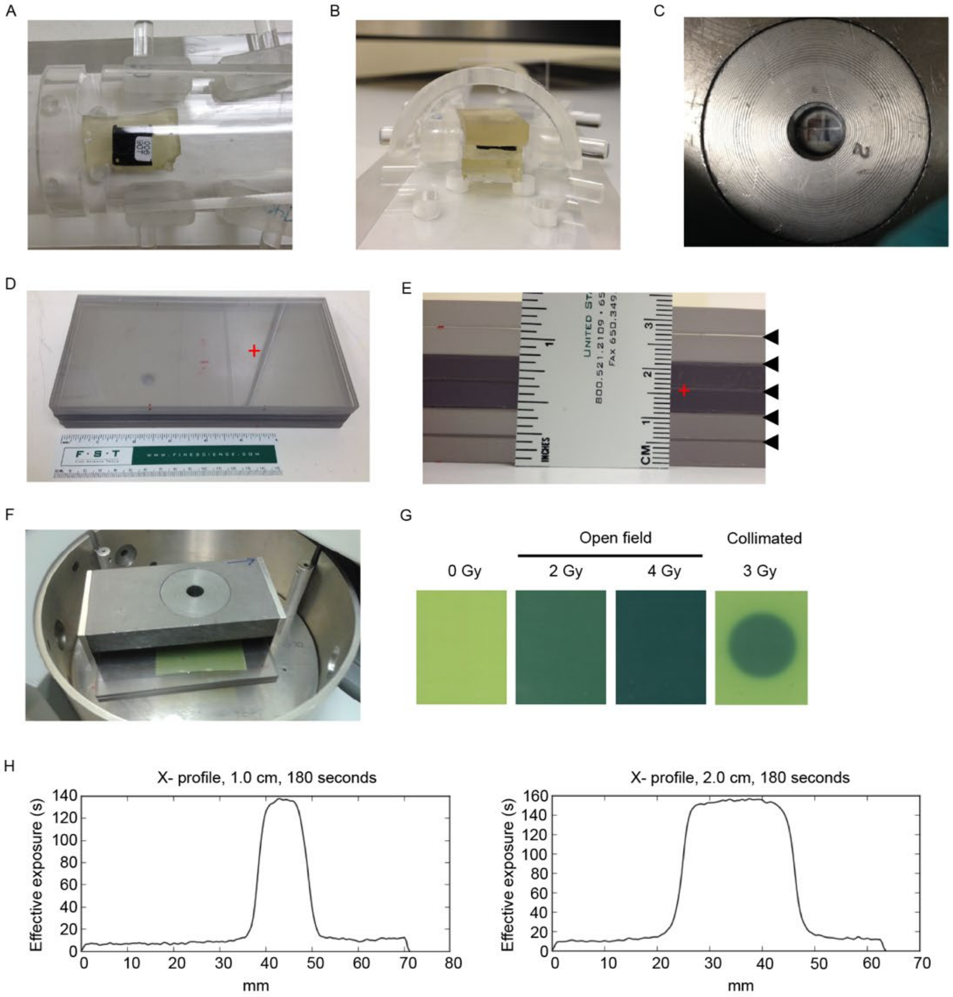


I

| | | |
|---|---|---|
| Collimator aperture | 1.00 cm | 2.00 cm |
| Measured diameter (50% intensity) | 1.14 cm | 2.21 cm |
| Measured penumbra (20-80% intensity) | 0.25 cm | 0.31 cm |
| Estimated output factor | 0.62 | 0.72 |

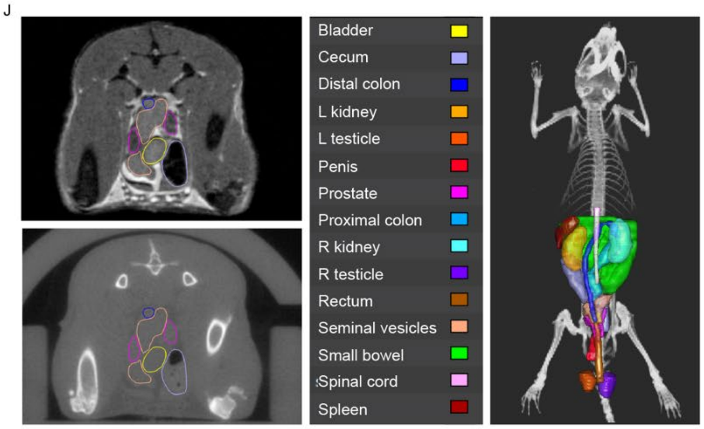


Supplementary Figure 2

K

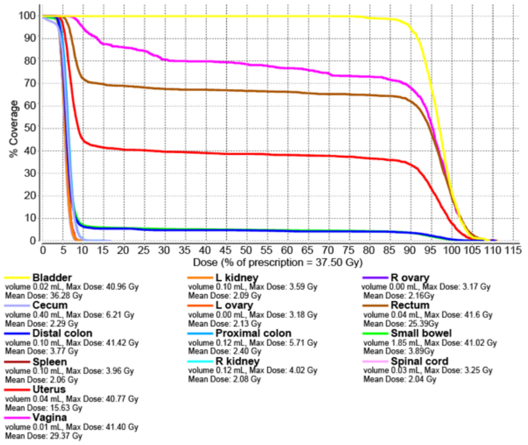


L

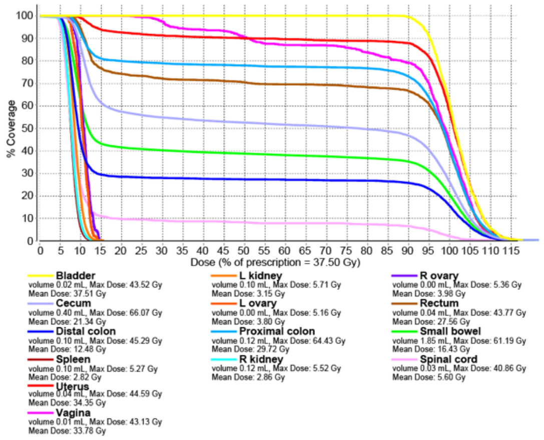


M

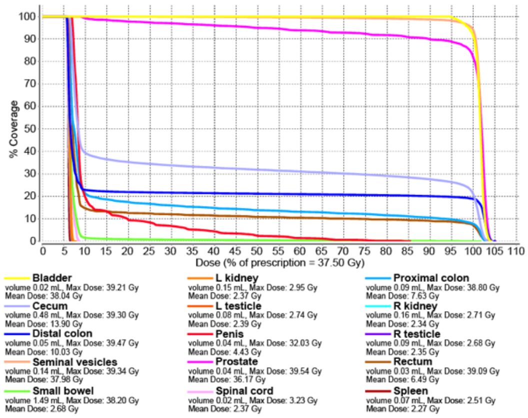


N

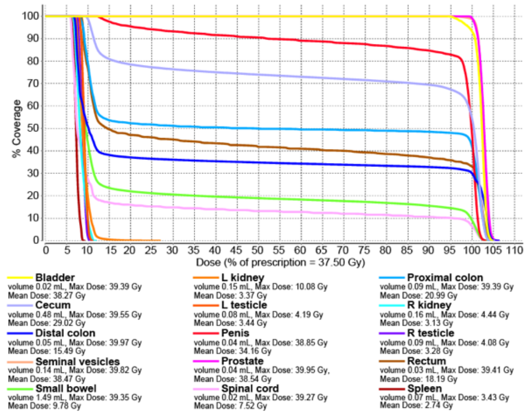


**Supplementary Figure 2. Dosimetry and modeling of radiation dose distribution in mice.** (**A**) Top view of Nanodot dosimeter exposure setup. The upper bolus slab is omitted for visualization of the dosimeter. (**B**) Side view of dosimeter exposure setup. (**C**) Top view of dosimeter in collimated field. The upper bolus slab is omitted for visualization of the dosimeter. (**D**) Top view of radiochromic film exposure setup. The red cross indicates the location at which film was centered to measure the horizontal dose distribution. (**E**) Side view of radiochromic film exposure setup. The red cross indicates the location at which film was centered to measure the horizontal dose distribution. The black arrowheads indicate the level at which film was placed to determine the dose distribution in the perpendicular plane. (**F**) Placement of radiochromic film in the collimated field with upper stack of plexiglass sheets removed for visualization of film. (**G**) Representative results of radiochromic film exposed in an open field at 2 dose levels and a 2 cm collimated field. (**H**) Representative plots of the horizontal dose distribution for circular collimated fields with 1 and 2 cm apertures evaluated at mid-plane of the radiation field by radiochromic film dosimetry. (**I**) Collimated field parameters evaluated by radiochromic film dosimetry. The average of two independent tests is shown. (**J**) Representative MRI and µCT images demonstrating how rigid image registration was used to transfer abdominopelvic organs contoured on MRI (upper left) to μCT (lower left). The panel at right shows a dorsal view 3D rendering of abdominopelvic organs of a male mouse defined by MRI onto the bony skeleton imaged by μCT. (**K**) Dose-volume histogram for a 1 cm pelvic radiation field in a female mouse. (**L**) Dose-volume histogram for a 2 cm pelvic radiation field in a female mouse. (**M**) Dose-volume histogram for a 1 cm pelvic radiation field in a male mouse. (**N**) Dose-volume histogram for a 2 cm pelvic radiation field in a male mouse.

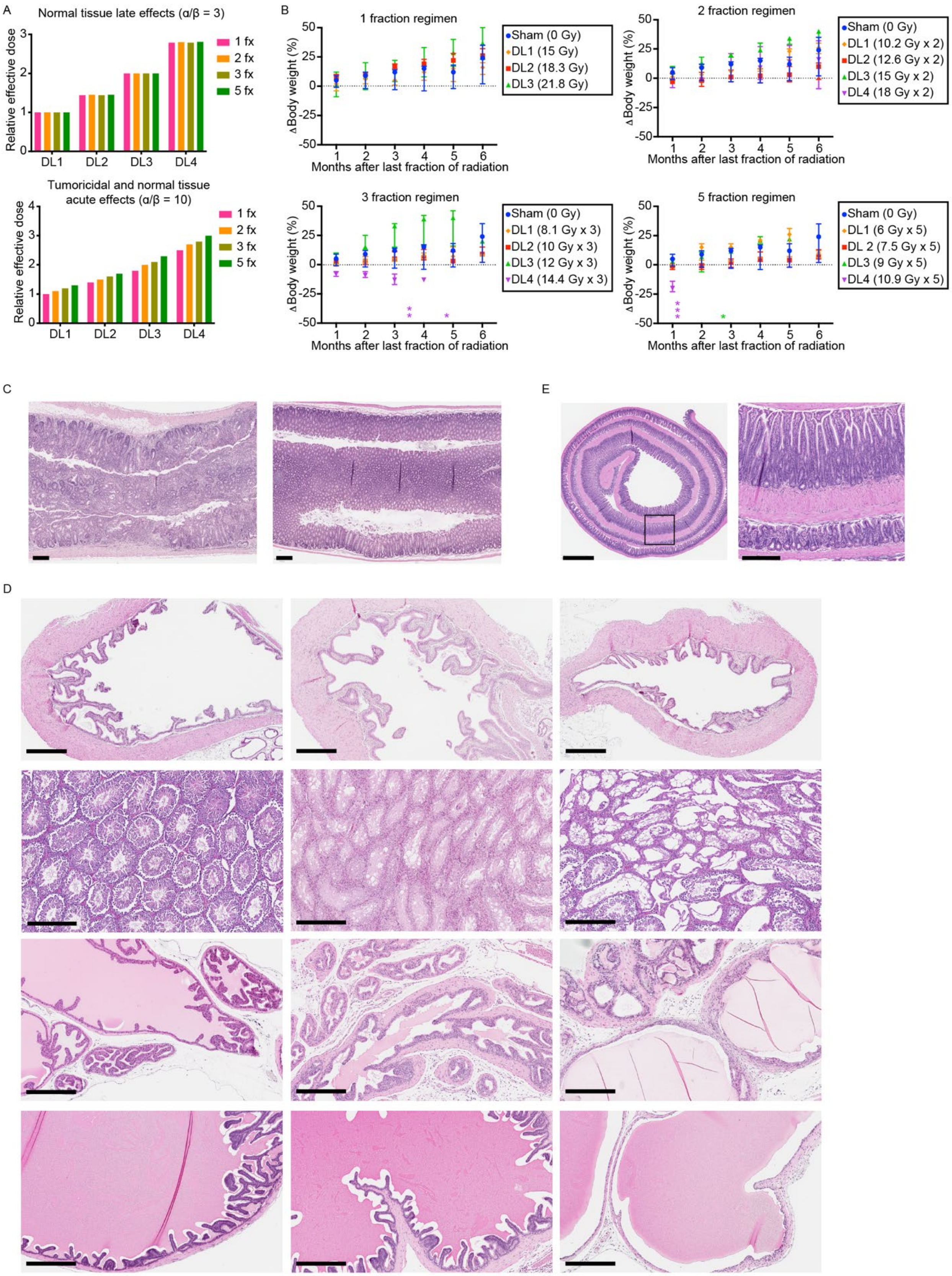
A
Normal tissue late effects (α/β = 3)
Relative effective dose
1 fx
2 fx
3 fx
5 fx
DL1
DL2
DL3
DL4
Tumoricidal and normal tissue acute effects (α/β = 10)
B
1 fraction regimen
Sham (0 Gy)
DL1 (15 Gy)
DL2 (18.3 Gy)
DL3 (21.8 Gy)
2 fraction regimen
Sham (0 Gy)
DL1 (10.2 Gy x 2)
DL2 (12.6 Gy x 2)
DL3 (15 Gy x 2)
DL4 (18 Gy x 2)
3 fraction regimen
Sham (0 Gy)
DL1 (8.1 Gy x 3)
DL2 (10 Gy x 3)
DL3 (12 Gy x 3)
DL4 (14.4 Gy x 3)
5 fraction regimen
Sham (0 Gy)
DL1 (6 Gy x 5)
DL 2 (7.5 Gy x 5)
DL3 (9 Gy x 5)
DL4 (10.9 Gy x 5)
ΔBody weight (%)
Months after last fraction of radiation
C
E
D

Supplementary Figure 3

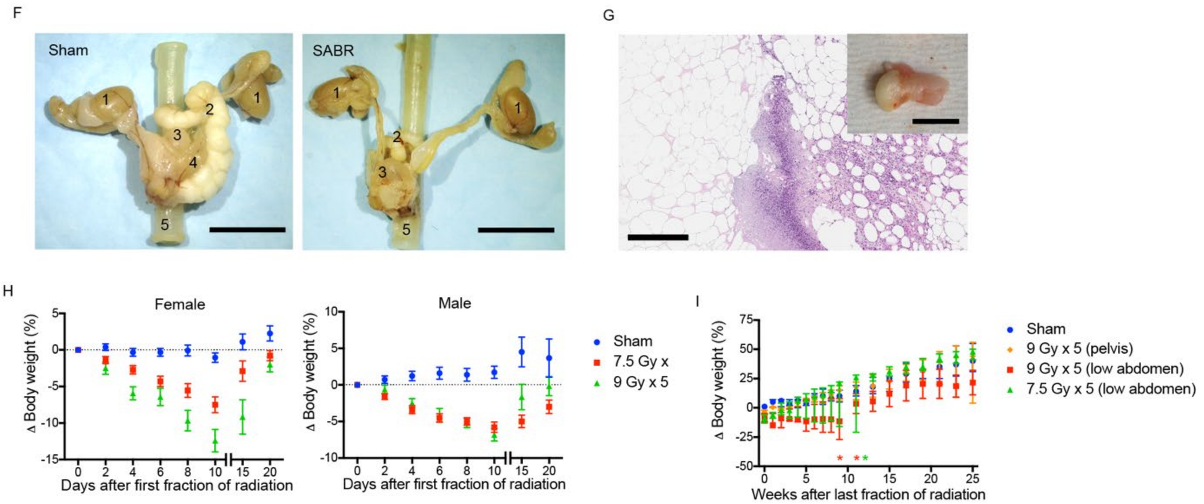


J

| Sex | Regimen | 2 days | 7 days | 14 days | 56 days |
|---|---|---|---|---|---|
| M | 7.5 Gy x 5 | 0 of 6 | 0 of 3 | 0 of 3 | 0 of 6 |
| | 9 Gy x 5 | 0 of 12 | 0 of 3 | 0 of 3 | 0 of 6 |
| F | 6 Gy x 5 | 0 of 3 | | | 0 of 3 |
| | 7.5 Gy x 5 | 3 of 12 | 2 of 3 | 2 of 6 | 1 of 6 |
| | 9 Gy x 5 | 3 of 3 | 3 of 3 | 2 of 3 | 3 of 6 |

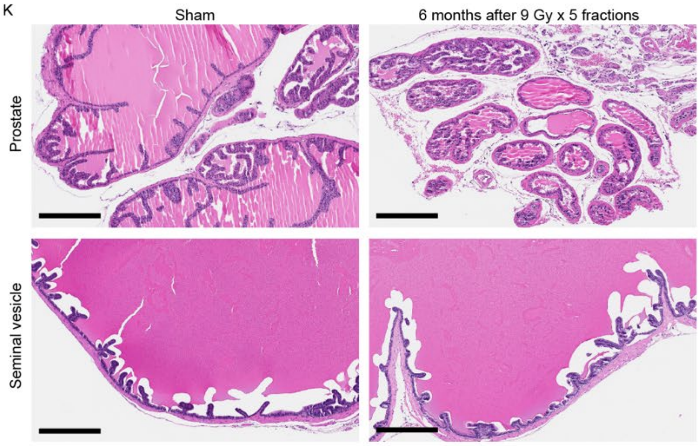


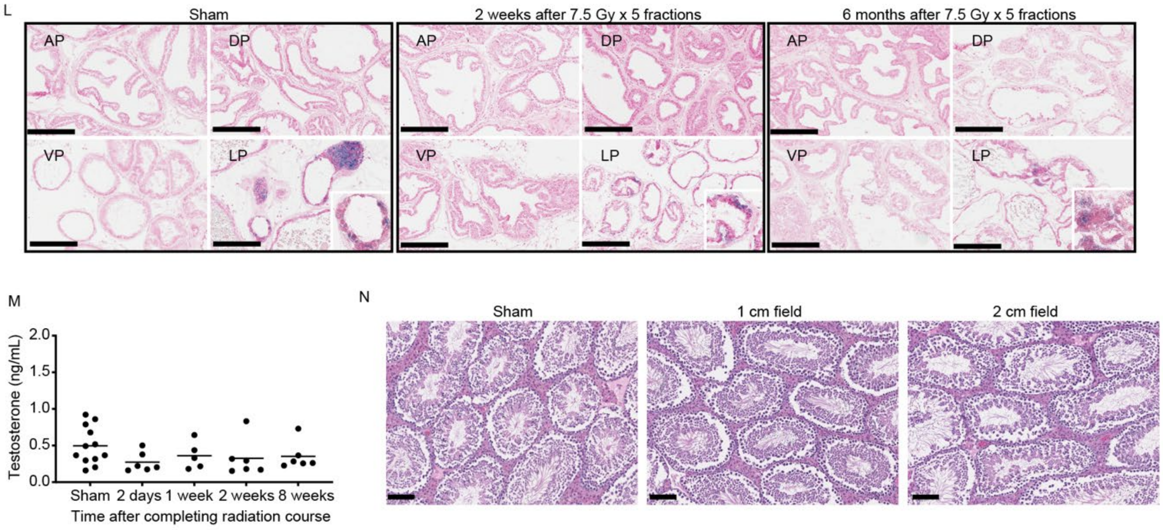


Supplementary Figure 3 (continued)

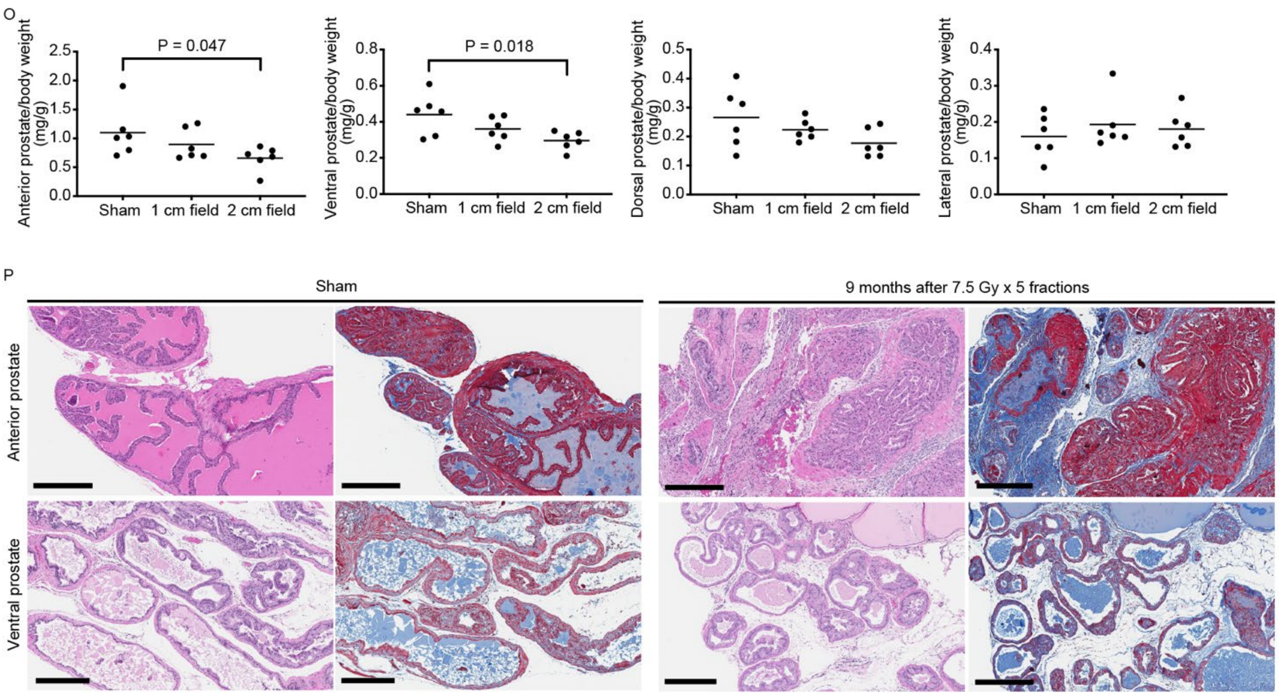


**Supplementary Figure 3. Pelvic SABR tolerance dose and effects on tissues.** (**A**) Graphical representation of the relative biologic effective dose (BED) for the dose levels shown in table 1. BED was calculated using alpha-beta ratio = 3 for late effects and alpha-beta ratio = 10 for acute effects and plotted relative to the BED for a single fraction at DL1. (**B**) Body weight change relative to pretreatment baseline. Median +/- 95% CI. All subjects are male. n=5 for 0 Gy, n=3 for all others (except n=2 for 21.8 Gy single fraction group, as one subject was removed from study due to injury unrelated to treatment). Each asterisk (*) indicates time of death/euthanasia of a single subject. (**C**) Representative histology of distal colon inside the radiation field ("in-field") and adjacent colon outside the radiation field ("out-of-field") 5 weeks after treatment with 5 fractions of 10.9 Gy. (**D**) Representative histology of GU tissue 5 weeks after 5 fractions of 10.9 Gy or 6 months after 2 fractions of 18 Gy to a 2 cm pelvic radiation field. GU tissue from a sham treated control is shown for comparison. Scale bar = 600 μm for bladder and 300 μm for all others. (**E**) Swiss roll of ilium with distal end at center showing hyperplasia of the mucosa and muscularis propria 11 weeks after completing 5 fractions of 9 Gy to a 2 cm pelvic radiation field. Corresponds to the specimen in Fig. 3F. Scale bar = 1 mm for the low power view and 300 μm for the high power view. (**F**) Gross pathology of genitourinary organs and adjacent rectum of male mouse 6 months after treatment with 18 Gy x 2 fractions to a 2 cm pelvic radiation field centered on the prostate. Sham treated control is shown for comparison. 1: testis, 2: seminal vesicles, 3: bladder, 4: anterior prostate, 5: rectum. Note: anterior prostate is small and obscured by the bladder in the SABR treated specimen. Scale bar 3 mm. (**G**) Histology of epididymal fat pad showing region of fibrosis and necrosis adjacent to normal fatty tissue 6 months after treatment with 18 Gy x 2 fractions to a 2 cm pelvic radiation field. Scale bar = 300 μm. Gross pathology is shown in the inset (scale bar = 5 mm). (**H**) Body weight change relative to pretreatment baseline during the first 3 weeks of treatment. Mean +/- SEM. Females n=12 for days 2–10, n=6 for days 15 and 20. Males n=15 for days 2–10, n=6 for days 15 and 20. (**I**) Body weight change relative to pretreatment baseline for female mice treated with a 1 cm radiation field targeted to the pelvis or low abdomen. Median +/- 95% CI. n=5 for sham group, n=4 for all treatment groups. Each asterisk (*) indicates time of death/euthanasia of a single subject. (**J**) Incidence of acute small bowel toxicity at the indicated time points after completing radiation. Toxicity was defined as any abnormal appearance of the intestine or luminal contents determined by visual inspection at time of necropsy. Segments of the intestine that appeared abnormal also underwent histologic evaluation. (**K**) Representative histology of anterior prostate and seminal vesicle 6 months after 5 fractions of 9 Gy to a 2 cm pelvic radiation field. Sham treated control is shown for comparison. Scale bar = 300 μm. (**L**) Representative SA-βgal staining of prostate tissue. AP, anterior prostate; DP, dorsal prostate; VP, ventral prostate; LP, lateral prostate. Scale bar = 200 μm. Irrespective of treatment group, some LP glands had βgal positive cells as shown at high power magnification in the inset. (**M**) Serum testosterone in C57Bl/6 male mice at the indicated time points after treatment with 5 fractions of 7.5 or 9 Gy to a 2 cm pelvic radiation field centered on the prostate. Sham treated control is shown for comparison. (**N**) Representative histology of testis from CD-1 mice 9 months after sham radiation or 5 fractions of 7.5 Gy to a 1 or 2 cm radiation field centered over the prostate. Scale bar = 100 μm. (**O**) Wet weight of prostate in male mice 9 months after sham radiation or 5 fractions of 7.5 Gy to a 1 or 2 cm pelvic radiation field. P value estimate by unpaired, two-tailed t test. (**P**) Representative H&E and trichrome stain showing fibrosis (blue) between glands in the anterior and ventral prostate of CD-1 male mice 9 months after sham treatment or 5 fractions of 7.5 Gy to a 2 cm pelvic field. Note that secretions in the prostate lumen also stain blue. Scale bar = 300 μm.

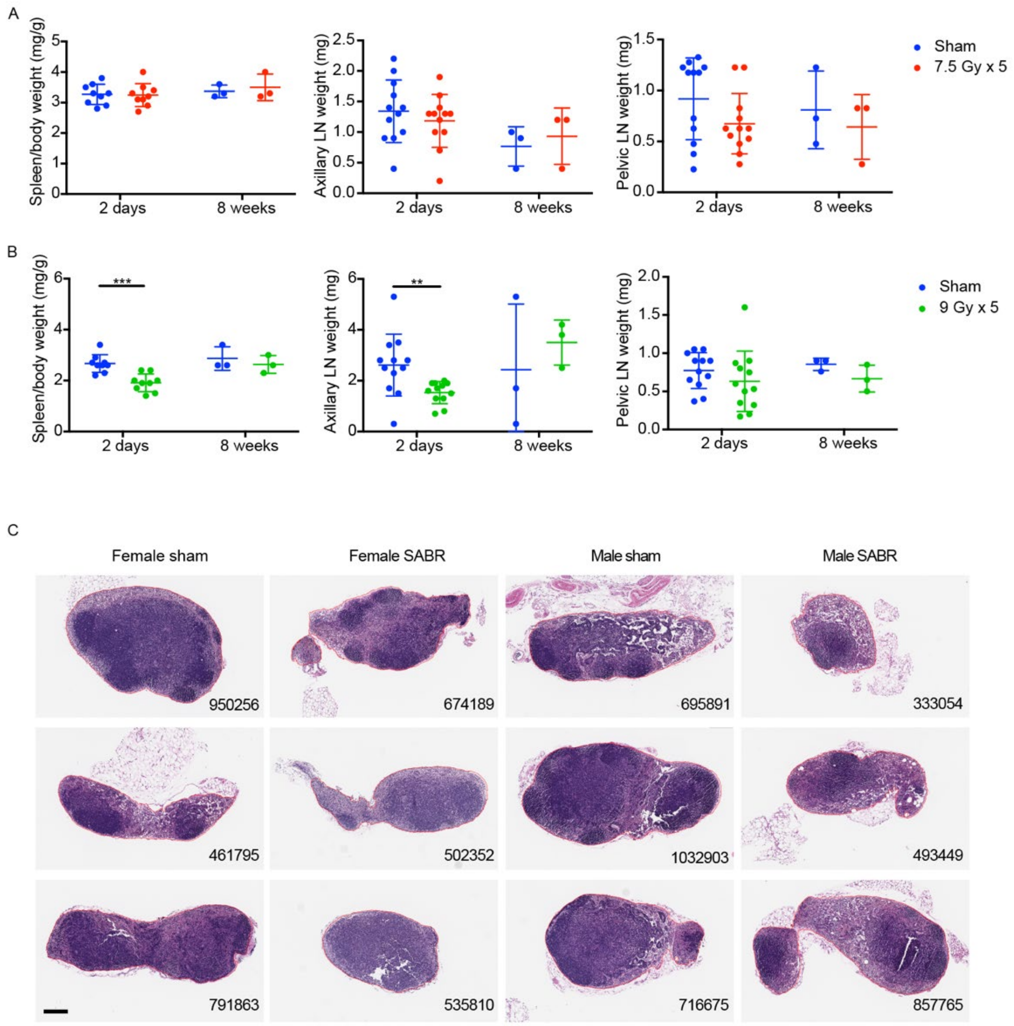


**Supplementary Figure 4. Pelvic SABR effects on spleen and lymph nodes.** (**A**) Weight of whole spleen, axillary lymph node, and pelvic lymph node tissue from female mice. Wet weight was determined at time of necropsy. n=9 for spleen and n=12 for lymph nodes at 2-day time point, n=3 for spleen and lymph nodes at 8-week time point. Mean +/- SD is shown. (**B**) Weight of whole spleen, axillary lymph node, and pelvic lymph node tissue from male mice. Wet weight was determined at time of necropsy. n=9 for spleen and n=12 for lymph nodes at 2-day time point, n=3 for spleen and lymph nodes at 8-week time point. Mean +/- SD is shown. **p<0.01, ***p<0.001 by unpaired 2-tailed t test. (**C**) Cross-section at midplane of pelvic lymph nodes (n=3) 2 days after sham treatment or 5 fractions of 7.5 Gy (female) or 9 Gy (male). Number at bottom right indicates the area ($\mu m^2$) within the red outline. All images are the same magnification. Scale bar = 200 μm.

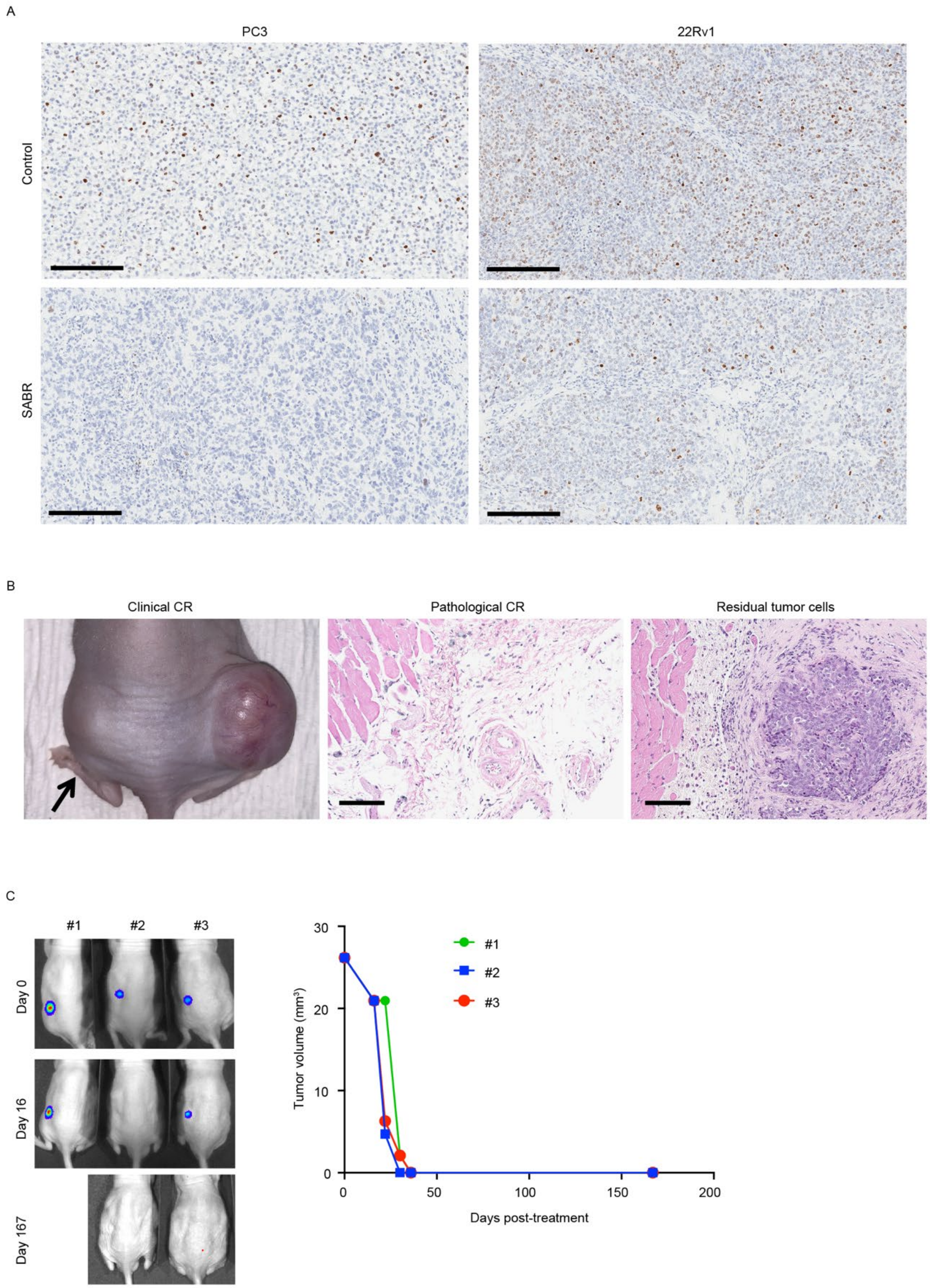


**Supplementary Figure 5. Clinical and pathological response of flank xenografts to SABR.** (**A**) Representative Ki67 IHC of PC3 and 22Rv1 xenografts one month after 4 fractions of 6 Gy. Untreated control is shown for comparison. Scale bar = 200 μm. (**B**)

Representative examples of clinical and pathological complete responses 4 weeks after completing SABR. Clinical complete response was defined as no visible or palpable tumor in the irradiated flank (black arrow). Pathological complete response was defined as no identifiable tumor cells on two step sections through the tumor scar. A SABR-treated 22Rv1 xenograft with residual tumor cells is shown for comparison. Scale bar = 100 µm. (**C**) Three nude mice bearing a single PC3-luc xenograft measuring at least 25 $mm^3$ were treated with 5 fractions of 7.5 Gy. Bioluminescence images obtained prior to fraction 1 and on day 16 and 167 thereafter. One animal was moribund on day 138. Necropsy showed no tumors and no evidence of radiation toxicity. Graph shows tumor volume based on caliper measurements.

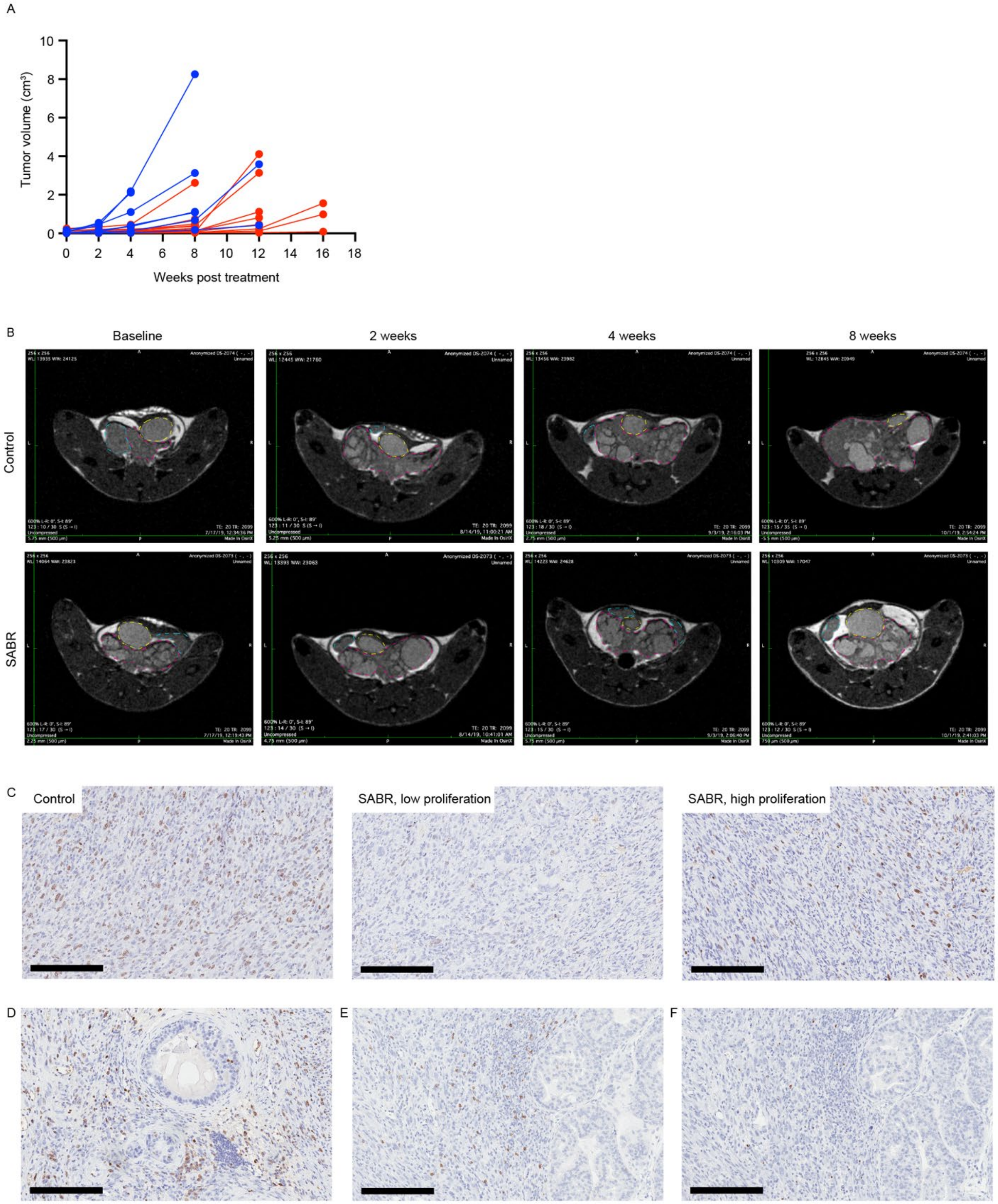


**Supplementary Figure 6. Radiographic and histopathologic response of prostate cancer to SABR.** (**A**) Tumor progression in study 1 as defined in Fig 6D assessed by serial MRI. Individual animal absolute volume data are shown. n=7 for sham (blue), n=14 for SABR (red). (**B**) Representative axial slice of pelvic MRI at 4 months of age (baseline), and repeat MRI at 2, 4, and 8 weeks after treatment. Areas outlined are bladder (yellow), seminal vesicle (blue), prostate tumor (pink). (**C**) Representative Ki67 IHC in an untreated tumor and two regions of a tumor 10 weeks post treatment with 5 fractions of 9 Gy. Scale bar = 200 µm. (**D**) Ki67 IHC in region of tumor with glands showing intraepithelial neoplasia and adjacent lymphocytic infiltrates. Scale bar = 200 µm. (**E**) Representative CD8 IHC in an untreated tumor. Scale bar = 200 µm. (**F**) Representative FOXP3 IHC in an untreated tumor. Scale bar = 200 µm.

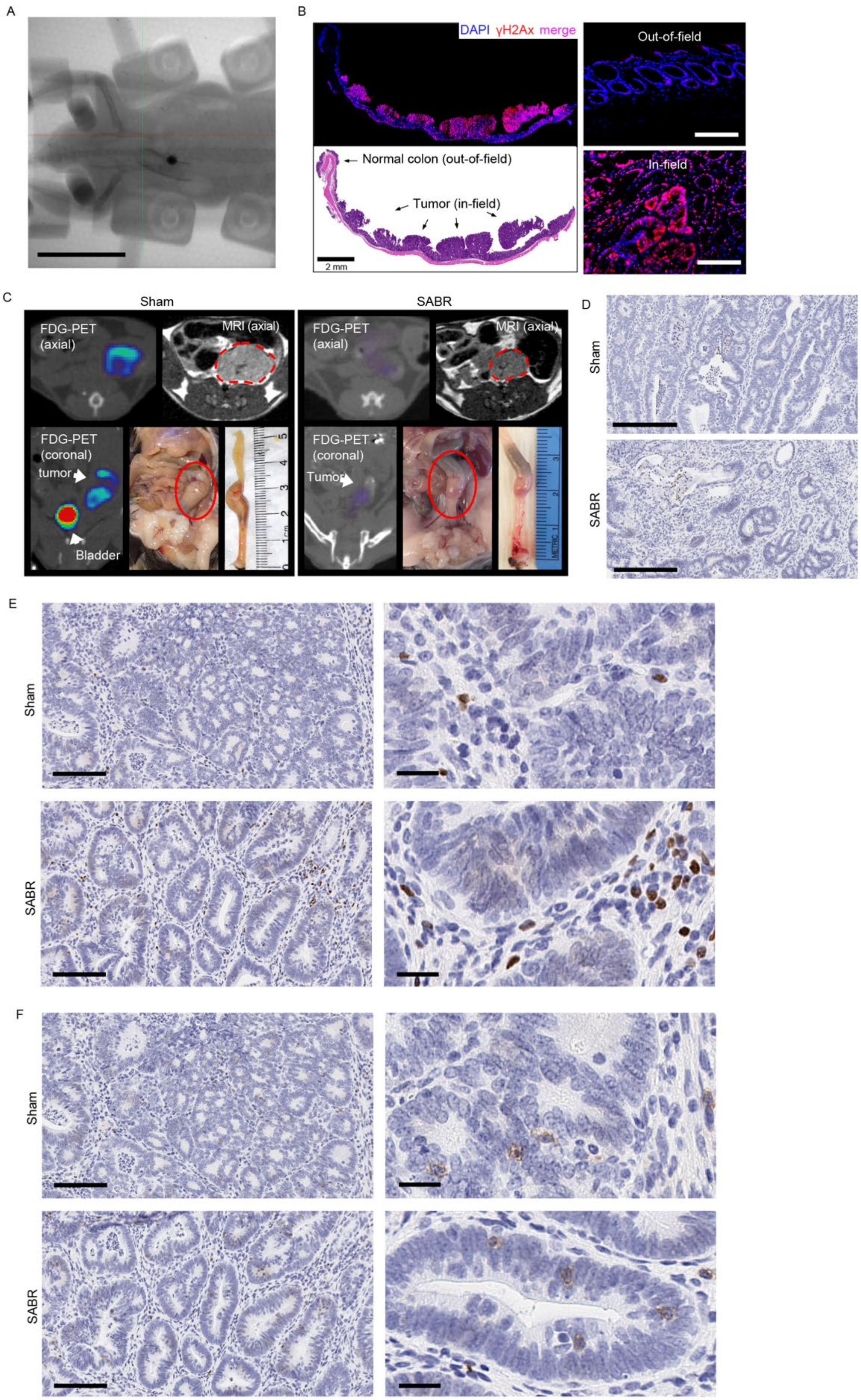


Supplementary Figure 7

**Supplementary Fig. 7. Evaluating SABR effects in a mouse colorectal adenoma model.** (**A**) Under endoscopic guidance 200 μL radiopaque contrast was injected into the submucosa 2-3 cm from the anal verge to identify the region of rectum at risk for tumor induction. The animal was then imaged in the restrainer with a radiopaque marker on the restrainer at center of the treatment field. The representative fluoroscopic image shows good placement of the rectal tumor target region relative to the treatment isocenter. Scale bar = 2 cm. (**B**) Evaluation of focal DNA damage in a longitudinal cross-section of colorectum by $\gamma$-H2AX staining. This mouse with multifocal adenoma was treated with 7.5 Gy to a 2 cm diameter radiation field 30 minutes prior to tissue collection. Nuclear $\gamma$-H2AX foci are seen only in the portion of the colorectum that was within the radiation field. Uniform DNA damage induction is seen throughout the entire tumor and small margin of adjacent normal tissue. Scale bar = 2 mm for the low power images and 0.1 mm for the high-power views. (**C**) FDG-PET, MRI, and gross pathology of a representative SABR-treated animal and untreated control. The right panel shows FDG-uptake and tumor size 6 months after treatment with 5 fractions of 7.5 Gy. The left panel shows an untreated control for comparison. (**D**) Representative cleaved caspase-3 IHC images of tumors from study 2 as defined in Fig. 7A. Scale bar = 200 μm. (**E**) Representative CD8 IHC images of tumors from study 2 as defined in Fig. 7A. Scale bar = 100 μm (low power view), 20 μm (high power view). (**F**) Representative FOXP3 IHC images of tumors from study 2 as defined in Fig. 7A. Scale bar = 100 μm (low-power view), 20 μm (high-power view).
