## Supplementary File 1: Restrainer assembly and use instructions for "Ablative radiotherapy improves survival in autochthonous cancer mouse models"

Step 2: Anterior set of hind limb pins are inserted medial to femur to prevent animal from exiting the chamber.

Step 4: Posterior set of hind limb pins are inserted lateral to the ankle. (note: the knee joint is now secured)

Step 6: Sliding nose piece is adjusted to mouse length and secured by tightening thumb screw. (note: animal is now secured)

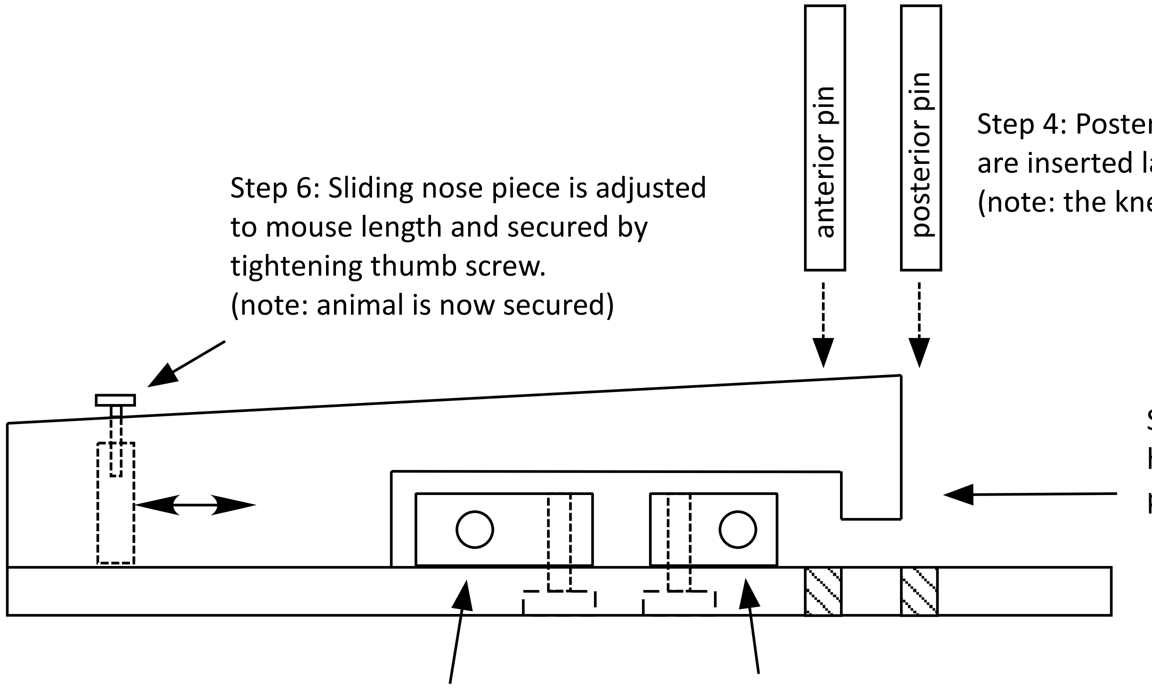


Step 1: Animal enters chamber head-first from rear of holder in prone ("belly down") position.

Step 5: Thorax blocks are adjusted to secure forelimbs into a forward ("arms up") position.

Step 3: Hind limbs are brought into caudal ("legs extended") position and secured by adjusting abdominopelvic blocks. (note: the femur is now secured)

(note: thorax and abdominopelvic blocks are secured by tightening nylon screws on underside of holder)

### Restrainer assembly and use instructions

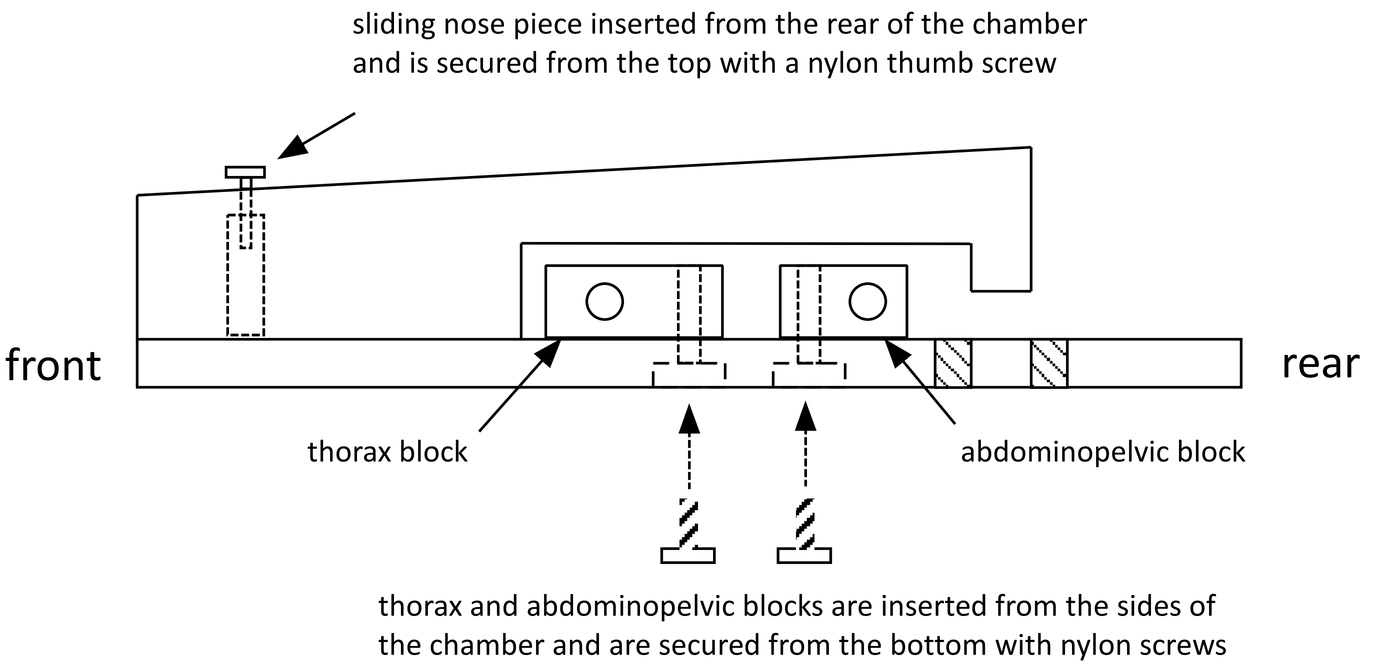


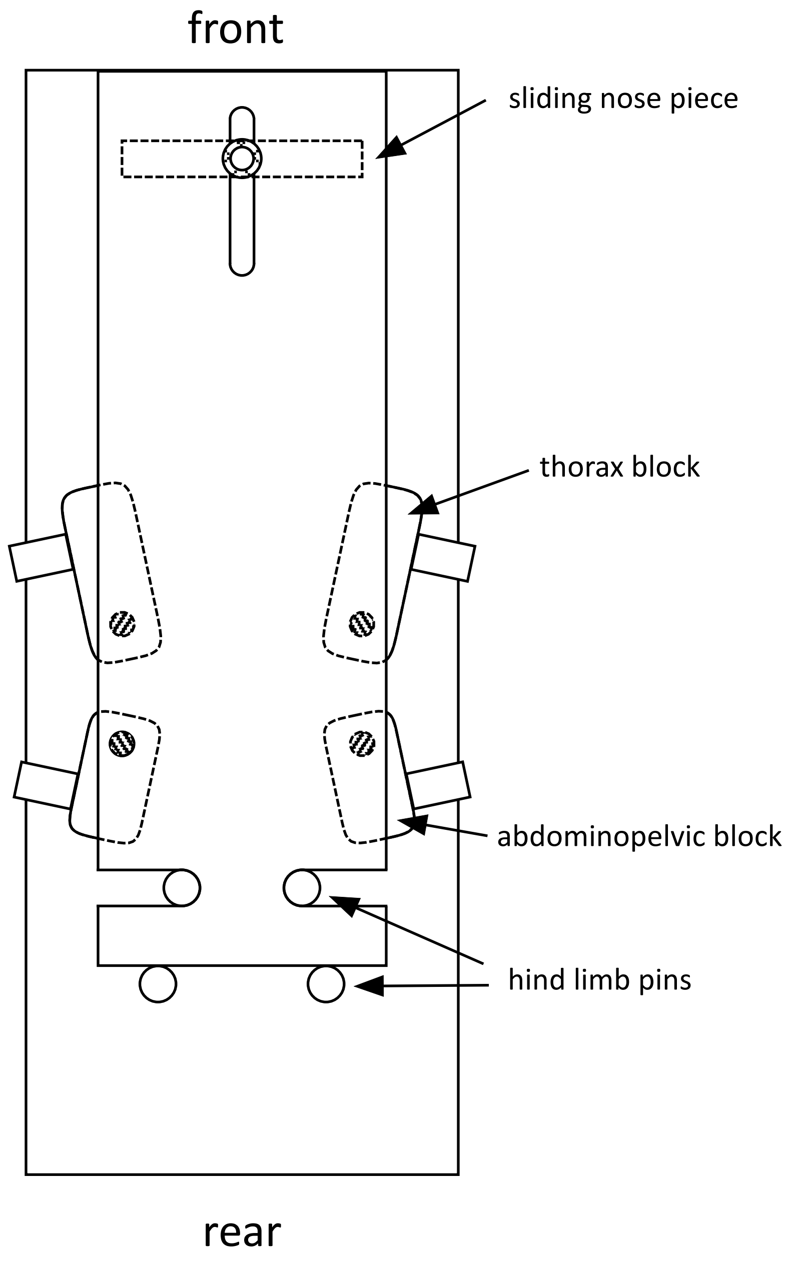


table of recommended block size by mouse weight

| mouse weight (grams) | thoracic block length (inches) | thoracic block length (mm) | abdominopelvic block length (inches) | abdominopelvic block length (mm) |
|---|---|---|---|---|
| 24 - 25 | 0.760 | 19.3 | 0.740 | 18.8 |
| 26 - 28 | 0.760 | 19.3 | 0.660 | 16.8 |
| 29 - 32 | 0.840 | 21.3 | 0.660 | 16.8 |
| 33 - 40 | 0.840 | 21.3 | 0.580 | 14.7 |
| 41 - 47 | 0.920 | 23.4 | 0.580 | 14.7 |
