## Supplementary File 2: Restrainer A blueprint for "Ablative radiotherapy improves survival in autochthonous cancer mouse models"

### Restrainer for 24-34 g mouse
(drawn to scale)

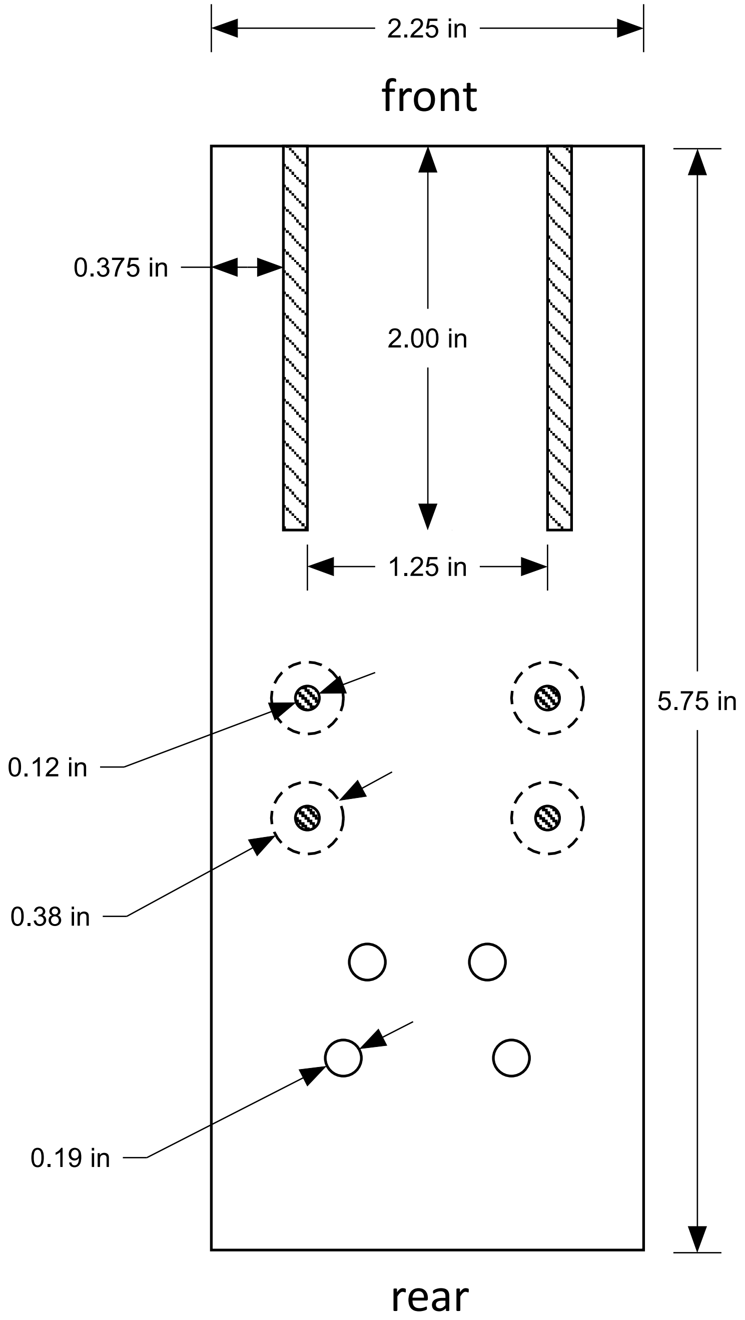


bottom view

### Restrainer for 24-34 g mouse
### (drawn to scale)

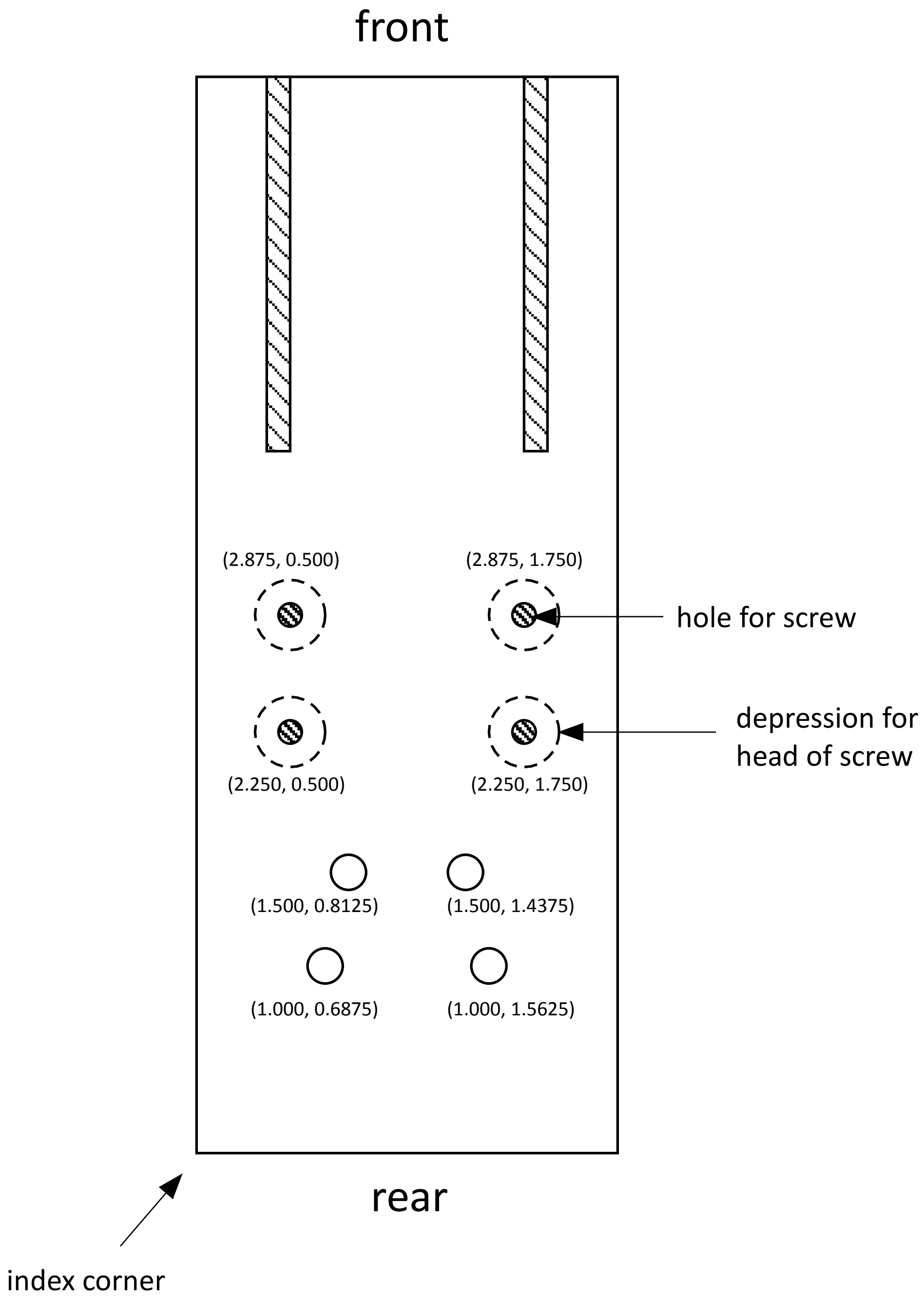


numbers in parentheses show distance (in.) from center (vertical, horizontal) to index corner

### bottom view

### Restrainer for 24-34 g mouse
(drawn to scale)

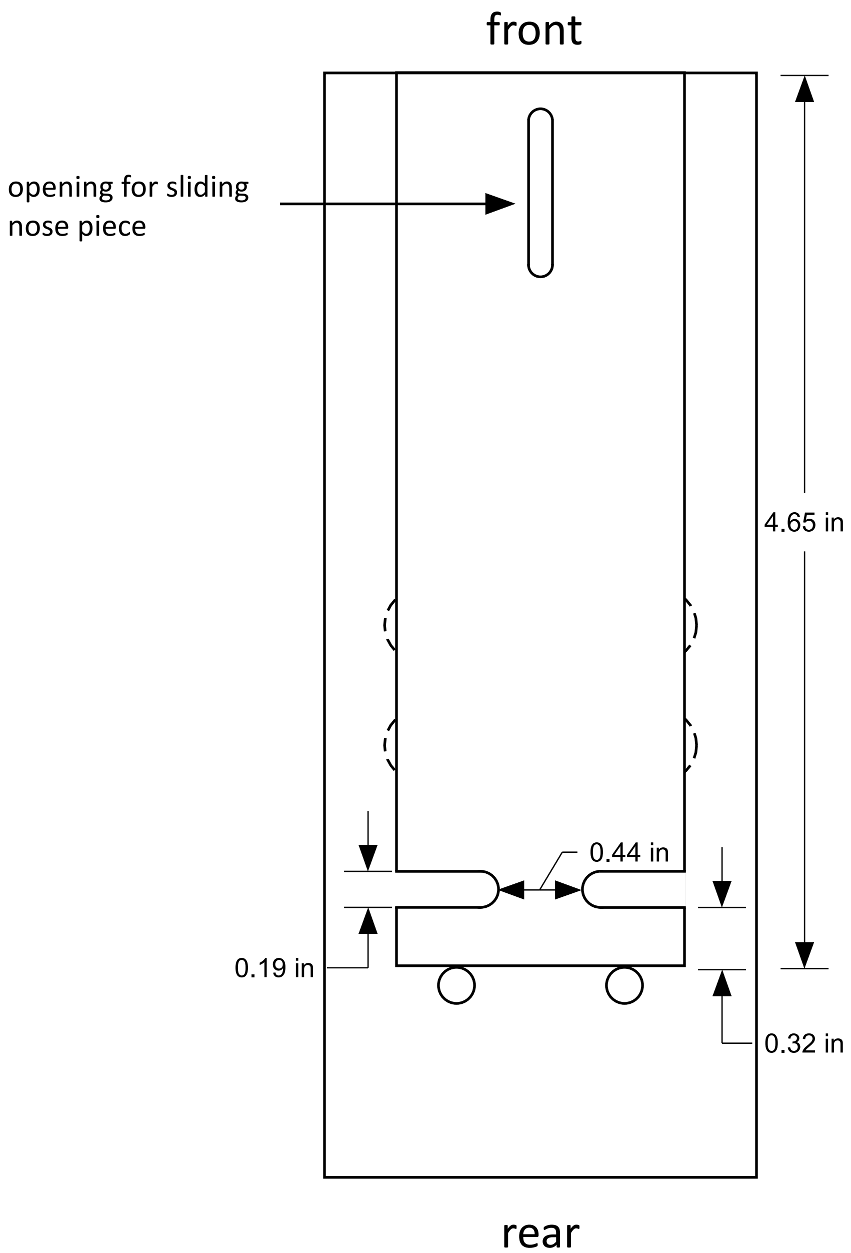


top view

### Restrainer for 24-34 g mouse
### (drawn to scale)

#### side view

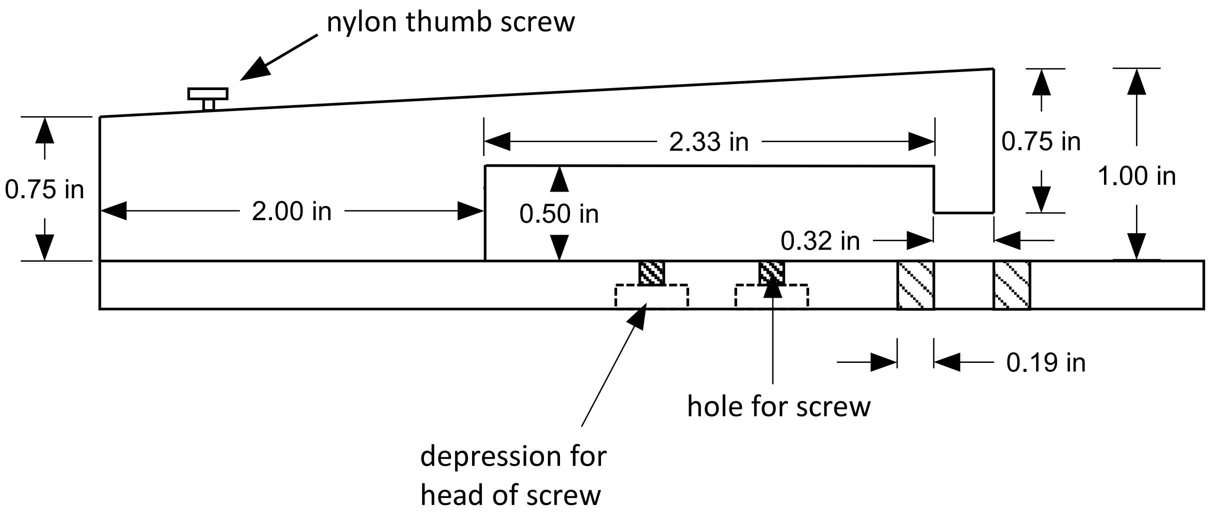


#### front view

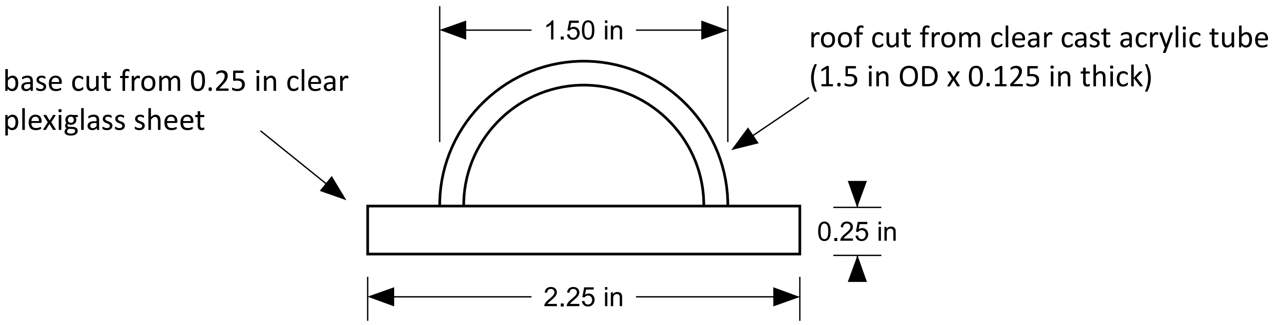

### Restrainer for 24-34 g mouse (drawn to scale)

#### nose piece

cut from acrylic rod

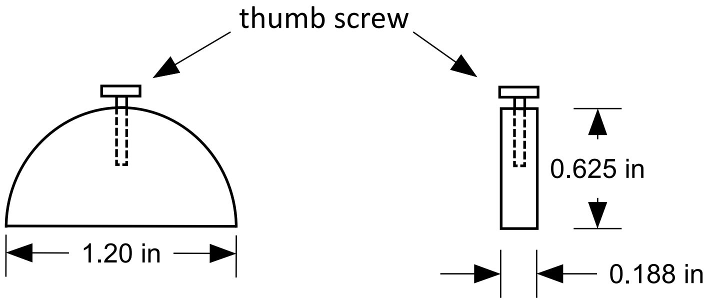


front view

side view

#### hind limb pins

cut from acrylic rod

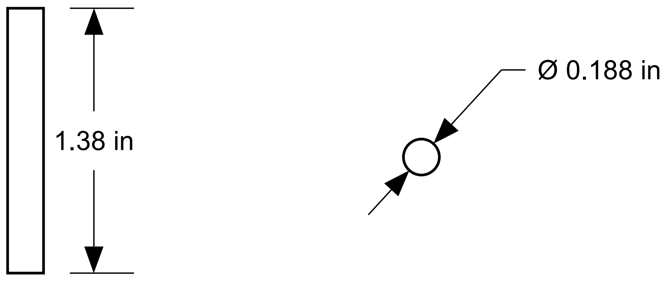


front view

top view
