## Supplementary File 4: Restrainer B blueprint for "Ablative radiotherapy improves survival in autochthonous cancer mouse models"

### Restrainer for 35-47 g mouse
### (drawn to scale)

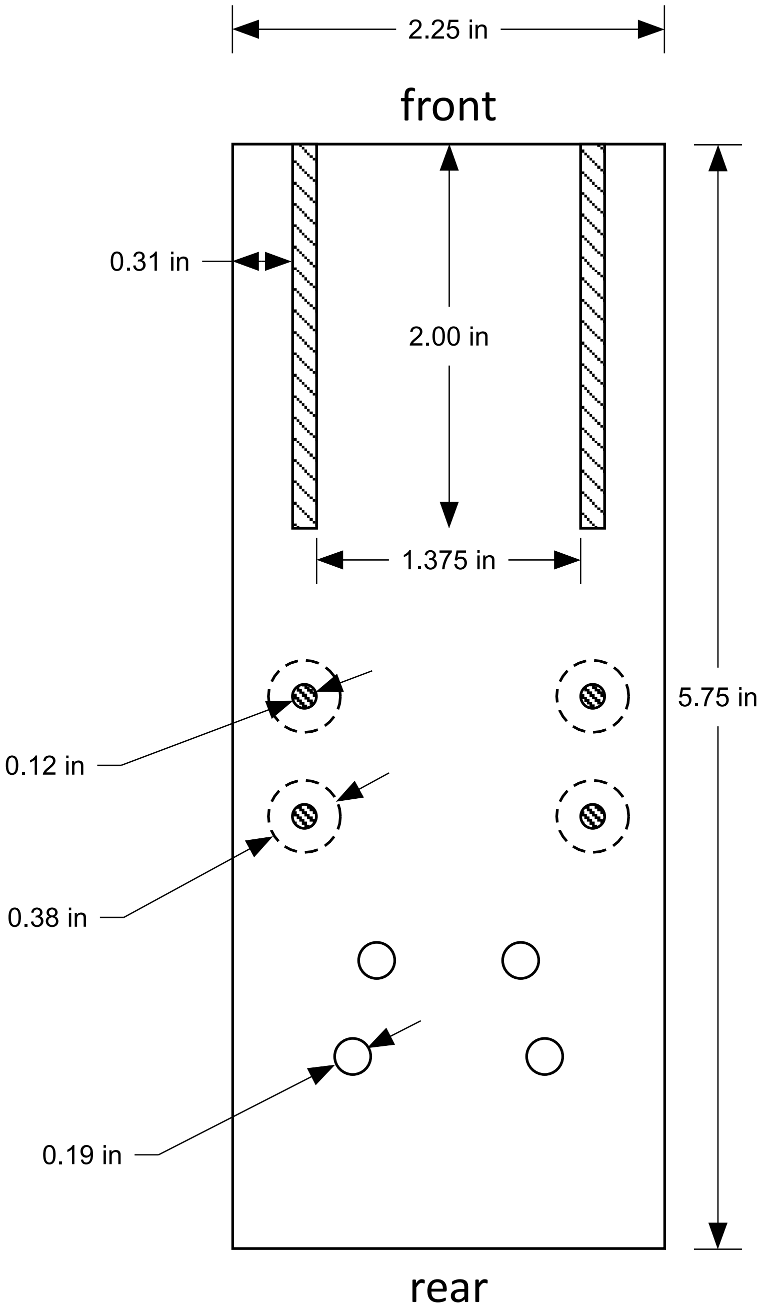


bottom view

### Restrainer for 35-47 g mouse
### (drawn to scale)

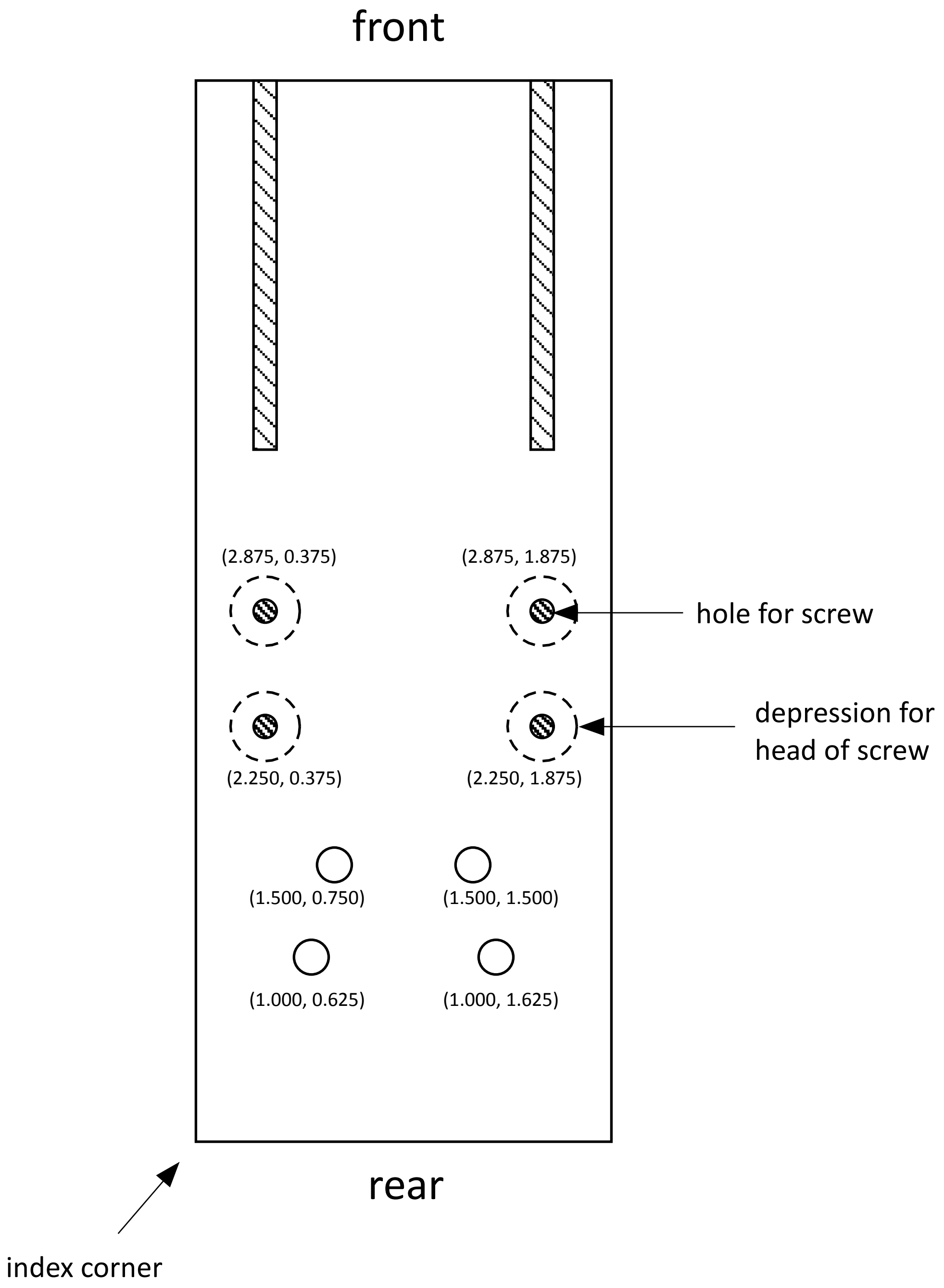


numbers in parentheses show distance (in.) from center (vertical, horizontal) to index corner

#### bottom view

### Restrainer for 35-47 g mouse
### (drawn to scale)

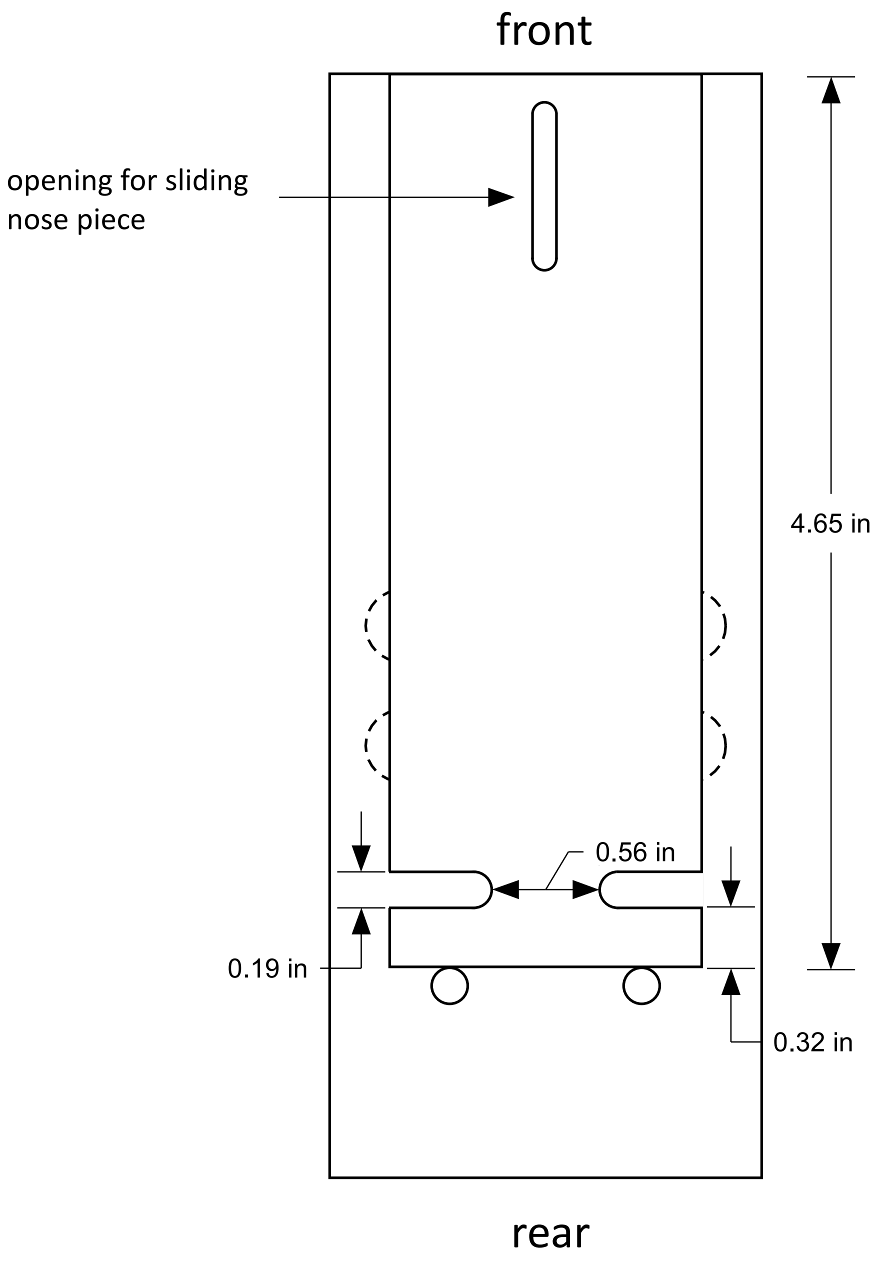


top view

### Restrainer for 35-47 g mouse
### (drawn to scale)

#### side view

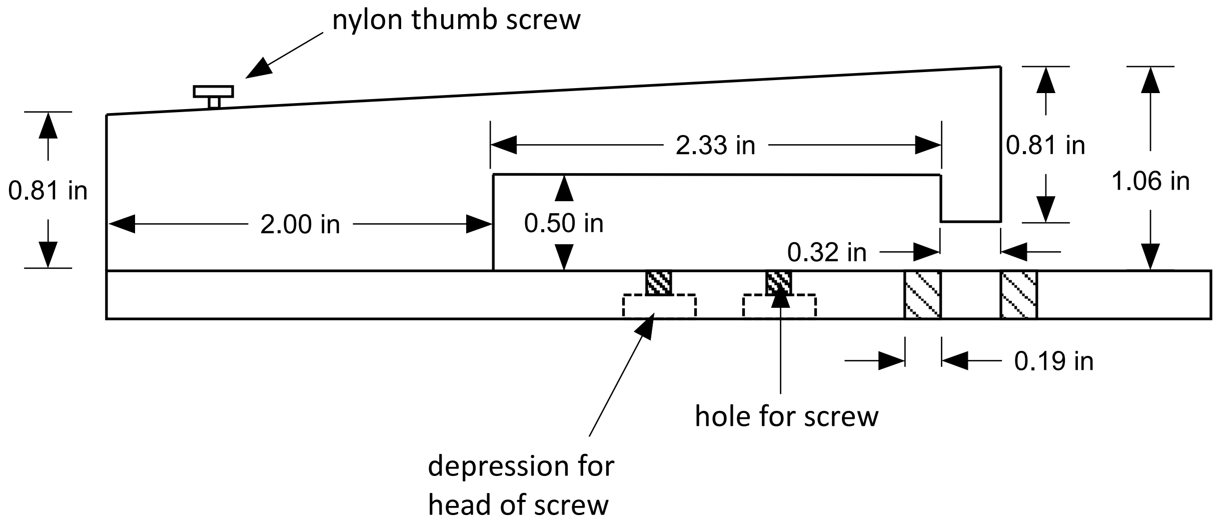


#### front view

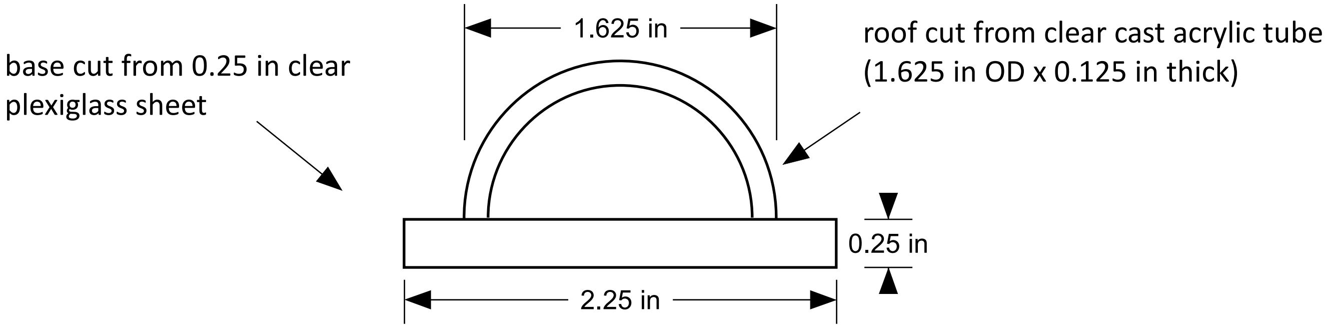

### Restrainer for 35-47 g mouse
### (drawn to scale)

#### nose piece

cut from acrylic rod

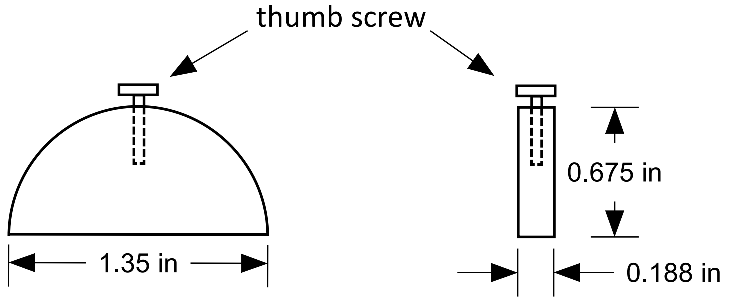


front view

side view

#### hind limb pins

cut from acrylic rod

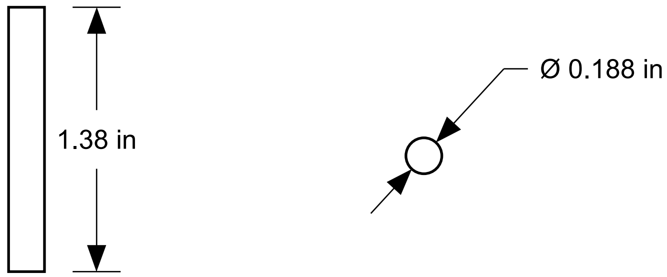


front view

top view
