## Supplementary File 6: Side blocks blueprint for "Ablative radiotherapy improves survival in autochthonous cancer mouse models"

### Universal side blocks
### (drawn to scale)

#### thorax blocks

side view

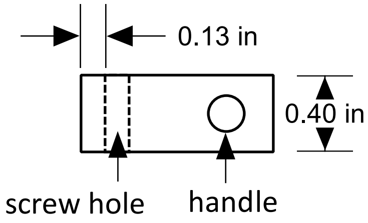


top view

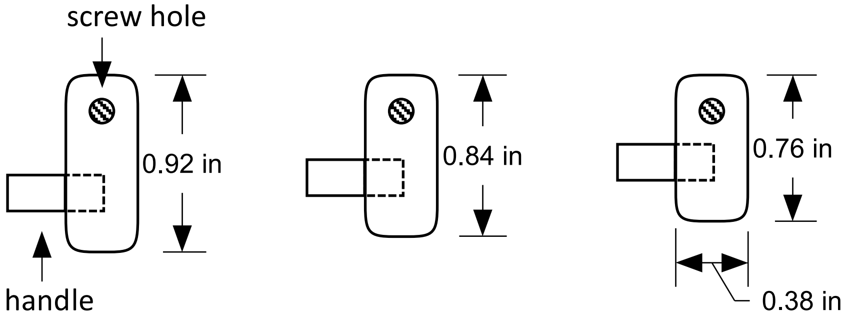


#### abdominal-pelvic blocks

side view

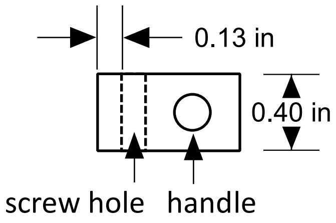


top view

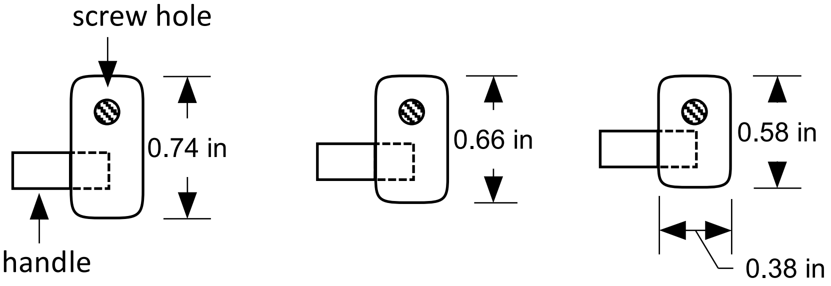
