## Supplementary figures and images for "Ablative radiotherapy improves survival in autochthonous cancer mouse models"

### Supplementary File 7: Side blocks blueprint metric

#### Universal side blocks
#### (drawn to scale)

##### thorax blocks

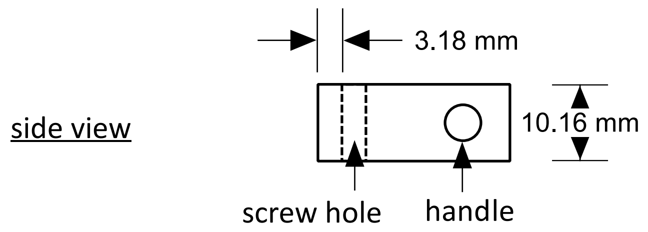


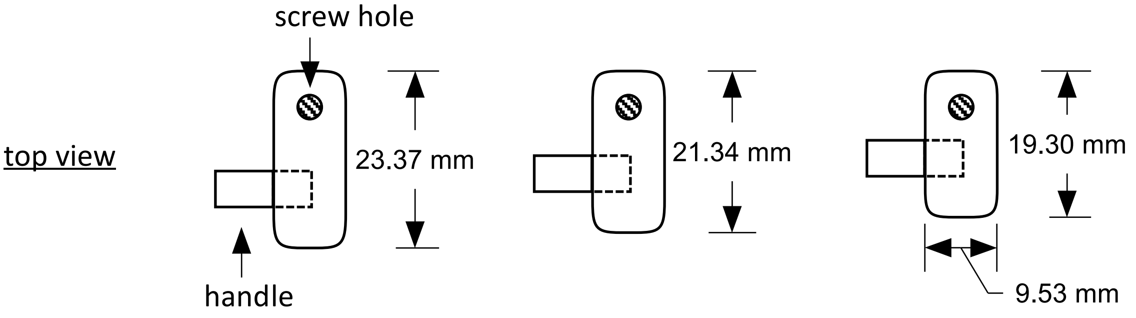


##### abdominal-pelvic blocks

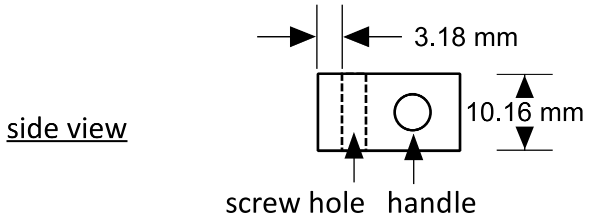


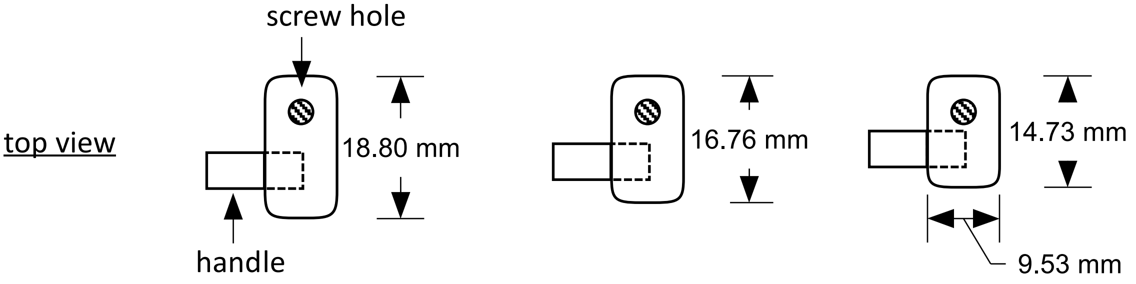
