## Supplementary File 9: Shield and collimators blueprint metric for "Ablative radiotherapy improves survival in autochthonous cancer mouse models"

### Lead shield
### (drawn at 1:2 scale)

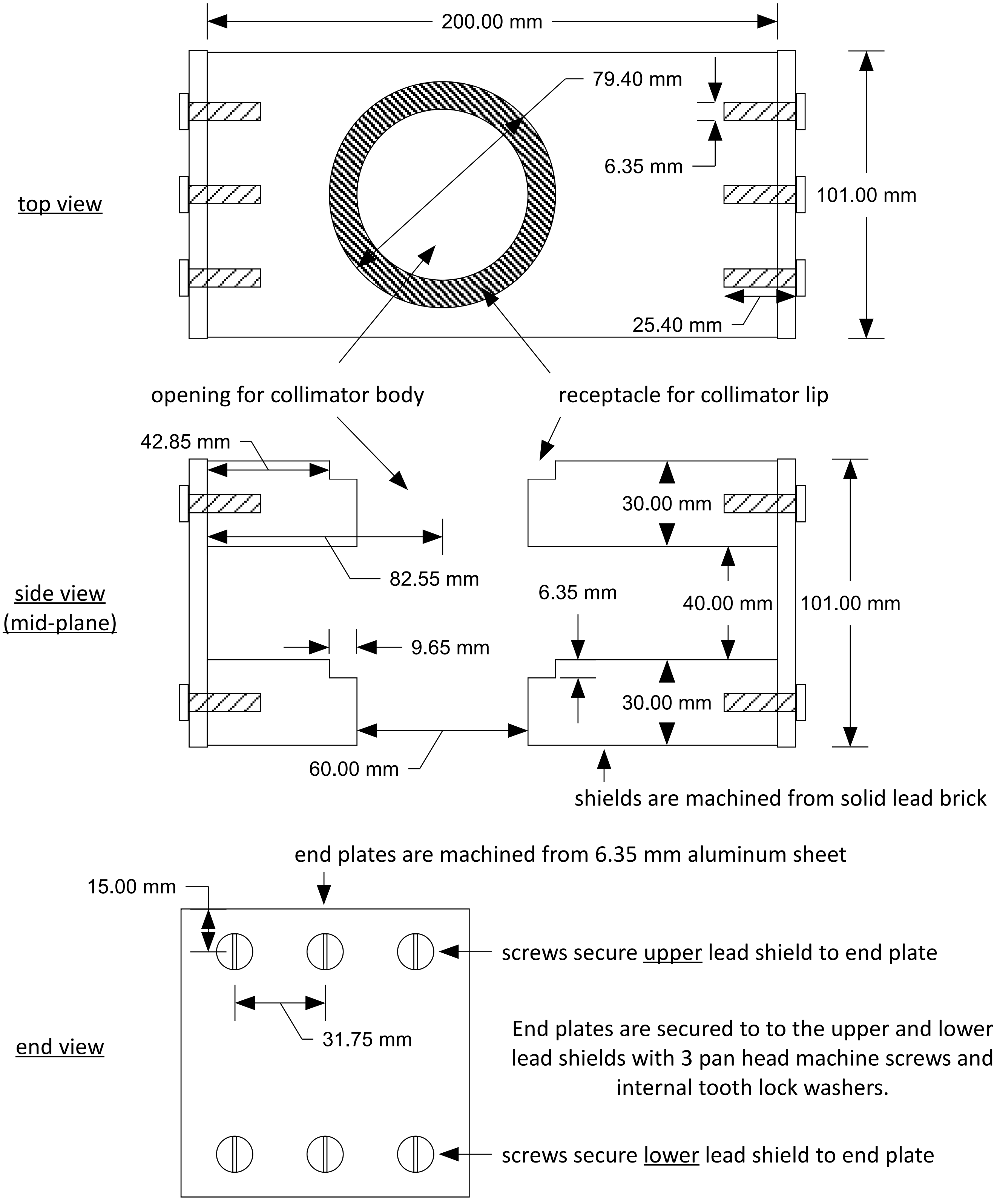

### Circular Collimators
### (drawn at 1:2 scale)

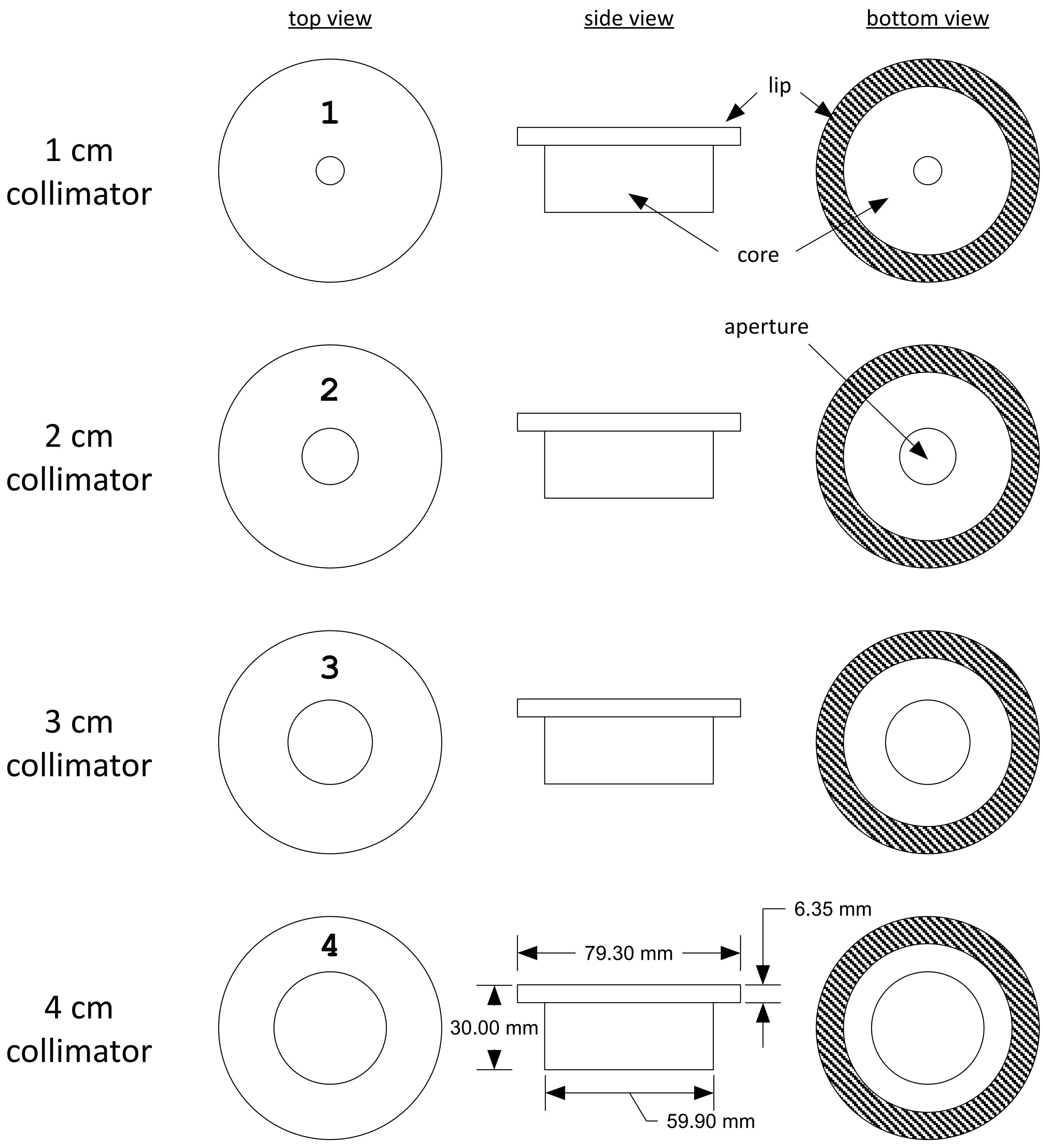


collimators are machined from solid lead bar

### base plate for restrainer
### (drawn at 1:2 scale)

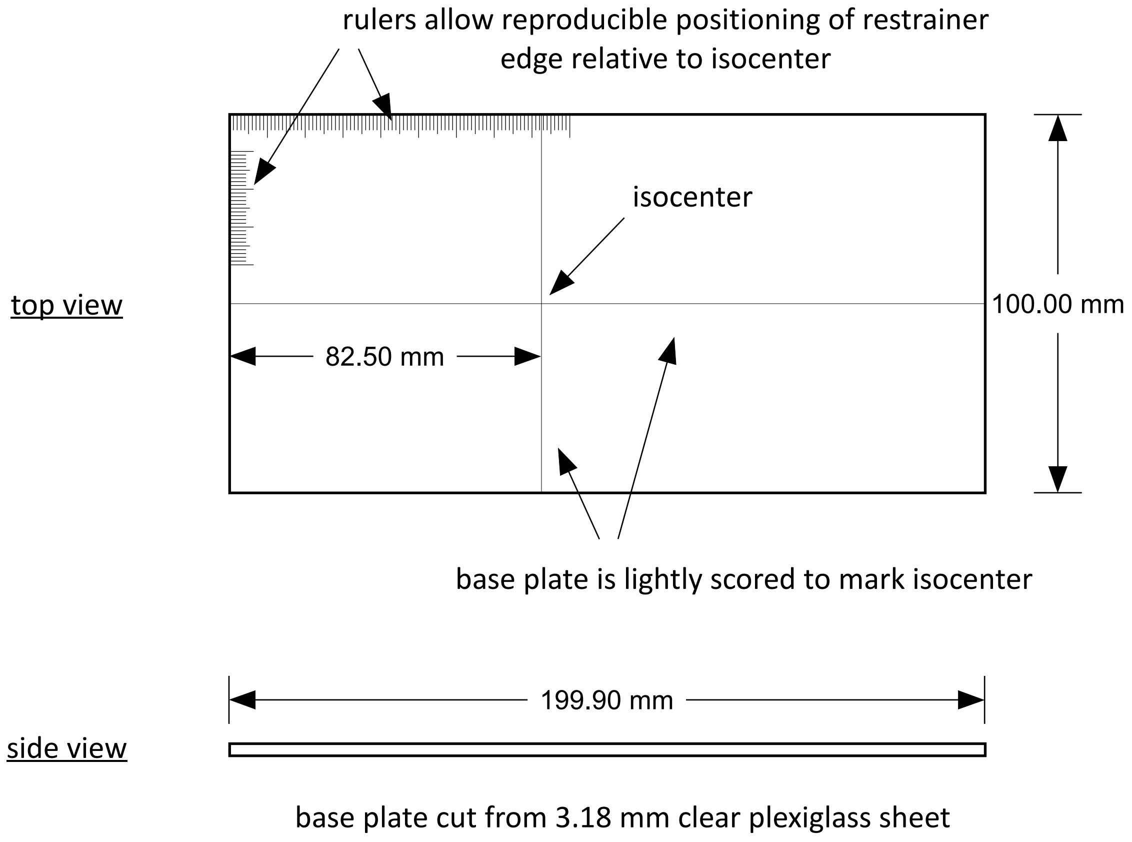

### Lead shield with circular collimator assembly and use instructions

(2 cm collimator is shown)

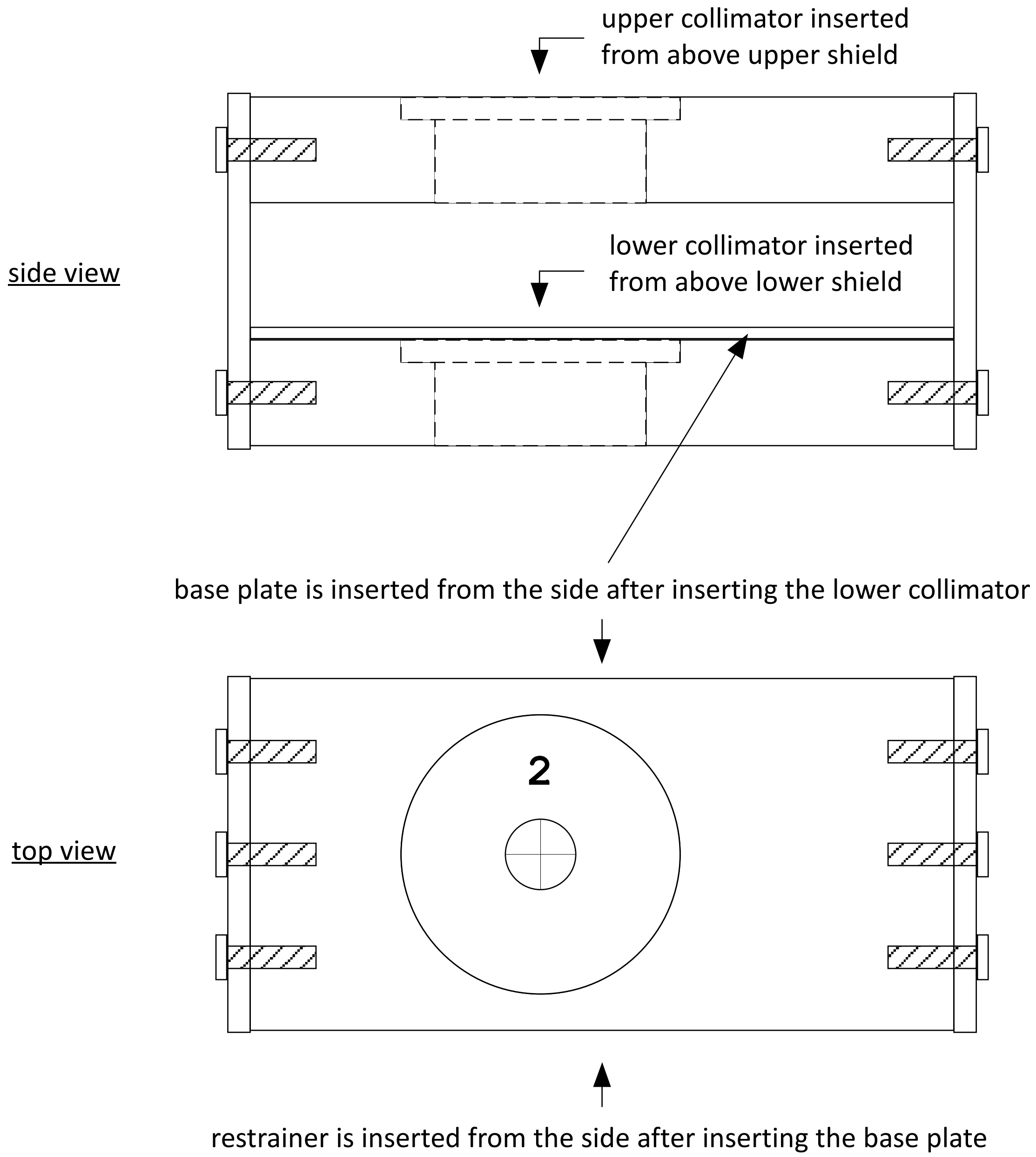

### cradle for lead shield/collimator
### (drawn at 1:2 scale)

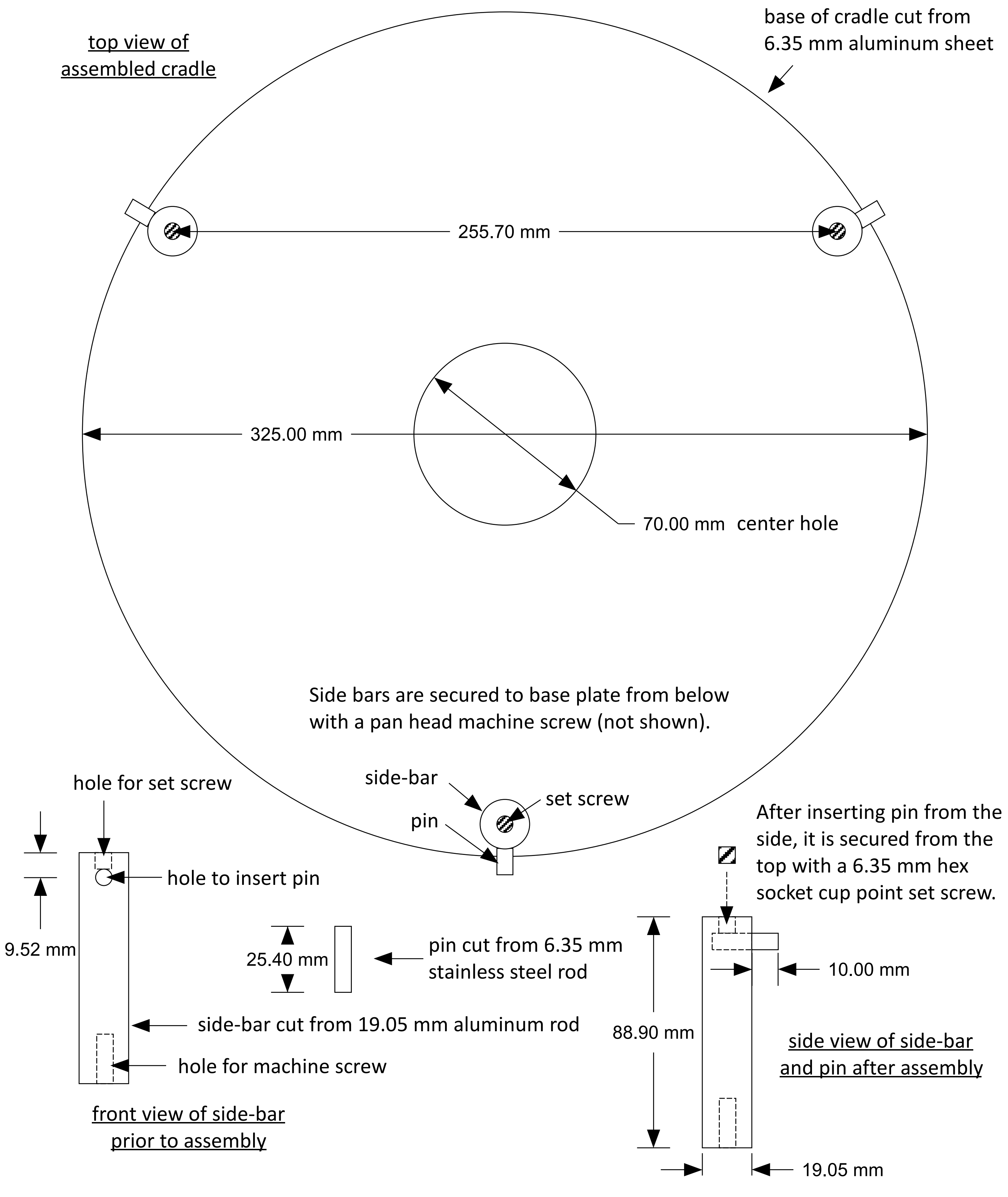

### schematic of collimated radiation field
### (drawn at 1:4 scale)

Note: the outer enclosures of the radiation chamber are not shown for clarity

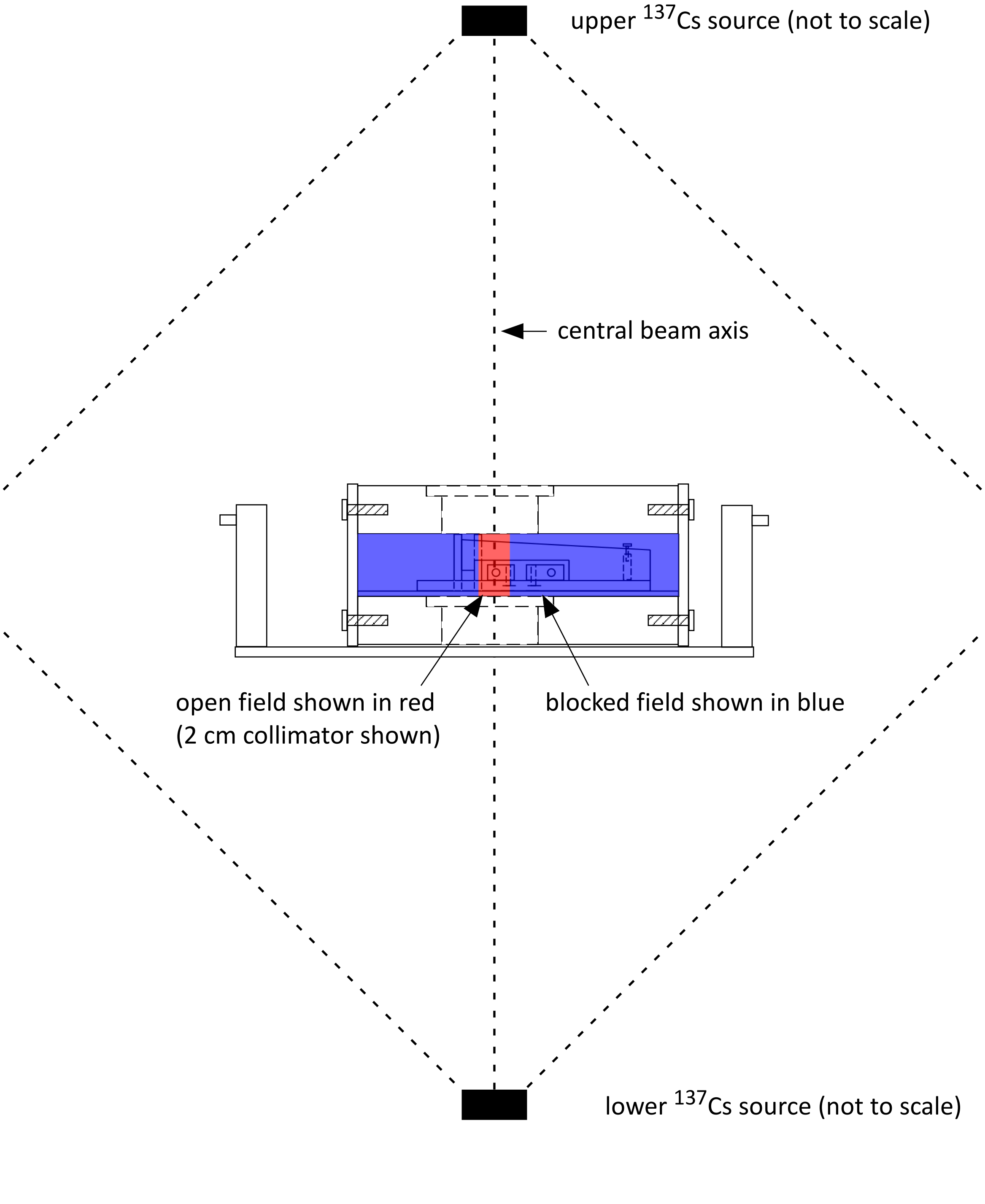
